# Foxg1 promotes neuron allocation to engrams and facilitates fear memorization

**DOI:** 10.64898/2026.02.13.705789

**Authors:** Maria Ayub, Antonello Mallamaci

## Abstract

In addition to orchestrate telencephalic development, the transcription factor Foxg1 is involved in mutual positive feedback with neuronal activity. Based on that, we hypothesized an involvement of it in learning and engram dynamics. We observed that its sparse and mild neuronal upregulation improved learning abilities as evaluated by a CFC assay. This effect was specifically pronounced upon early memory retrieval and disappeared when Foxg1-GOF neurons were silenced by a Gi DREADD effector. A prevailing positive relationship was detectable between Foxg1 expression level and the probability of a neuron getting recruited into training and recall engrams, in structures involved in both short- and long-term memory formation. Moreover, Foxg1 upregulation elicited a generalized shrinkage of engrams and increased the fraction of late recall engram cells already active at the time of training. Together, these findings establish Foxg1 as a key effector linking neuronal excitability to engram allocation and memory recall.

## INTRODUCTION

The ability of the brain to acquire, store, and retrieve information underlies the core principles of memory formation, a fundamental process essential for adaptive behavior and survival of an organism. The fact that memory persists even after the initial experience suggests that an internal representation of this acquired information is stored in the brain and can be retrieved when needed. Technological advancements in the field of cell tagging and selective single-cell manipulations have made it possible to pinpoint the storage units of information/memory called memory traces or engrams. An engram refers to a population of neurons activated by a learning experience, which are physically or chemically modified during this experience and capable of driving memory recall when reactivated either naturally or artificially^1,2,3^. Mounting evidence suggests that a single memory is supported by multiple engram ensembles, collectively called engram complex, which are distributed brain-wide and interconnected via an engram cell pathway^4^.

A defining feature of engram neurons is their elevated intrinsic excitability compared to neighboring neurons, making them likely to be recruited during learning^5,6,7^. In addition, engram neurons have increased signs of synaptic plasticity, e.g., larger spine density and more effective connectivity to other downstream engram cells^8,9,10^. These key characteristics of engram cells are regulated by molecular drivers such as transcription factors (TFs) and epigenetic modifiers. For example, upregulation of phospho-Creb1 (pCreb1) enhances neuronal excitability and neuronal allocation to engrams, whereas its reduction impairs memory recall^5^, Npas4 regulates excitatory-inhibitory neuronal balance in hippocampus and contributes to contextual discrimination^11^, KLF family members, and SRF stabilize memory associated circuits^12,13^.

The forkhead box protein G1 (Foxg1) is an evolutionary conserved transcription factor essential for development of forebrain, where it regulates neural precursor regionalization and self-renewal, as well as neuronal inhibitory/excitatory differentiation and laminar commitment^14,15,16^. Accumulating evidence indicates that Foxg1 also exerts crucial postnatal roles in cortical and hippocampal neurons. In pyramidal neurons, it regulates dendritic growth, synaptic plasticity, and excitability^17,18,19,20^. Loss of FOXG1 is associated with a variant of the synonymous syndrome, marked by intellectual disability, epilepsy, and social deficits^21,22,23^.

Given that elevated excitability is a key feature of engram neurons, we hypothesized that Foxg1 may modulate memory formation and recall, by regulating neuronal allocation into engram ensembles. To address this, we combined adeno-associated viral manipulation of Foxg1 expression via artificial micro-RNAs (miRNAs)^24^, chemogenetic control of neuronal activity, contextual fear conditioning (CFC) assays, activity-dependent genetic labeling (TRAP2)^25^ and immunoprofiling of immediate early gene (IEG)-expressing engram cells. We found that Foxg1 persistently regulates neuronal recruitment into engrams across hippocampal, amygdalar, cortical, and hypothalamic regions, also influencing engram size and short-term memory retrieval. Moreover, chemogenetic analysis revealed that Foxg1 effects on memory dynamics are partly mediated through modulation of neuronal excitability.

## RESULTS

### Foxg1 promotes neuron recruitment into short-term memory engrams and facilitates fear memory retrieval

As mentioned, the probability of a neuron being recruited into an engram is strongly influenced by its intrinsic excitability^5,26^. Since Foxg1 promotes neuronal activity and excitability^19,27^, we asked whether manipulating its expression levels within a random subset of neurons accordingly modulates their recruitment into an engram.

To achieve sparse (and gentle) modulation of neuronal Foxg1 expression levels, we employed adeno-associated viruses encoding for artificial, Foxg1-activating or -inhibiting microRNAs. These AAVs and their control also encoded for Emerald Green Fluorescent Protein, EmGFP, allowing the identification of transduced cells. To track neurons engaged in the training engram, we utilized the tamoxifen-activatable TRAP2:Ai9 reporter system^25,28^. Specifically, on post-natal day 28 (P28), TRAP2:Ai9 reporter mice received retro-orbital injections of Foxg1-modulating AAVs. Next, on P56, they were intraperitoneally injected with 100mg tamoxifen per kg of body weight and, 16-20 hours later, they underwent contextual fear conditioning (CFC), to permanently label neurons activated during memory encoding (Figure 1A,B). Interestingly, at this time all treatment groups displayed comparable freezing times, suggesting similar efficiency in encoding fear experience (Figure 1C and Supplementary Figure 1A,B). One day later, trained mice were interrogated for memory retrieval in the original A training context. Animals made sparsely gain-of-function (“GOF”) for Foxg1 exhibited longer freezing times compared to both controls (Ctrl) and sparsely loss-of-function (“LOF”) models [GOF: (168.7±12.9)s vs Ctrl: (130.4±12.6)s, with *p*<0.03, and vs LOF: (130.8±14.9)s, with *p*<0.03], indicating a positive impact of Foxg1 on memory recall (Figure 1D). One more day later, the same animals were further interrogated for memory retrieval in a novel B context, with distinct tactile and visual cues. Here, GOF mice showed an appreciable trend to outperform LOF models in contextual discrimination (*p*<0.09) (Figure 1E). This suggests that sparsely elevated Foxg1 not only strengthens memory retrieval but may also improve its precision.

**Figure 1.**
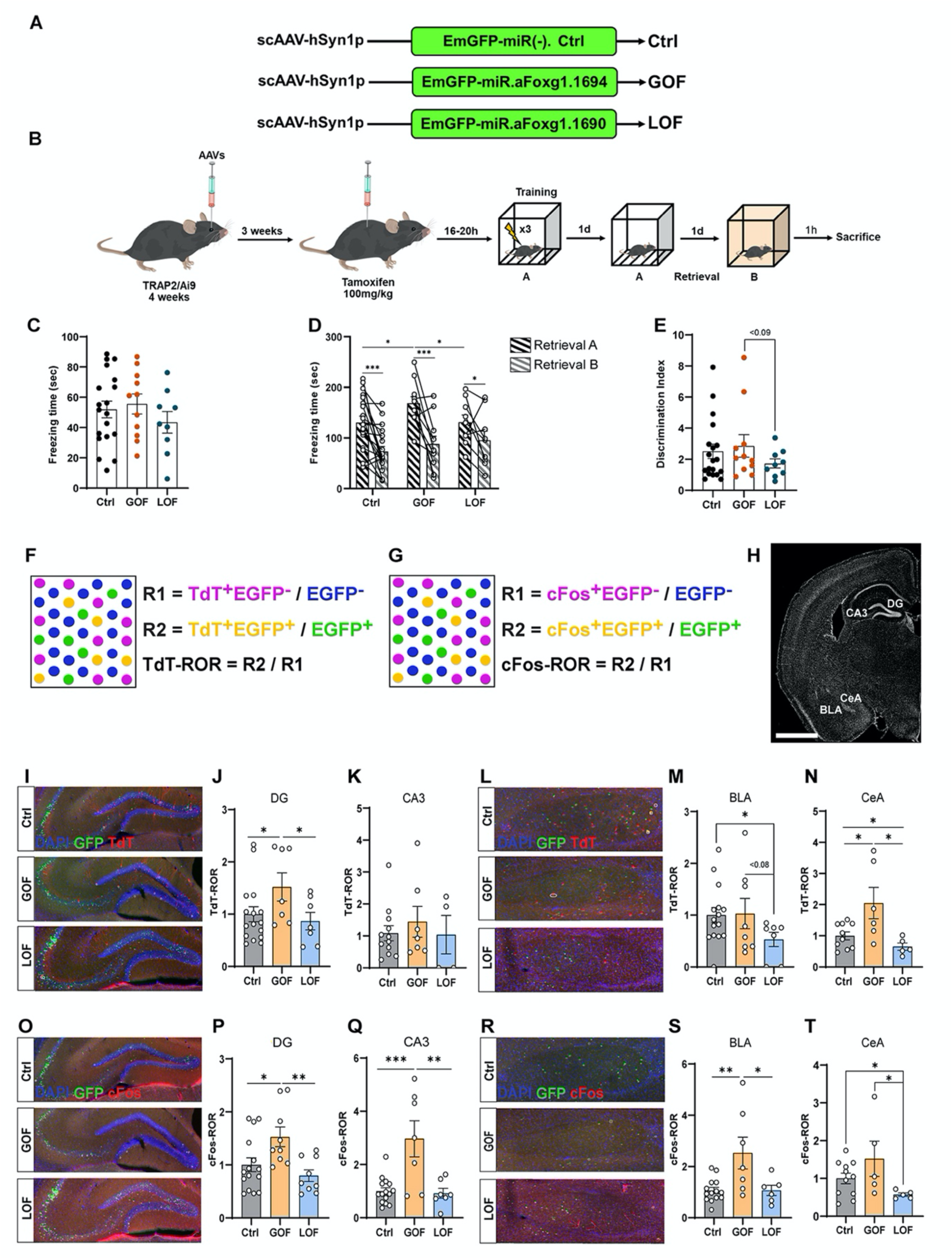
Foxg1 promotes neuron recruitment into short-term memory engrams and facilitates fear memory retrieval. **A**: Schematics of adeno associated viruses (AAVs), control (Ctrl), gain-of function (GOF), and loss of function (LOF) for Foxg1, delivered retro-orbitally into TRAP2/Ai9 mice. **B:** Experimental timeline. **C**: Distribution of total freezing times (sec) during training session of contextual fear conditioning (CFC), in Ctrl, GOF, and LOF groups. **D:** Distribution of total freezing time (sec) during retrieval sessions, in the training context (Retrieval A) and a novel context (Retrieval B). **E:** Context discrimination ratio (Retrieval A vs Retrieval B) in Ctrl, GOF, and LOF groups. **F, G:** Engram analysis procedure: quantification of Ctrl-normalized TdT-ROR (**F**) and cFos-ROR (**G**). **H:** Representative image of mouse brain coronal section, Scale bar, 1000 µm. **I-N:** Quantification of TdT-RORs in selected regions of brains treated with Ctrl, GOF, and LOF AAVs. **O-T:** Quantification of cFos-RORs in selected regions of brains treated with Ctrl, GOF, and LOF AAVs. **C, D, E:** n = 19 (Ctrl), 11 (GOF), 9 (LOF). **I-T:** n = 10-16 (Ctrl), 5-9 (GOF), 5-9 (LOF). ROR, ratio of ratios; DG, dentate gyrus; CA3, hippocampal CA3 field; BLA, basolateral amygdala; CeA, central amygdala. Error bars indicate standard error of the mean (SEM). Statistical significance of results was evaluated by t-test assay (unpaired between groups, paired within groups, one-tailed). * *p*<0.05, ** *p*< 0.01, *** *p*< 0.001.

Finally, two hours after the latter retrieval session, to quantify the impact of distinctive Foxg1 manipulations on neuron recruitment into engrams underlying training and retrieval processes, the animals were perfused and analyzed (Figure 1B).

As for the training engram, the prevalence of TRAPed cells (TdTomato^+^ cells) among transduced (EGFP^+^) and non-transduced neurons (EGFP^-^) was evaluated, and the ratio between the former and the latter (ratio of ratios, ROR) was employed as an index of this impact (Figure 1F). This analysis was primarily performed across key memory-related regions, such as the hippocampal formation, including dentate gyrus (DG) and Cornu Ammonis 3 field (CA3), and the amygdala, including basolateral (BLA) and central amygdala (CeA) (Figure 1H). Compared to controls, TdTomato-ROR (TdT-ROR) was increased in DG [+(52.01±26.87)%, *p*<0.03] and CeA [+(104.64±50.45)%, *p*<0.01] upon Foxg1 upregulation and decreased in BLA [-(47.06±13.92)%, *p*<0.03] and CeA [-(34.97±12.02)%, *p*<0.05] upon Foxg1 downregulation (Figure 1I-N), as expected. Additional memory-associated regions were further profiled. Mirroring the pattern observed in DG and amygdala, Foxg1 upregulation increased TdT-ROR in medial prefrontal cortex, including prelimbic (PL) and infralimbic (IL) cortices, as well as in the hypothalamus (Hypo), while the opposite manipulation reduced TdT-ROR in Hypo (Supplementary Figure 2B,C). Curiously, Foxg1 downregulation increased neuronal engagement in motor and somatosensory cortex (MC, SC) (Supplementary Figure 2A). These results indicate that, rather than generally promoting neuronal recruitment into the training engram, Foxg1 exerts a region-specific bidirectional effect, whereby higher Foxg1 promotes neuronal recruitment into hippocampal, amygdalar, and prefrontal ensembles, and reduces this process within sensory and motor cortices.

Concerning Foxg1 impact on neuronal engagement in the retrieval engram, the prevalence of cFos^+^ cells among EGFP^+^ and EGFP^-^ neurons was evaluated, and the corresponding ROR (cFos-ROR) was employed as an index of this impact (Figure 1G). Compared to controls, cFos-ROR was increased in DG [+(47.29±17.88)%, *p*<0.02], CA3 [+(196.60±67.80)%, *p*<10^-4^], BLA [+(152.65±61.90)%, *p*<0.002], and CeA [+(52.03±46.77)%, *p*<0.08] upon Foxg1 upregulation, and decreased in CeA [-(42.95±4.70)%, *p*<0.03] upon Foxg1 downregulation (Figure 1O-T). Interestingly, Foxg1 also positively affected cFos-ROR in PL, IL, Hypo, MC and piriform cortex (PC) (Supplementary Figure 2D-F). Altogether, these results further suggest that at the time of memory retrieval, elevated Foxg1 generally favors neuronal engagement in the engram.

### Foxg1 reduces engram size at the time of memory encoding and short-term retrieval

A key feature of memory traces is the dynamic regulation of engram size, i.e., the proportion of neurons recruited into an engram during encoding and active during recall^29,30,31^.

To test whether Foxg1 has any effect on this engram dynamics, first we analyzed engram size during memory encoding (TdTomato^+^/DAPI^+^) and retrieval (cFos^+^/DAPI^+^) across hippocampal, amygdalar, cortical, and hypothalamic regions. Foxg1 manipulations produced consistent effects on engram size. Its upregulation resulted into a shrinkage of the training engram, in DG [GOF: (0.62±0.08)% vs. Ctr: (1.40±0.23)%, *p*<0.02], CA3 [GOF: (0.24±0.06)% vs. Ctr: 0.33±0.03%, *p*<0.08], and CeA [GOF: (1.11±0.25)% vs. Ctr: (2.42±0.30)%, *p*<0.004] (Figure 2A-D), and the recall engram, in DG [GOF: (2.26±0.25)% vs. Ctr: (2.86±0.29)%, *p*<0.07] and CA3 [GOF: (0.63±0.09)% vs. Ctr: (1.27±0.26)%, *p*<0.05] (Figure 2E,F). Similar phenomena were observed in neocortex, PC, prefrontal cortices and Hypo (Supplementary Figure 3A-F) Surprisingly, a size reduction of training engram was observed in CA3 upon Foxg1 downregulation [LOF: (0.20±0.01)% vs. Ctr: (0.33±0.03)%, *p*<0.02] (Figure 2B). Considering the small fraction of AAV-transduced cells, these results indicate that the effects of Foxg1 manipulations are also largely not-cell autonomous.

**Figure 2.**
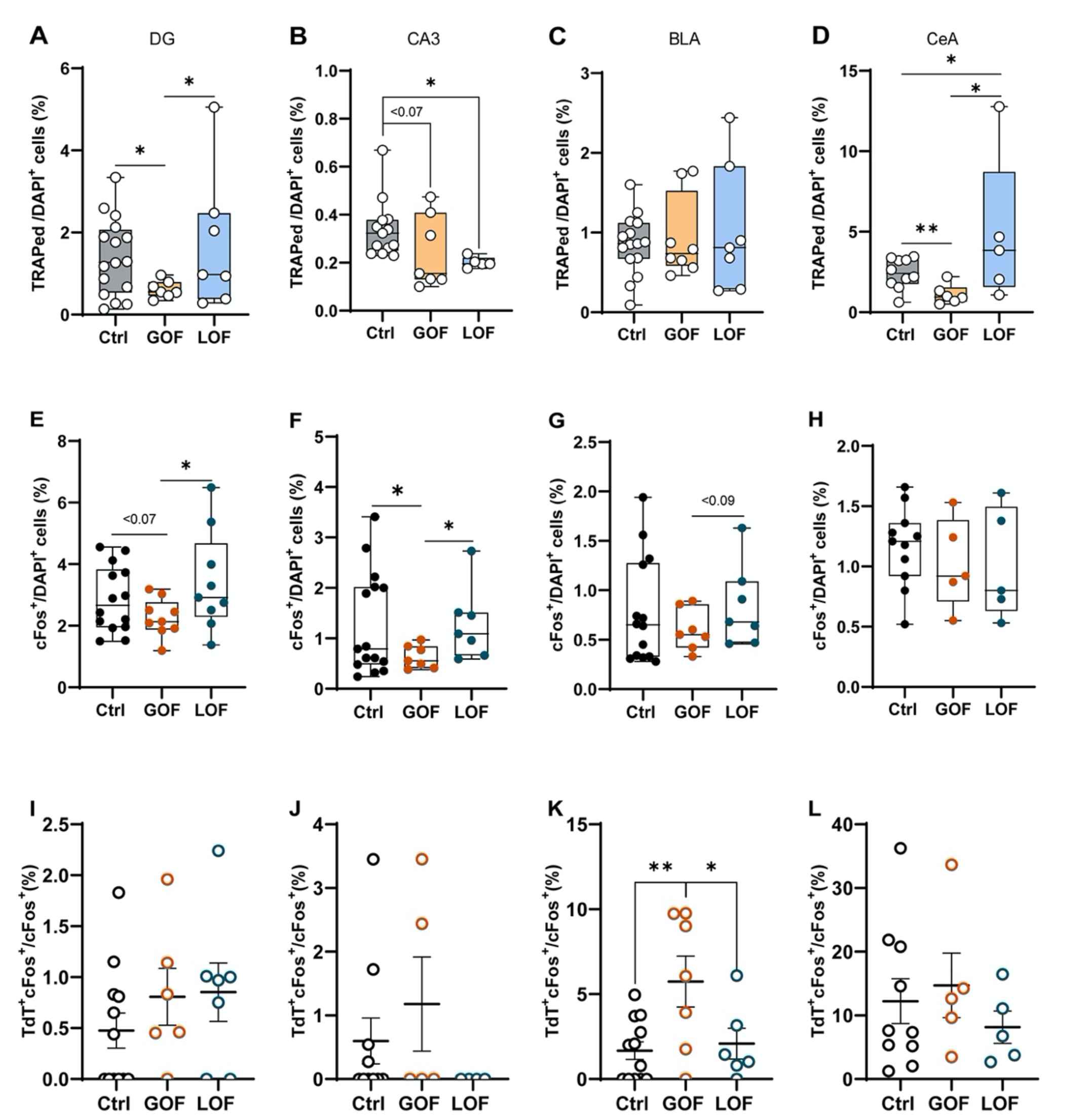
Foxg1 reduces engram size at the time of memory encoding and short-term retrieval. **A-L**: Quantification of TRAPed cells (TdTomato^+^ cells/DAPI^+^ cells, A-D), cFos^+^ cells (cFos^+^ cells/DAPI^+^ cells, E-H), and double positive cells (TdTomato^+^ cFos^+^ cells/cFos^+^ cells, I-L), all displayed as absolute percentages. n = 10-17 (Ctrl), 4-10 (GOF), 5-9 (LOF). Error bars indicate standard error of the mean (SEM). Statistical significance of results was evaluated by t-test assay (unpaired, one-tailed). * *p*<0.05, ** *p*<0.01. DG, dentate gyrus; CA3, hippocampal CA3 field; BLA, basolateral amygdala; CeA, central amygdala.

Next, we evaluated the fraction of recall engram neurons that were also part of the training engram (TdTomato^+^Fos^+^/Fos^+^). We found that this parameter was positively correlated to Foxg1 expression level, in most of the regions analyzed (CA3, BLA, CeA, MC, SC, AC, PC, PL, IL, and Hypo), displaying statistically significant changes in same cases (Figure 2I-L and Supplementary Figure 3G-I).

In synthesis, the engram downsizing and stabilizing effects elicited by Foxg1 upregulation apply to both training and retrieval engrams.

### Foxg1 promotes neuron engagement in late memory engrams

Long-term memory recall requires the persistence of engram ensembles across time^32–35^. Previous work has shown that transcriptional regulators such as CREB, NFAT, and NPAS4 contribute to the stabilization of memory traces during remote recall^36,5,11^. However, whether Foxg1 manipulations exert persisting effects on long-term memory recall and engram allocation remains elusive. To test this, AAV-treated wild-type mice were trained in CFC training chamber on P49 and tested for recall in the same training context on P56 (Figure 3A,B). All groups displayed robust memory retention, indicated by strong freezing behavior during recall. Although differences did not reach statistical significance, GOF mice exhibited the longest freezing time, (173.93±18.45)s, followed by the Ctrl group with (168.76±24.16)s, whereas LOF mice showed the shortest, (147.34±27.72)s (Figure 3C). Consistent differences were also detectable in number of freezing episodes and freezing latency (Supplementary Figure 4A,B).

**Figure 3:**
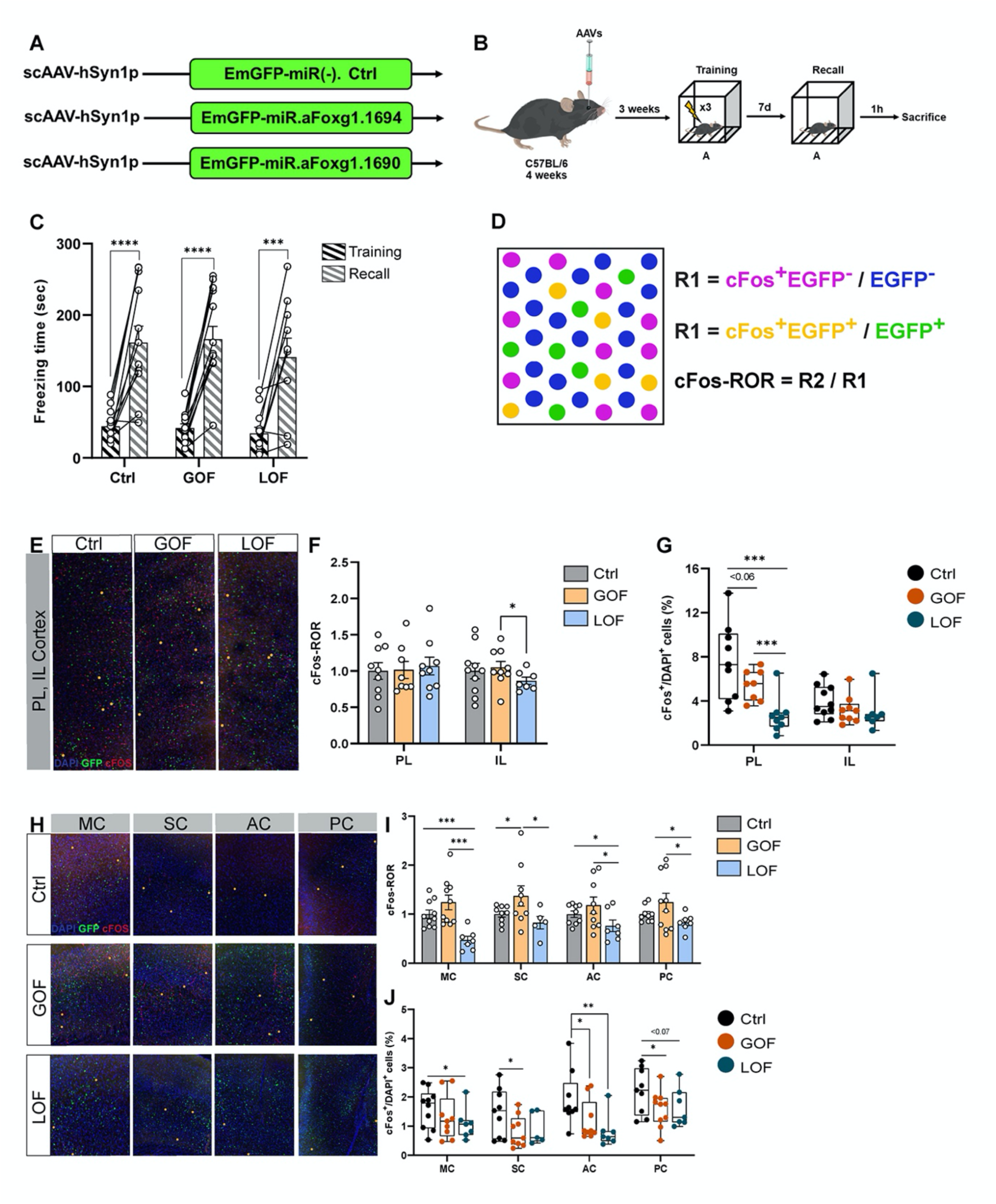
Foxg1 promotes neuron engagement in late memory engrams. **A**: Schematics of adeno associated viruses (AAVs), control (Ctrl), gain-of function (GOF), and loss of function (LOF) for Foxg1, delivered retro-orbitally into C57Bl/6 mice. **B:** Experimental timeline. **C**: Distribution of total freezing times (sec) during training and late recall sessions of contextual fear conditioning (CFC), in Ctrl, GOF, and LOF groups. **D:** Quantification analysis procedure. **E,H:** Representative immunofluorescence images employed for this analysis. **F,G,I,J**: Quantification of Ctrl-normalized cFos-ROR (F,I) and absolute cFos^+^/DAPI^+^ cell ratio (G,J) in selected regions of brains treated with Ctrl, GOF, and LOF AAVs. **C:** n = 11 (Ctrl), 12 (GOF), 9 (LOF). **F,G,I,J**: n = 9-11 (Ctrl), 8-11 (GOF), 5-9 ( LOF). Error bars indicate standard error of the mean (SEM). Statistical significance of results was evaluated by t-test assay (unpaired between groups, paired within groups, one-tailed). * *p*<0.05, ** *p*<0.01, *** *p*<0.001, ROR, ratio of ratios; PL, pre-limbic cortex; IL, infra-limbic cortex; MC, motor cortex; SC, somatosensory cortex; AC, auditory cortex; PC, piriform cortex.

To assess whether Foxg1 influences underlying neuron allocation to engrams, we quantified cFos-ROR in cortical regions, preferentially involved in long-term memory formation. Here, “GOF mice” showed higher cFos-ROR compared to the “LOF group”, with pronounced differences in IL [GOF: (104.83±8.48)% vs. LOF: (86.53±5.04)%, *p*<0.05)], MC [GOF: (124.05±15.06)% vs LOF: (47.09±7.89)%, *p*<0.001], SC [GOF: (137.17±20.74)% vs LOF: (82.63±12.75)%, *p*<0.05], AC [GOF: (118.25±16.96)% vs LOF: (76.15±11.86)%, *p*<0.04], and PC [GOF: (124.20±19.53)% vs LOF: (83.13±6.62)%, *p*<0.05]. Ctr cFos-RORs generally were between GOF and LOF values (Figure 3E,F,H,I). Consistently, Foxg1 down-regulation reduced the recall engram cFos-ROR in CA3, BLA, CeA and Hypo. Curiously, a reduction of this index was also achieved upon Foxg1 upregulation in Hypo (Supplementary Figure 4C-E).

We next examined whether Foxg1 manipulations altered overall engram size, quantified as the percentage of cFos^+^ cells relative to total (DAPI^+^) cells. Compared to Ctrl, even at this stage, Foxg1 upregulation led to a shrinkage of engram in all analyzed regions except for Hypo. Differently from what we saw at short-term memory formation, a reduced engram size was also generally observed upon Foxg1 down-regulation (Figure 3G,J and Supplementary Figure 4F-H).

Together, these results demonstrated that Foxg1 continues to regulate engram dynamics even during long-term recall, i.e., 7 days after initial training, however according to a non-monotonic pattern. Specifically, GOF enhanced preferential recruitment of neurons while reducing engram size, whereas LOF diminished both parameters.

### Foxg1 regulates memory retrieval partly through activity-dependent mechanisms

Previous work from our laboratory demonstrated that increasing Foxg1 expression in primary neocortical neurons enhances their electrical activity, whereas reducing Foxg1 diminishes it^19^. Given that a key feature of engram neurons is their higher intrinsic excitability, we hypothesized that Foxg1 may regulate engram allocation and memory formation through activity-dependent mechanisms. To answer this question, we tested whether suppressing neuronal activity by inhibitory (hM4Di, Gi) Designer Receptors Exclusively Activated by Designer Drugs (DREADDs) could counteract the effects of Foxg1 gain-of-function (GOF) manipulation on memory retrieval. The experimental session included five groups: control without DREADDs (Ctrl), control with inhibitory DREADD (Ctrl-Gi), Foxg1 GOF with Gi (GOF-Gi), and, as specificity controls, control with excitatory DREADD (Ctrl-Gq), Foxg1 LOF with Gq (LOF-Gq) (Figure 4A). All mice were trained in the CFC chamber and, the following day, tested for memory recall in the same context (Figure 4B). Fifteen to thirty minutes prior to recall, Deschloroclozapine (DCZ) was administered to either inhibit (Gi) or activate (Gq) transduced neurons. Memory recall was quantified using three parameters, i.e., total freezing time, number of freezing episodes, and freezing latency.

**Figure 4.**
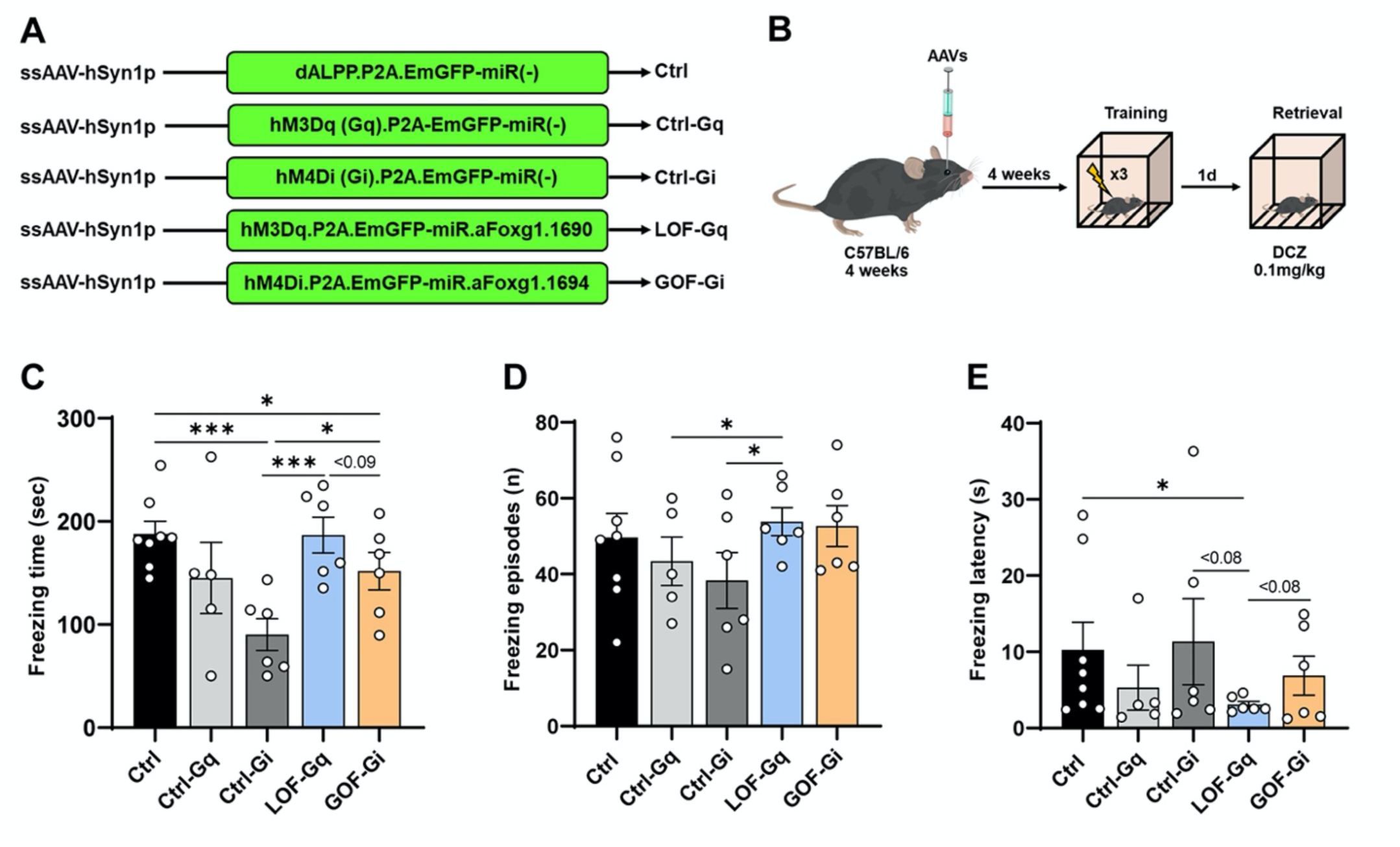
Foxg1 regulates memory retrieval partly through activity-dependent mechanisms. **A**: Schematics of adeno-associated viruses (AAVs), control (Ctrl), Foxg1-Ctrl coupled with excitatory DREADD (Ctrl-Gq), Foxg1-Ctrl coupled with inhibitory DREADD (Ctrl-Gi), Foxg1 LOF with excitatory DREADD (LOF-Gq), and Foxg1-GOF with inhibitory DREADD (GOF-Gi), delivered retro-orbitally into C57Bl/6 mice. **B**: Experimental timeline. **C-E**: Distribution of total freezing time, freezing episodes, and freezing latency in Ctrl, Ctrl-Gq, Ctrl-Gi, LOF-Gq, and GOF-Gi AAVs treated mice, during memory retrieval in the training context. n = 8 (Ctrl), 5 (Ctrl-Gq), 6 (Ctrl-Gi), 6 (LOF Gq), and 6 (GOF-Gi). Error bars indicate the standard error of the mean (SEM). Statistical significance of results was evaluated by t-test assay (unpaired, one-tailed). * *p*<0.05, ** *p*<0.01, *** *p*<0.001.

Ctrl-Gi mice displayed impaired memory recall, with the lowest total freezing and fewer freezing episodes, indicating that suppressing neuronal activity impairs memory expression^37,38^ (LOF-Gq mice conversely exhibited freezing behavior comparable to Ctrl). GOF-Gi mice also displayed reduced freezing relative to Ctrl (Figure 4C), further indicating that suppressing activity can counteract the effects of elevated Foxg1 (Figure 1D). Consistently, changes in freezing episodes and latency evoked by Foxg1-GOF alone (Supplementary Figure 1) were also abolished upon GOF-Gi manipulation (Figure 4F).

Together, these results point to a contribution of activity-dependent mechanisms to Foxg1 regulation of memory recall.

## DISCUSSION

Our study identifies the transcription factor Foxg1 as a critical regulator of neuronal engram allocation, memory retrieval, and, possibly, context discrimination. To address these aspects, we employed a combination of approaches including TRAP2-based genetic labeling, viral vector-mediated Foxg1 modulation, and chemogenetic control of neuronal activity. We observed that sparse and mild Foxg1 upregulation improved learning abilities as evaluated by a CFC assay (Figure 1). This effect was pronounced when 1 day elapsed between conditioning and retrieval sessions and lose statistical significance 7 days after conditioning (Figures 1,3). Interestingly, this effect also disappeared when neurons gain-of-function for Foxg1 were silenced by a Gi DREADD effector (Figure 4). Analysis of TRAPed and cFos^+^ cells displayed an overt positive relationship between Foxg1 expression level and the probability of a neuron getting recruited into training, retrieval and recall engrams. This phenomenon was detectable in DG, CA3, BLA and CeA, regions mainly involved in CFC. Here, it was observable at time of training and early retrieval upon Foxg1 upregulation, and at the time of delayed recall upon the opposite manipulation (Figure 1 and Supplementary Figure 4). Moreover, a similar phenomenon was observed at the time of delayed recall in MC, SC, AC and PC, brain structures reported to be involved in late memory retrieval (Figure 3). Interestingly, Foxg1 upregulation also elicited a generalized shrinkage of engrams, at time of training, early retrieval and delayed recall (Figures 2,3). This shrinkage might account for better context discrimination displayed by Foxg1-GOF mice at time of early retrieval (Figure 1). Lastly, Foxg1 upregulation also increased the fraction of recall engram cells already active at the time of training (Figure 2 and Supplementary Figure 3).

### Foxg1 promotes neuron allocation to engrams in an activity-dependent way

Engram neurons exhibit elevated excitability and enhanced plasticity that bias their allocation during learning and recall^5,38,39^. Prior studies have identified TFs such as CREB^5^, NPAS4^11^, and SRF^40^ as positive regulators of this process. Our work extends this framework by positioning Foxg1 among TFs involved in engram allocation and memory expression.

To note, artificial modulation of Foxg1 leads to anticorrelated fluctuations of Npas4 and Srf (Ref^19^ and our unpublished results). This suggests that Foxg1 impact on neuronal engram allocation is not mediated by Npas4 and Srf. On the contrary, pCreb1 protein is upregulated upon Foxg1 overexpression and its functional shut down by a dominant-negative effector decreases Foxg1-mRNA^20^. This conversely suggests that a mutual positive feedback between Foxg1 and pCreb1 may favor this allocation.

These dynamics were observed to be region-, recall time- and Foxg1 expression level-specific. In particular, a positive relationship between Foxg1 levels and neuronal recruitability into engrams could be observed in hippocampus and amygdala at the time of training and short-term memory retrieval (Figure 1 and Supplementary Figure 4), and in cortical regions upon late memory recall (Figure 3 and Supplementary Figure 2). Depending on cases, this effect was detectable upon Foxg1 upregulation, downregulation or both (Figures 1 and 3). These findings resonate with recent work suggesting that neuronal engram allocation is circuit-specific and dynamically regulated^4,41^. In this context, the ability of Foxg1 to bias neuronal allocation over time suggests it may act as a key gatekeeper of engram selectivity.

Region/time-specific transcription cofactors and epigenetic programs are likely to underlie differential effects elicited by Foxg1 on neuronal recruitment into the engram. Future studies focusing on these cofactors and programs will be needed to clarify how this takes place.

Our chemogenetic experiments also showed that Foxg1 may drive memory dynamics by promoting neuronal activity. In fact, suppressing this activity reversed the enhanced memory performance elicited upon Foxg1 upregulation (Figures 1 and 4). Previous work from our lab reported that Foxg1 enhances neuronal activity both in vitro and in vivo^19,27^. Instrumental in these phenomena can be changes in expression levels of genes encoding ion channels, receptors, plasticity and dendritic development effectors, dependent on Foxg1^18–20^. Their relative contribution to Foxg1-driven memory dynamics waits to be investigated.

### Activity-dependent promotion of Foxg1 might stabilize engrams

Previous in vitro studies from our lab showed that artificial neuronal activation results in upregulation of Foxg1 mRNA and protein levels^19,24^. To validate these findings in vivo, we administered the established proconvulsant kainic acid^42,43^ (KA) to juvenile wild-type mice and we evaluated these levels in neocortex and hippocampal formation. As expected, we found a timed and transient upregulation of both Foxg1 mRNA and protein (Supplementary Figure 5A-J). Remarkably, in the same structures of mock-injected, control mice, Foxg1 protein expression was enriched in cFos⁺ neurons compared to cFos⁻ cells (Supplementary Figure 5K), pointing to a physiological regulatory role of neuronal activity in promoting Foxg1 expression. This basal enrichment in active neurons and activity-dependent induction, suggest that Foxg1 participates in a positive feedback loop: neurons with higher Foxg1 levels are more excitable and more likely to be reactivated during learning, leading to further Foxg1 upregulation and stabilization of engram identity (Figure 2J,K and Supplementary Figure 3G-I). Such feedback loops have been proposed for pCreb1 and other transcriptional regulators that integrate excitability and plasticity to define memory allocation^41,44^. In this context, the timed non-monotonic progression of Foxg1 products evoked by neuronal hyperactivity^19^ could have evolved for proper dynamical replacement of a subset of early engram neurons with new ones.

### Foxg1 tunes cumulative engram size

An intriguing aspect of the phenotype evoked by sparse Foxg1 manipulation was the modulation of the total frequency of neurons belonging to the engram. In general, Foxg1 upregulation reduced this prevalence, regardless of the structure taken into account and the time of the evaluation, whereas Foxg1 down-regulation gave rise to a more variegated scenario (Figure 2 and Supplementary Figure 3). These phenomena likely originated from the not-cell autonomous effects exerted by manipulated cells on excitability of surrounding ones, including glutamatergic stimulation and GABAergic inhibition. In this respect, the variable outcome of Foxg1 manipulation on engram size could reflect the relative weights of these two processes. Moreover, until now the capability of Foxg1 to promote neuronal activity has been documented in bulk neuronal populations, likely reflecting its actual impact on largely prevailing glutamatergic neurons. An evaluation of Foxg1 activity specifically restricted to GABAergic cells will be needed to further clarify this point.

### Limitations and future directions

First, our study primarily relied on CFC; whether Foxg1 exerts similar effects in other behavioral paradigms, such as hippocampal-dependent spatial navigation or amygdala-driven emotional memories remains to be determined. Second, while our chemogenetic manipulations support an activity-dependent mechanism, the precise downstream effectors of Foxg1 (e.g., ion channel genes, synaptic proteins, or epigenetic regulators) require direct investigation. Finally, the region-specific effects observed suggest that Foxg1 may interact with local transcriptional landscapes or network states, an avenue for future single-cell and circuit-level studies.

## Conclusion

In summary, we identified Foxg1 as a key transcriptional regulator of neuron engram allocation and memory recall. As such, Foxg1 provides a mechanistic bridge between transcriptional control and memory dynamics. This expands our understanding of the molecular programs governing memory traces. It further suggests a special caution in precision treatment of neuropathogenic Foxg1 haploinsufficiencies and highlights Foxg1 as a potential therapeutic target for disorders of memory and cognition.

## MATERIALS AND METHODS

### Mouse genetics

Homozygous *Fos^2A-iCreER/2A-iCreER^*(TRAP2^+/+^) mice on C57Bl/6N genetic background were bought from Jackson Labs and were bred and raised on site. TRAP2^+/+^ male mice were then mated with homozygous *B6.Cg-Gt(ROSA)26Sor^tm9(CAG-tdTomato)Hze^* (Ai9^+/+^) females, kept on C57Bl/6J genetic background, generated starting from the corresponding pairs of founders available to us (a kind gift by prof. Paul Heppenstall, SISSA, Trieste). TRAP2^+/−^;Ai9^+/−^ mice were group-housed on a 12h light/dark cycle with food and water ad libitum. All experiments took place during the light phase and included mice from both sexes. Each experimental treatment included mice derived from at least 2 separate litters, each providing an identical or closely similar number of animals to each experimental group. At postnatal day 21 TRAP2^+/−^;Ai9^+/−^ mice were weaned from the breading cages and group-housed with same sex-littermates in standard mouse housing cages (3-5 mice per cage) till postnatal day 30. These animals were specifically employed for short-term fear conditioning assays.

C57BL/6 wild-type mice, males and females, were conversely used for long-term memory and chemogenetics experiments. They underwent same maintenance, light/dark parameters and experimental grouping procedures mentioned above.

CD1 wild-type mice, males and females, were used for KA experiments and underwent same maintenance and light/dark parameters mentioned above. On postnatal day 30, they were treated with KA for different time periods and were then subjected to sacrifice for further analysis.

### Drugs

#### Tamoxifen

Tamoxifen (Sigma, T5648) was dissolved in corn oil at a concentration of 20 mg/ml by shaking at 65°C for one hour (tube protected from light with tin foil). The stock was kept at 4°C for up to a week. Before injecting, tamoxifen was warmed at 65°C for 10min. A 27-gauge needle was used to do a single intra-peritoneal injection of tamoxifen, 100mg per kg of body weight^28^.

#### DREADD agonist Deschloroclozapine (DCZ)

DCZ dosage was determined based on previous studies^45^. Specifically, 100mg DCZ (Sigma, cat# SML3651) were administered per kg of animal body weight. To this aim, DCZ was dissolved in 2% dimethyl sulfoxide (DMSO) in saline, to achieve a working solution to be injected at 0.1ml/animal.

#### Kainic acid (KA)

KA (Sigma, cat# K0250) was prepared in normal saline and was administered via intra-peritoneal route at a dosage of 20mg/kg of mouse body. This dosage was enough to induce sufficient increase in neuronal activity without any seizures, for which mice were monitored throughout the experimental procedure^42,43^.

### AAVs constructs and production

We used AAVs for mildly increasing or decreasing neuronal Foxg1 expression levels, as well as for altering neuronal excitability.

AAVs employed in this study include:

scAAV_hSyn1p-EmGFP-miR(-);
scAAV_hSyn1p-EmGFP-miR.aFoxg1.1690;
scAAV_hSyn1p-EmGFP-miR.aFoxg1.1694;
ssAAV_hSyn1p-dALPP.P2A.EmGFP-miR(-);
ssAAV_hSyn1p-hM3D(Gq).P2A.EmGFP-miR(-);
ssAAV_hSyn1p-hM4D(Gi).P2A.EmGFP-miR(-);
ssAAV_hSyn1p-hM3D(Gq).P2A.EmGFP-miR.aFoxg1.1690;
ssAAV_hSyn1p-hM4D(Gi).P2A.EmGFP-miR.aFoxg1.1694.
Full sequences of their cargo plasmids are reported in Supplementary Table 1.

These AAVs were pseudotyped as BBB-permeant, CPP21 particles^46^. They were generated and titrated in house as described^47^.

### Surgery/AAVs injections

All surgical procedures took place at P30 unless otherwise specified. Mice were anesthetized by intraperitoneal administration of xylazine (20 μg/gram of body mass) and ketamine (80 μg/gram of body mass) and were transferred to a heated surface (at 37°C) to prevent hypothermia. Afterwards, a single AAV injection was done into the retro-ocular sinus, carried by a 300µl zero dead volume insulin syringe, equipped with 31-gauge needle. To ease the access of the retro-orbital vein, slight pressure was applied on both sides of the eye in order to make the eyeball protrude, paying attention not to damage its walls. The required number of recombinant AAVs (10^8^ viral genomes/mouse) was resuspended in a PBS volume not exceeding 100ul and then injected into the vein. The injection was performed slowly, in approximately 10 seconds, and the needle was left in the venous sinus for another 5 seconds and then gently withdrawn. Once AAV administration was completed, mice received a subcutaneous injection of non-steroidal anti-inflammatory Carprofen (5μg per gram of body mass) and an intraperitoneal injection of antibiotic Baytril (5μg per gram of body mass) to prevent any discomfort/pain or infections. Immediately afterwards, mice were transferred to a hot plate set at 37°C and monitored for the recovery. About 30 minutes after they woke up, mice were transferred back to their respective home cages and monitored regularly for infection symptoms (hypomobility or hyperactivity) till the day of behavioral testing.

### Fear conditioning

Contextual fear conditioning (Figure 1) was performed in test chambers (UgoBasile 46000-596) equipped with an electrified grid floor and dual video camera (for visible and infrared light) connected with AnyMaze software for the automated quantification of time of immobilization of the animal. Mice were trained at P49, 16-20 hrs after tamoxifen administration, in the contextual chamber A (grid-floor with black and white square walls). During the training session, mice were kept in the chamber for 2min, then they received three 0.5mA foot-shocks, each lasting 2sec and spaced by 1min intervals, and finally, 1min after the last foot-shock, they were removed from the chamber. The next day, mice were placed in the training chamber (context A) and (a) total freezing time (cessation of movement, except for breathing, with threshold set at 2sec), (b) number of freezing episodes, and (c) latency to freeze, were scored during the entire testing session, automatically by the Anymaze software. One day later, mice were also tested in a similar but novel chamber (Context B), which was the same size as context A, equipped with white plastic floor and plain gray walls, where they were scored for the same parameters described above.

A similar contextual fear conditioning paradigm (Figure 3) was followed for the long-term memory retrieval. In this case, mice were trained at P49 in the context A and were tested in the same chamber at P56 for the memory recall, scoring them for total freezing time, number of freezing episodes, and latency to freeze.

As for the chemogenetics experiments (Figure 4), a similar contextual fear conditioning paradigm was also followed. In this case, all groups were trained on P56, and one day later, 20min before the memory retrieval session, DCZ was administered to either activate or inhibit neuronal activity of transduced cells. Saline was given to control animals. Mice were then tested in the training context for memory retrieval.

### Brain dissection

For engram studies, mice subjected to contextual fear conditioning were anesthetized one hour after the last retrieval step via intraperitoneal administration of xylazine (60 μg/gram of body mass) and ketamine (240 μg/gram of body mass). Subsequently, they were perfused transcardially with cold (4°C) phosphate buffered saline (PBS) followed by 4% paraformaldehyde (PFA) in PBS. Immediately afterwards, they were decapitated. Brains were extracted, kept in 4% PFA at 4°C for 24 hours and transferred to 30% sucrose in PBS solution till sectioning.

For KA experiments, KA administered mice were anesthetized as above, perfused with cold PBS and decapitated. Half the brain from each animal was used to dissect neocortex, and hippocampal formation, for RNA extraction. The other half was kept in 4% PFA overnight and then transferred to 30% sucrose solution, till sectioning for immunofluorescence.

### Total mRNA profiling

Total RNA was extracted from neocortex and hippocampal formation by Trizol reagent (Thermofisher) according to manufacturer’s instructions. RNA was precipitated by isopropanol overnight at 321 −80°C, washed with 75% ethanol twice and resuspended in 20µl sterile nuclease-free deionized water. Spectrophotometric measurements (NanoDrop ND-1000) and agarose gel electrophoresis were used to estimate RNA concentration, quality and purity. Before retro-transcription, all RNA samples were treated with TURBO DNA-free^TM^Kit (ThermoFisher Scientific) to remove any residual DNA. For each experimental session at least 500ng of purified RNA was retro-transcribed by SuperScript-III^TM^ (Invitrogen) in the presence of random hexamers, according to the manufacturer’s instructions. 1/100 of the resulting cDNA was used as a substrate of any subsequent qPCR reaction. Negative controls were run on RT (-) cDNA preparations and included in each PCR session. qPCR reactions were done using SsoAdvanced SYBR Green Supermix^TM^ platform (Biorad) according to the manufacturer’s instructions. Three technical replicates were run per each cDNA sample, analyzed against absolute standards and averaged. These averages were then normalized against *Gapdh*, and further normalized against controls.

qPCR oligonucleotides are listed below:

GAPDH-Forward: 5’ ATC TTC TTG TGC AGT GCC AGC CTC GTC 3’
GAPDH-Reverse: 5’ GAA CAT GTA GAC CAT GTA GTT GAG GTC AAT GAA GG 3’
Foxg1-midF- Forward: 5’ GAC AAG AAG AAC GGC AAG TAC GAG AAG C 3’
Foxg1-midF- Reverse: 5’ GAA CTC ATA GAT GCC ATT GAG CGT CAG G 3’
mFos/F1N- Forward: 5’ CTG ACA GAT ACA CTC CAA GCG GAG ACA G 3’
mFos/R1N- Reverse: 5’ ACA TCT CCT CTG GGA AGC CAA GGT CAT C 3’
ARC-Forward: 5’ CCC TCA TCT GTC TGC CCT GG 3’
ARC-Reverse: 5’ ACC CAA AGA GCC CTG GAC AC 3’

### Histology and Immunostaining

Brains were sectioned using a microtome into 50µm thick coronal sections (for engram studies) and 20µm thick sagittal sections (for kainic acid treatment experiments). These sections were collected in 0.01% sodium azide in PBS for long-term storage at 4°C. For Foxg1, cFos and GFP immunostaining, sections were incubated in permeabilization buffer (0.1% Triton-X100 in PBS) for 1hour at RT, then in blocking mix (5% bovine serum albumin in PBS) for 1hour at RT and finally stained with rabbit polyclonal anti-Foxg1 (a gift from G.Corte, 1:200), mouse anti-cFos (Abcam 208942, 1:1000), and chicken anti-GFP (GeneTex GTX13970, 1:4000) antibodies in blocking mix overnight at 4°C. All sections were then washed 3 times for 10min with washing solution (0.05% Triton-X100 in PBS) and then incubated with secondary antibodies, anti-mouse Alexa 647 (Invitrogen, 1:600) and anti-chicken Alexa 488 (Invitrogen, 1:600) in blocking mix, along with DAPI, for 2hours at room temperature. Sections were then washed 3 times for 10min with washing solution, mounted on Superfrost Plus slides, coverslipped by Vectashield mounting medium (Vector), and kept at 4°C till imaging. Confocal images were finally obtained with a Nikon A1 confocal microscope.

### Image processing and analysis

Cell counts were calculated by an operator blind to experimental conditions. At least 8-9 images were analyzed from each mouse brain and for accurate regional analysis these images were aligned to mouse Allen Brain Atlas. Cells stained for DAPI, EGFP, cFos and tdTomato (in *Ai9* mice) were quantified using the quPath software, with cell detection parameters optimized and validated against negative controls.

Quantification of Foxg1 expression in active (cFos^+^) vs non-active (cFos^-^) neuronal populations upon KA administration was done by using Volocity software. Per each animal, absolute Foxg1 signal intensity was calculated in both populations and cFos^+^ cells Foxg1 intensity was then normalized against cFos^-^ cells.

### Statistical analysis

Primary data originating from each experimental session were pre-normalized against the control average of such session and subsequently merged with data form other sessions, for further analysis. All these data are shown in Supplemental Table 2. Statistical significance of differences among behavioral (freezing time, number of freezing episodes and latency) and engram parameters (ROR and absolute percentages of cFos^+^ and TdTomato^+^ cells) was calculated by one-tailed t-test (paired and unpaired in case of intra- and inter-group comparisons). In any case the number of statistical replicates equaled the number of animals analyzed.

## DATA AVAILABILITY

All relevant data can be found within the article and its supplementary information.

## SUPPLEMENTARY INFORMATION

### LEGENDS TO SUPPLEMENTARY FIGURES

**Supplementary Figure 1.**
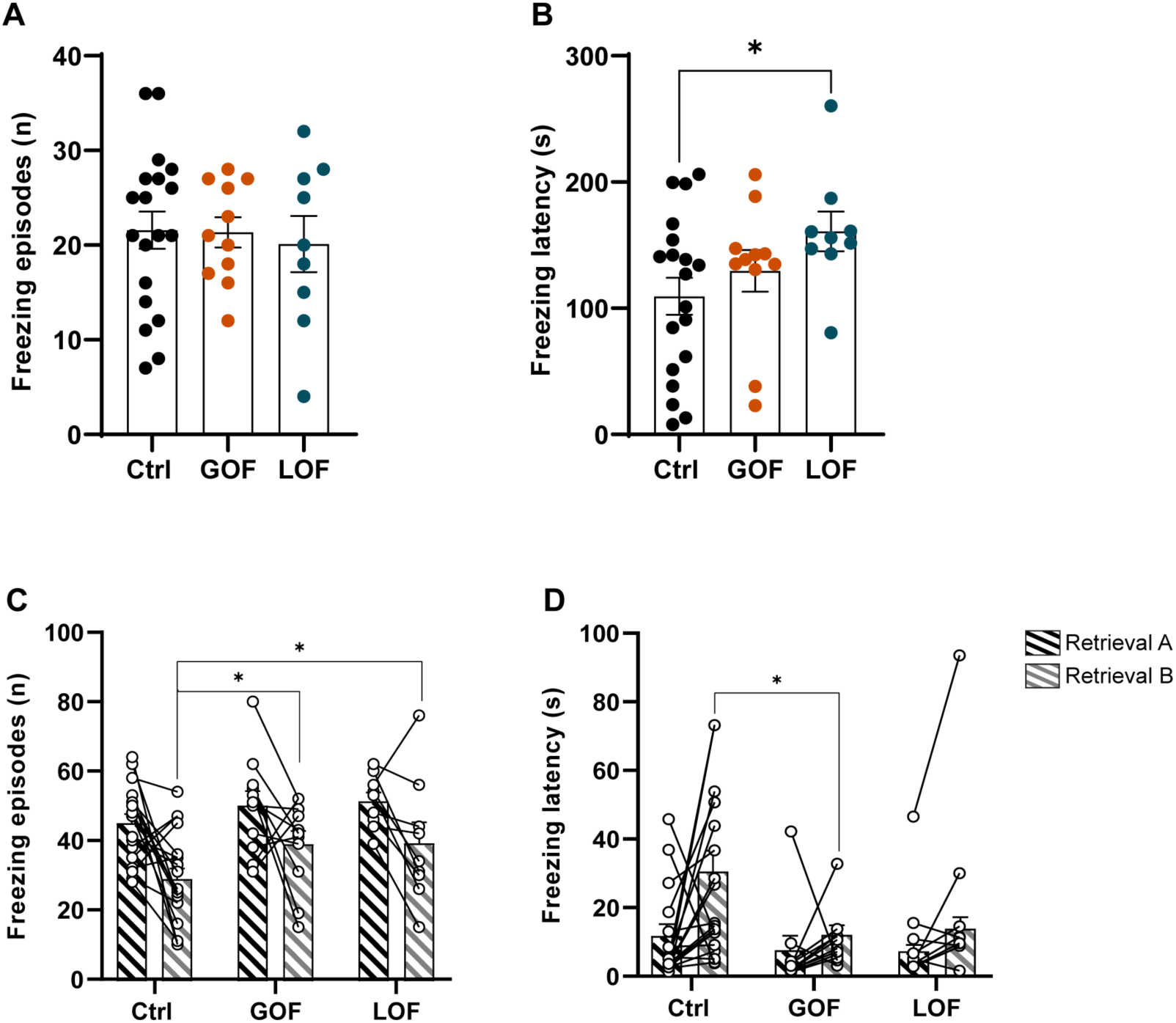
Supplementary data linked to Figure 1A-E. **A-B**: Total freezing episodes and freezing latency during training session of contextual fear conditioning (CFC) in Ctrl, GOF, and LOF groups. **C-D:** Total freezing episodes and freezing latency during retrieval session, in the training context (Retrieval A) and a novel context (Retrieval B). **A-D:** Error bars indicate standard error of the mean (SEM), n = 19 (Ctrl), 11 (GOF), 9 (LOF). Statistical significance of results was evaluated by t-test assay (unpaired between groups, paired within groups, one-tailed). * *p*<0.05.

**Supplementary Figure 2.**
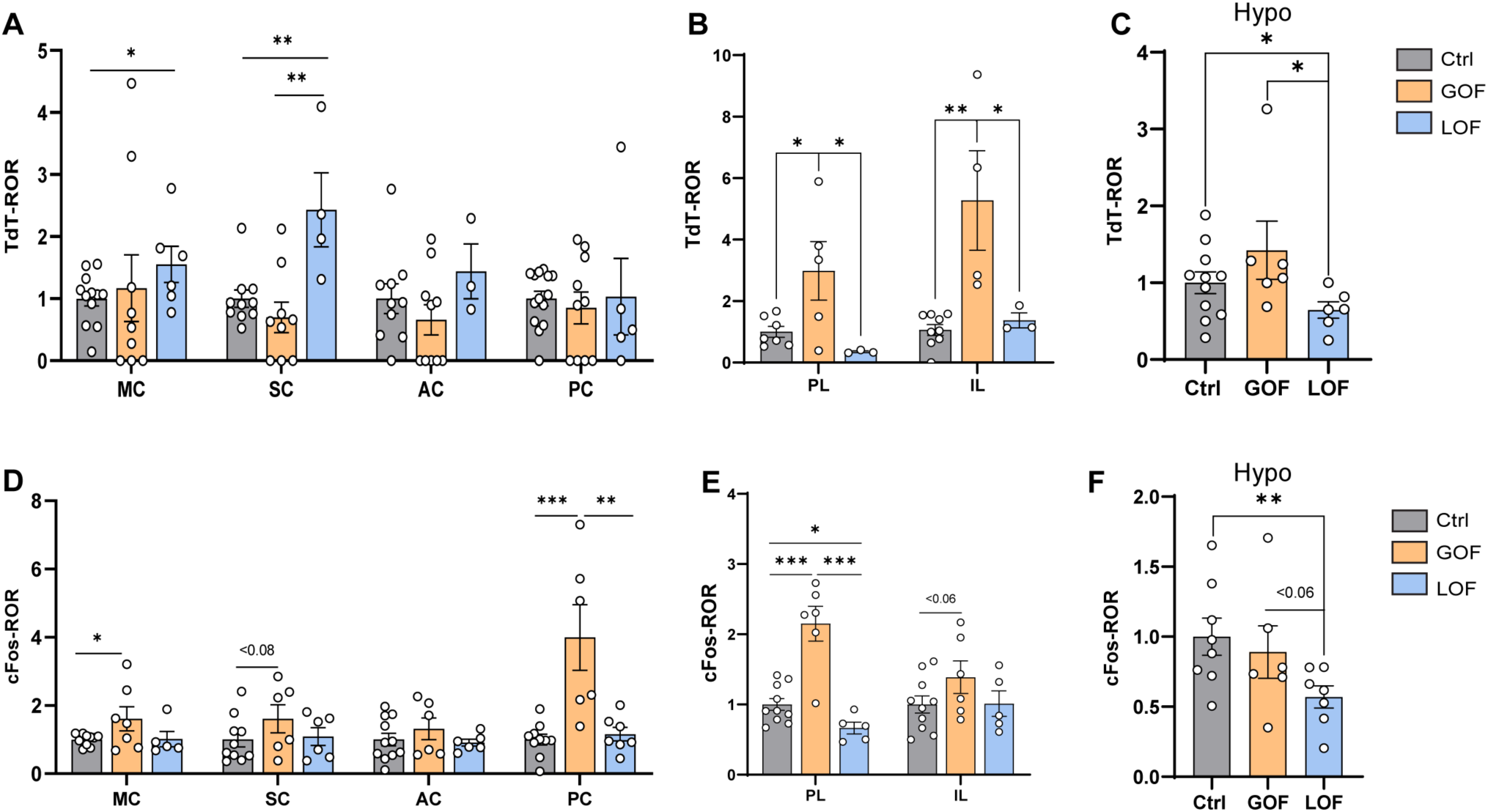
Supplementary data linked to Figure 1F-T. **A-F**: quantification of Ctrl-normalized TdT-ROR (**A-C**) and cFos-ROR (**D-F**) in selected regions of brains treated with Ctrl, GOF, and LOF AAVs, n = 7-14 mice per Ctrl group, 4-10 mice per GOF group, 3-7 mice per LOF group. Error bars indicate standard error of the mean (SEM). Statistical significance of results was evaluated by t-test assay (unpaired, one-tailed). * *p*<0.05, ** *p*<0.01, *** *p*<0.001. ROR, ratio of ratios; MC, motor cortex; SC, somatosensory cortex; AC, auditory cortex; PC, piriform cortex; PL, pre-limbic cortex; IL, infra-limbic cortex; Hypo, hypothalamus.

**Supplementary Figure 3.**
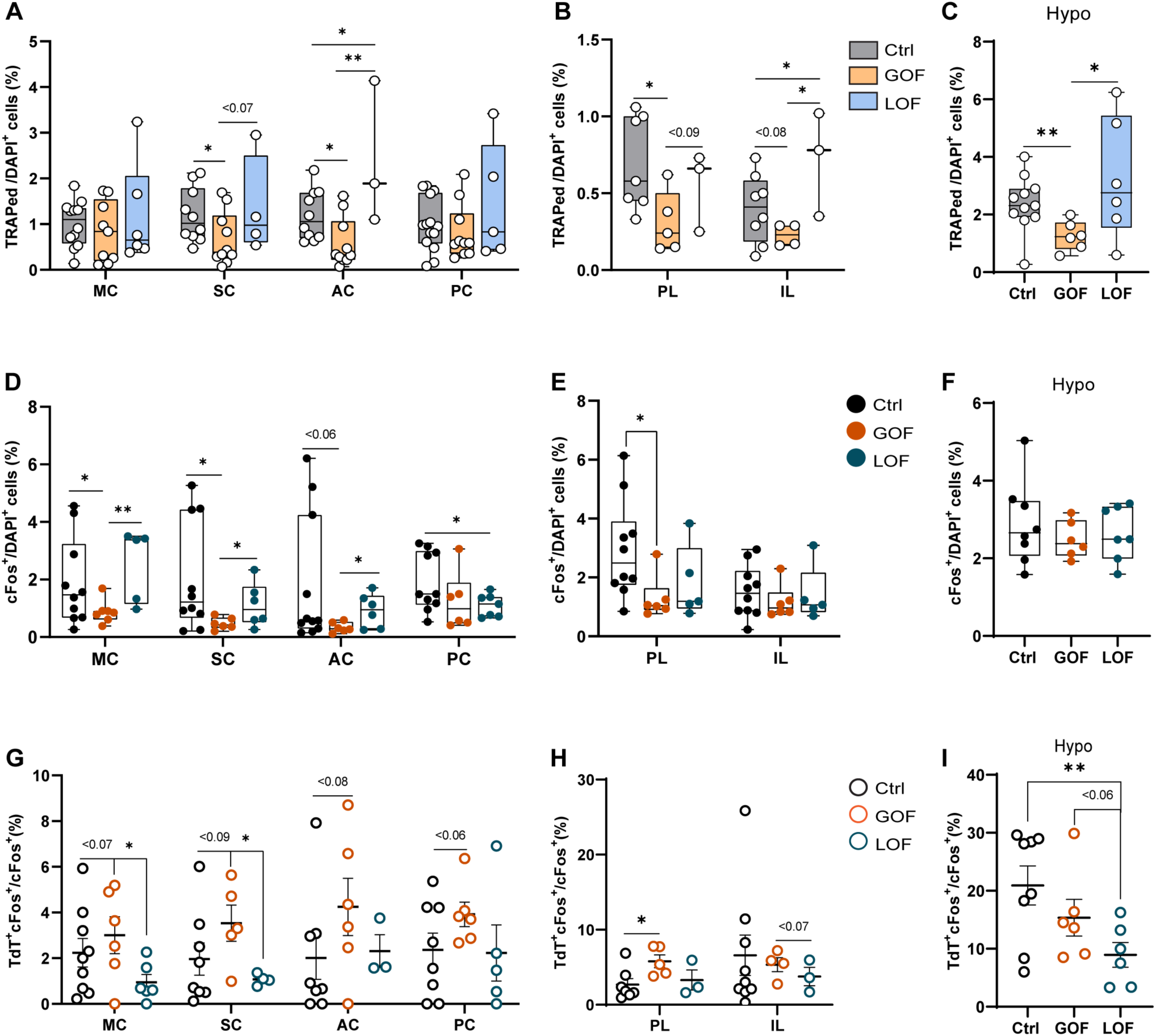
Supplementary data linked to Figure 2. **A-I**: Quantification of TRAPed cells (TdTomato^+^ cells/DAPI^+^ cells, A-C), cFos^+^ cells (cFos^+^ cells/DAPI^+^ cells, D-F), and double positive cells (TdTomato^+^ cFos^+^ cells/cFos^+^ cells, G-I), all displayed as absolute percentages. n = 7-12 (Ctrl), 5-8 (GOF), 4-7 (LOF). Error bars indicate standard error of the mean (SEM). Statistical significance of results was evaluated by t-test assay (unpaired, one-tailed). * *p*<0.05, ** *p*<0.01, *** *p*<0.001. MC, motor cortex; SC, somatosensory cortex; AC, auditory cortex; PC, piriform cortex; PL, pre-limbic cortex; IL, infra-limbic cortex; Hypo, hypothalamus.

**Supplementary Figure 4.**
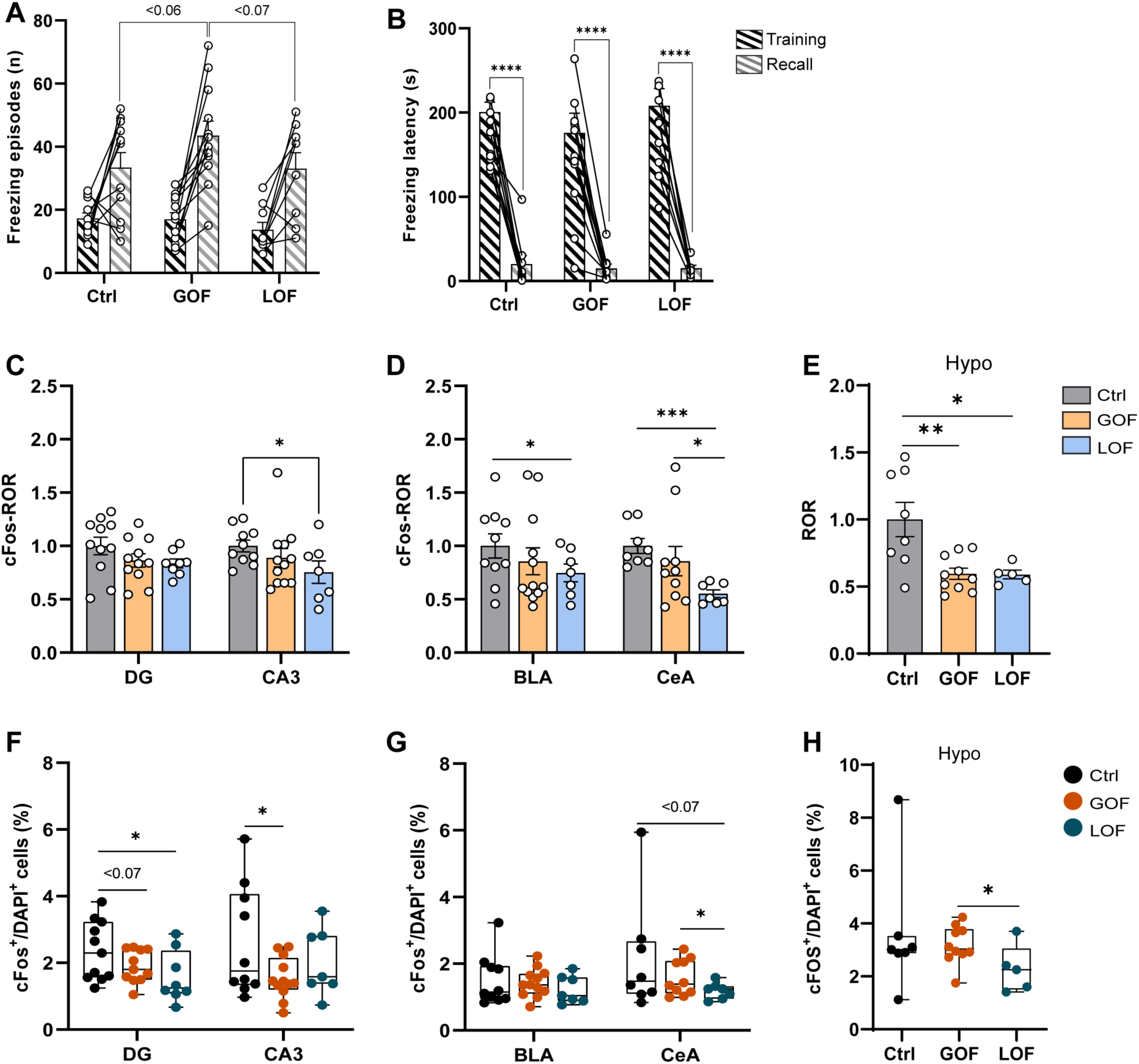
Supplementary data linked to Figure 3. **A-B**: Distribution of total freezing episodes and freezing latency in Ctrl, GOF and LOF groups during training and late memory recall in training context. **C-H:** Quantification of Ctrl-normalized cFos-ROR and cFos^+^/DAPI^+^ cell ratio in brains of Ctrl, GOF, and LOF groups. **A-B:** n = 11 (Ctrl), 12 (GOF), 9 (LOF). **C-H:** n = 8-11 (Ctr), 9-12 (GOF), 5-8 (LOF). Error bars indicate standard error of the mean (SEM). Statistical significance of results was evaluated by t-test assay (unpaired between groups, paired within groups, one-tailed). * *p*<0.05, ** *p*<0.01, *** *p*<0.001, **** *p*<0.0001. ROR, ratio of ratios; DG, dentate gyrus; CA3, hippocampal CA3 field; BLA, basolateral amygdala; CeA, central amygdala.

**Supplementary Figure 5.**
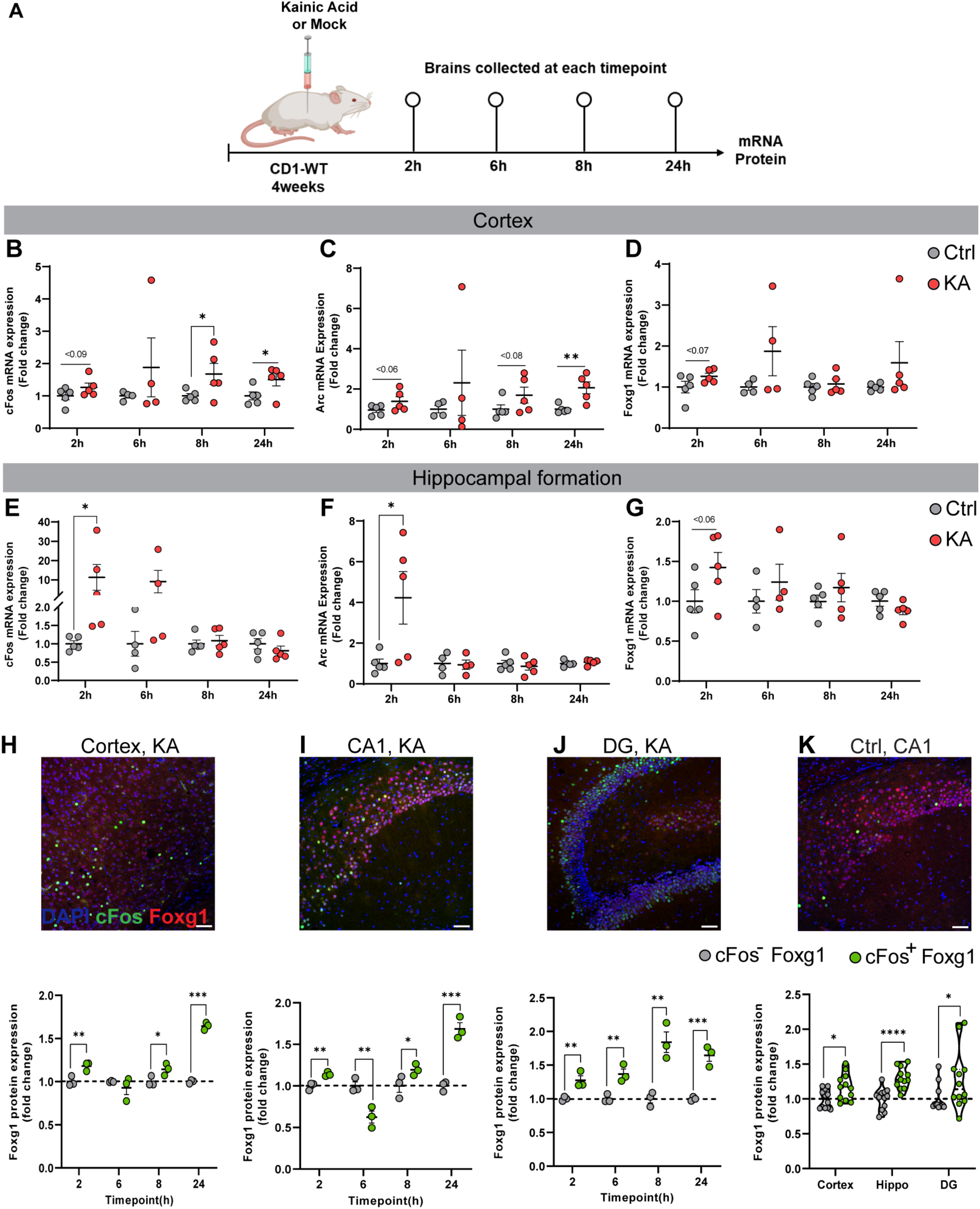
In vivo, activity-dependent Foxg1 modulation. **A**: Experimental design. Mice administered intraperitoneally with 20mg/kg of kainic acid (KA) and their time-matched mock-injected controls (Ctrl) were sacrificed 2h, 6h, 8h, and 24h after drug administration, and profiled for gene expression. **B-G**: Immediate early genes (IEGs) and Foxg1 mRNA levels in cortex and hippocampal formation of KA and Ctrl mice. **H-J**: Representative images of cFos and Foxg1 immunofluorescence, in Cortex, Hippocampus (CA1), and Dentate Gyrus (DG) from KA groups, scale bar 100µm (upper lane). Foxg1 protein expression levels in the above-mentioned structures, specifically within cFos^+^ and cFos^-^ cells, shown as cFos^-^normalized values (lower lane). **K**: Representative image of cFos and Foxg1 immunofluorescence, in Hippocampus (CA1) from Ctrl mice, scale bar 100µm (upper lane). Foxg1 protein expression levels from Ctrl groups, specifically within cFos^+^ and cFos^-^ cells, shown as cFos^-^ normalized values. **B-G**: n = 5 (2h,Ctrl), 4 (6h,Ctrl), 5 (8h,Ctrl), 5 (24h,Ctrl), 5 (2h,KA), 4 (6h,KA), 4 (8h,KA), 5 (24h,KA). **H-J**: n = 3 (per each group). **K**: n = 12 (Ctrl). Error bars indicate standard error of the mean (SEM), Statistical significance of results was evaluated by t-test assay (unpaired between groups, paired within groups, one-tailed). * *p*<0.05, ** *p*<0.01, *** *p*<0.001.

**Supplementary Table 1.**
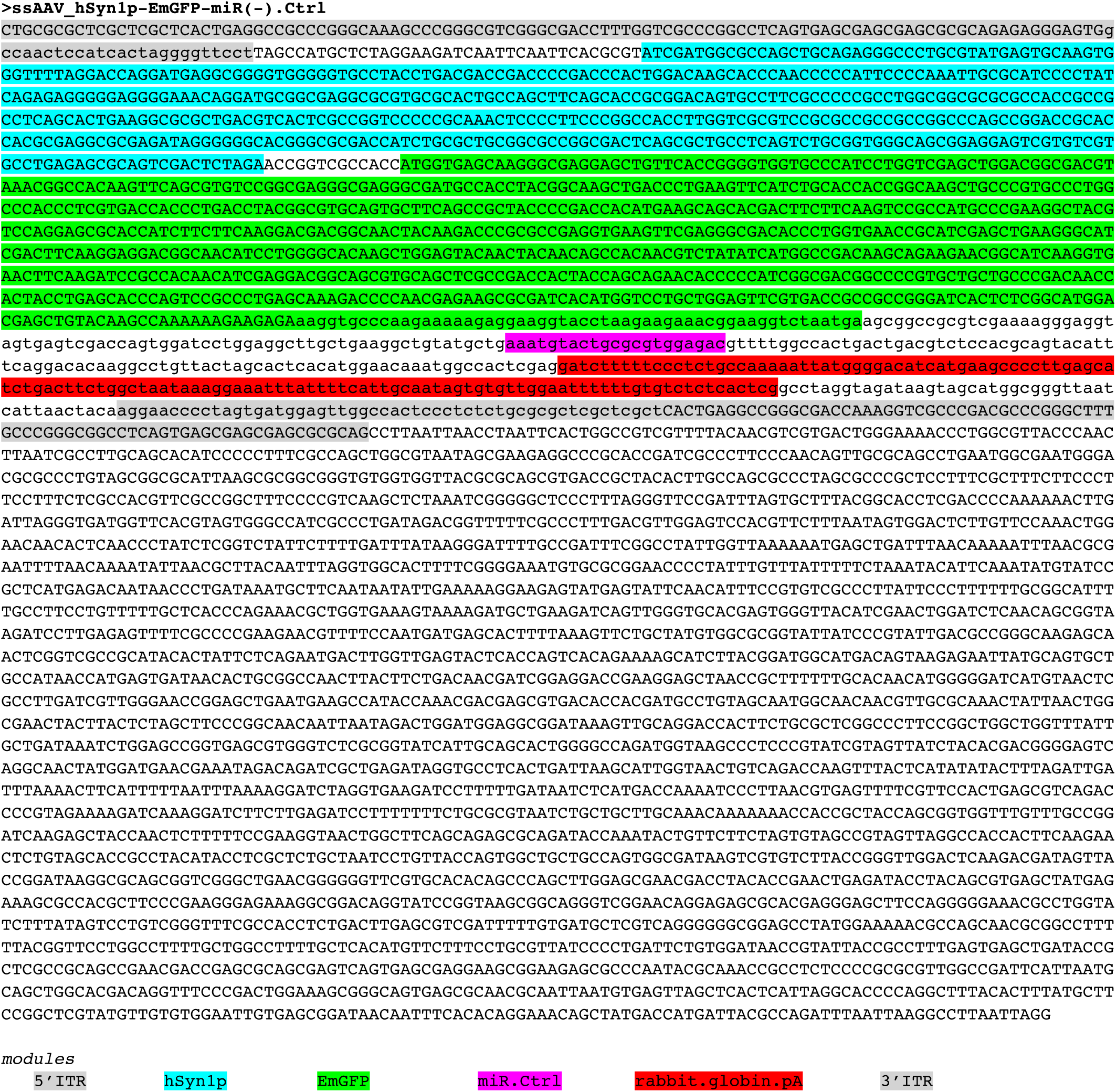
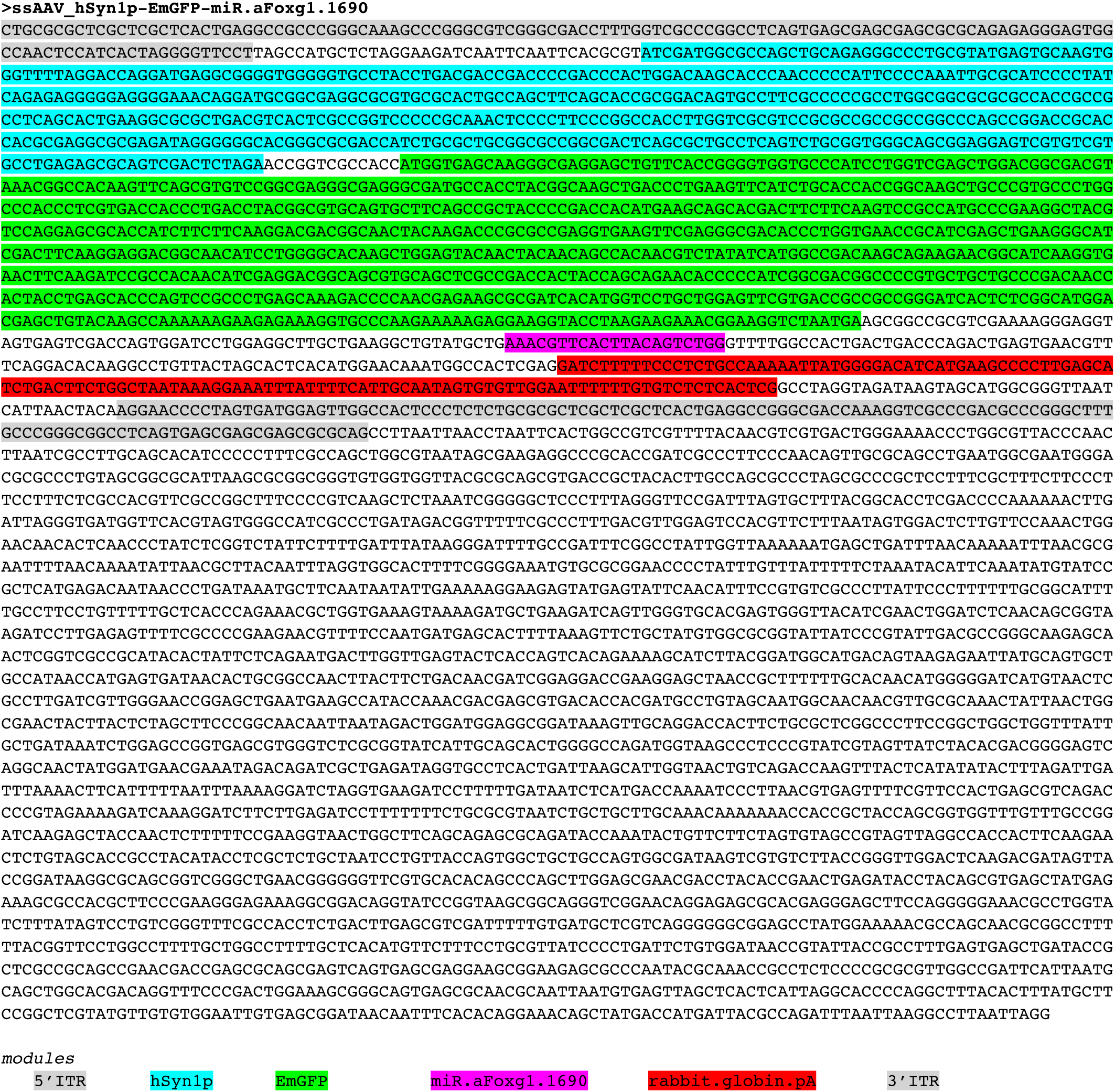
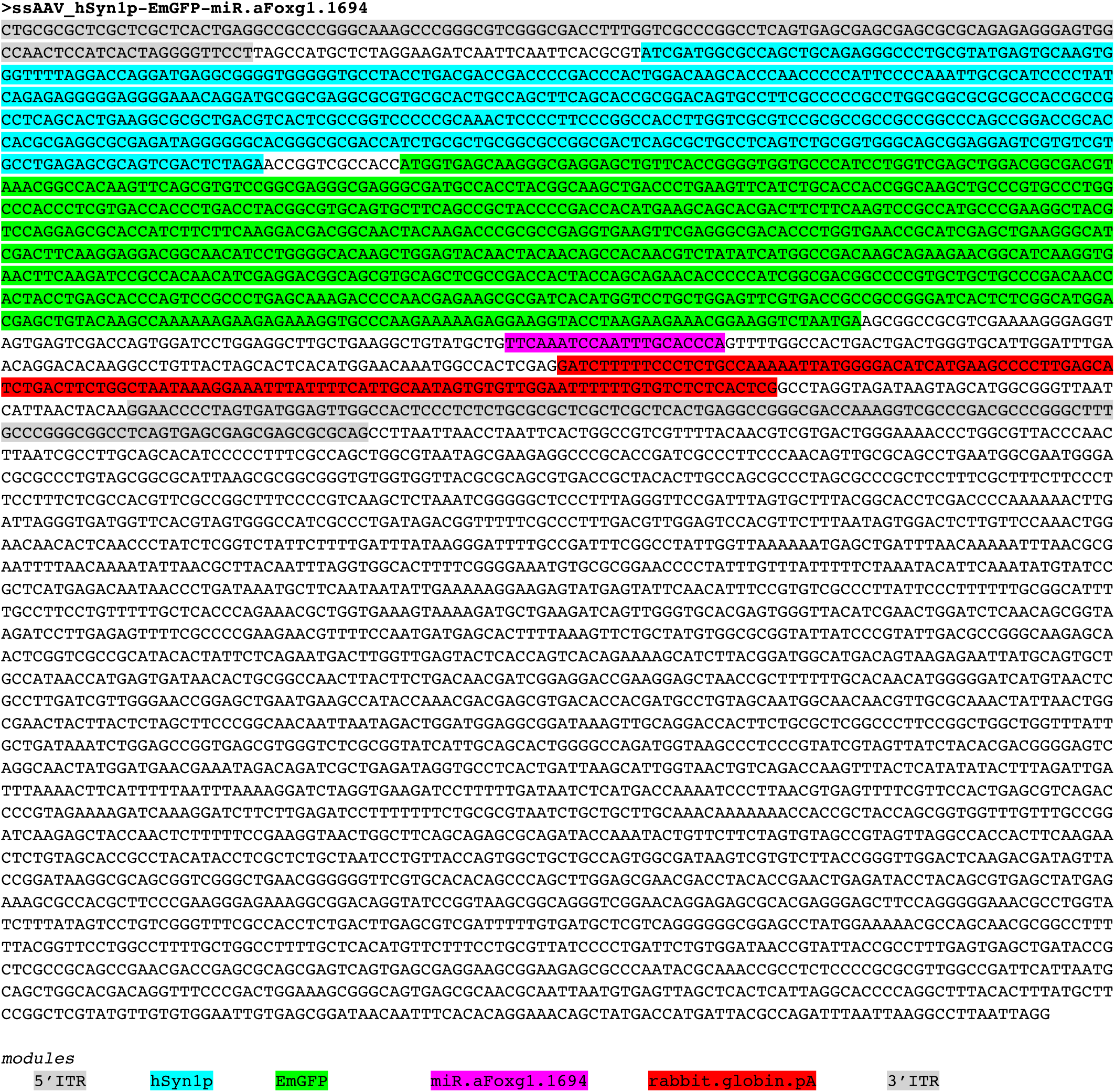
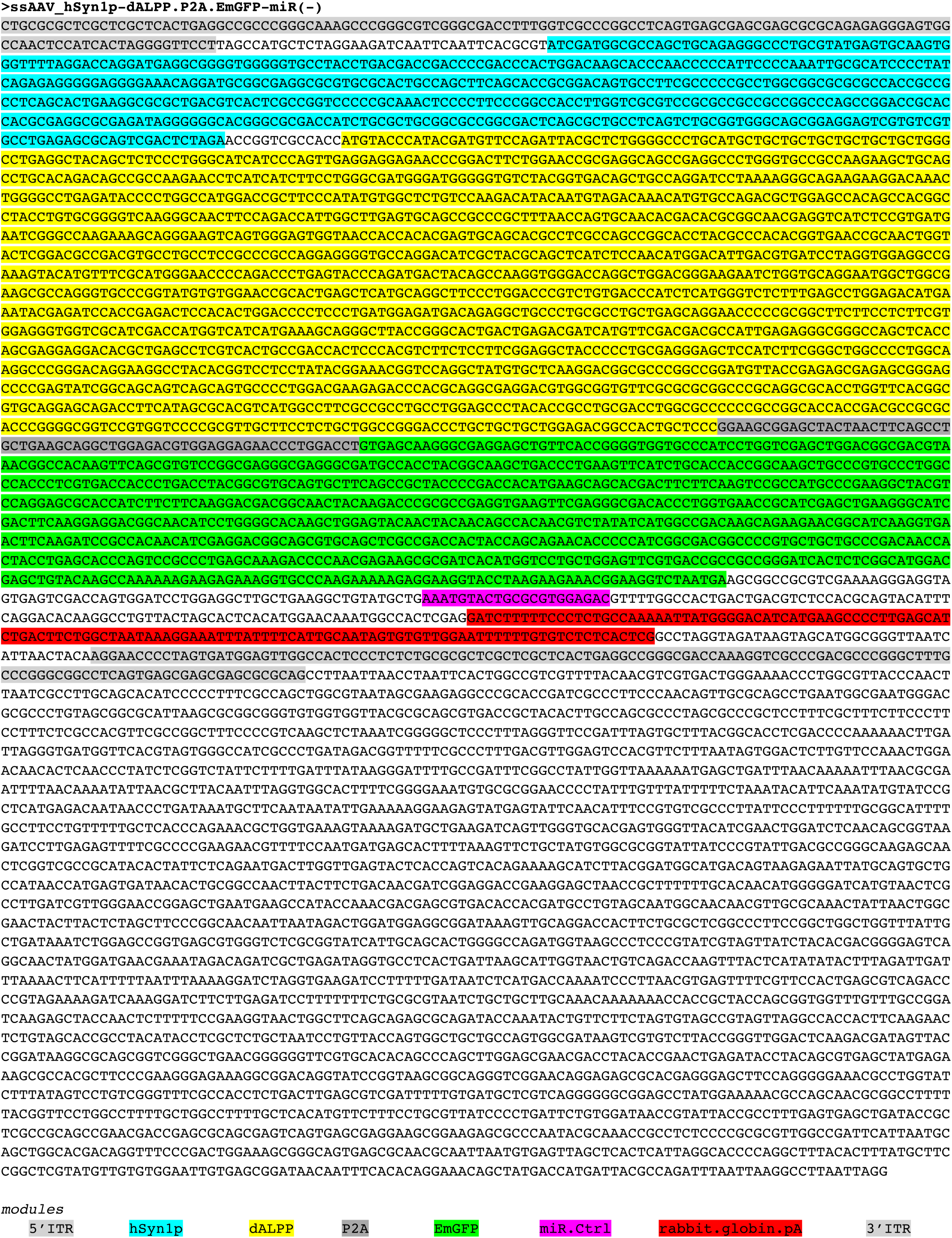
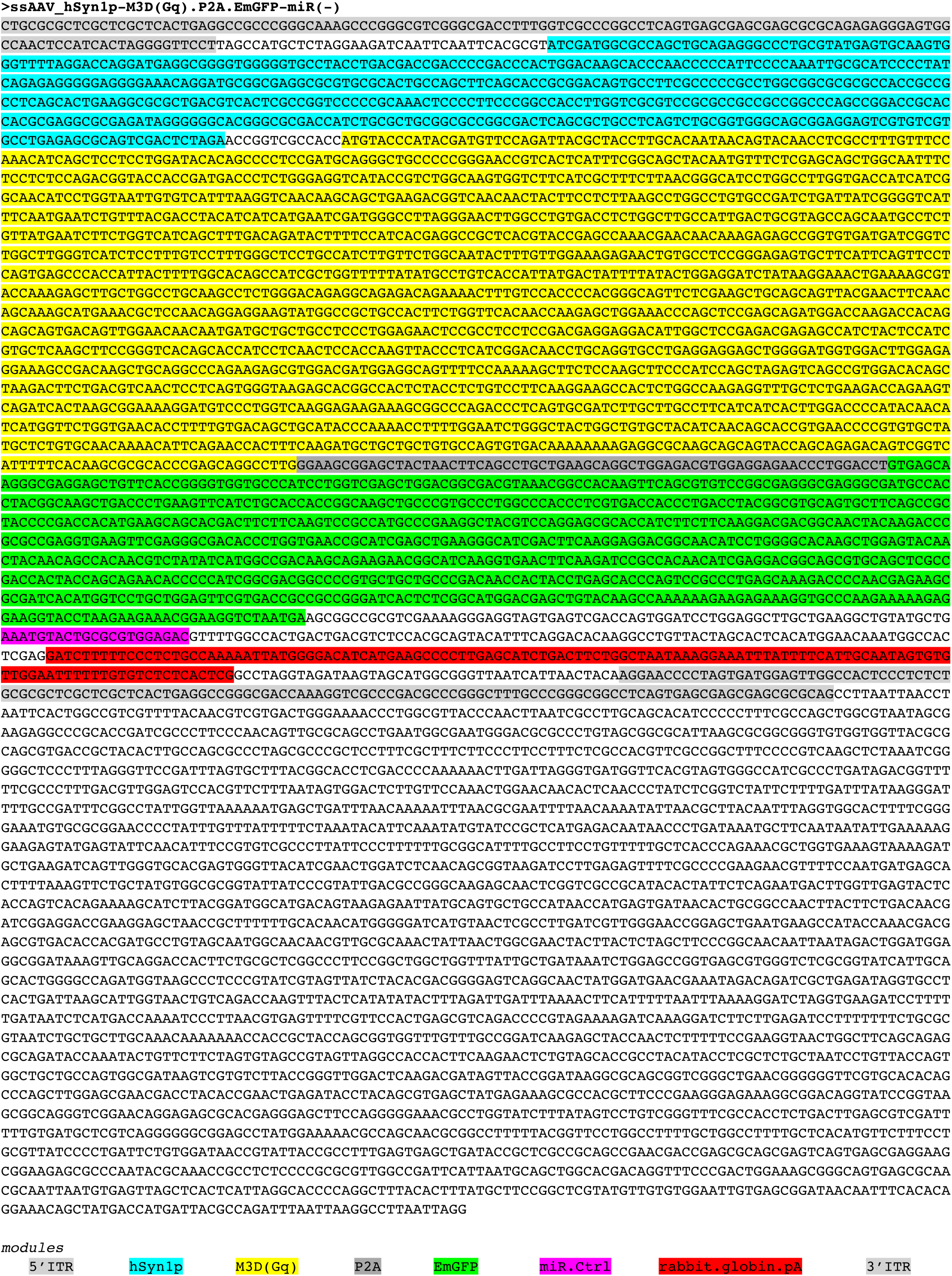
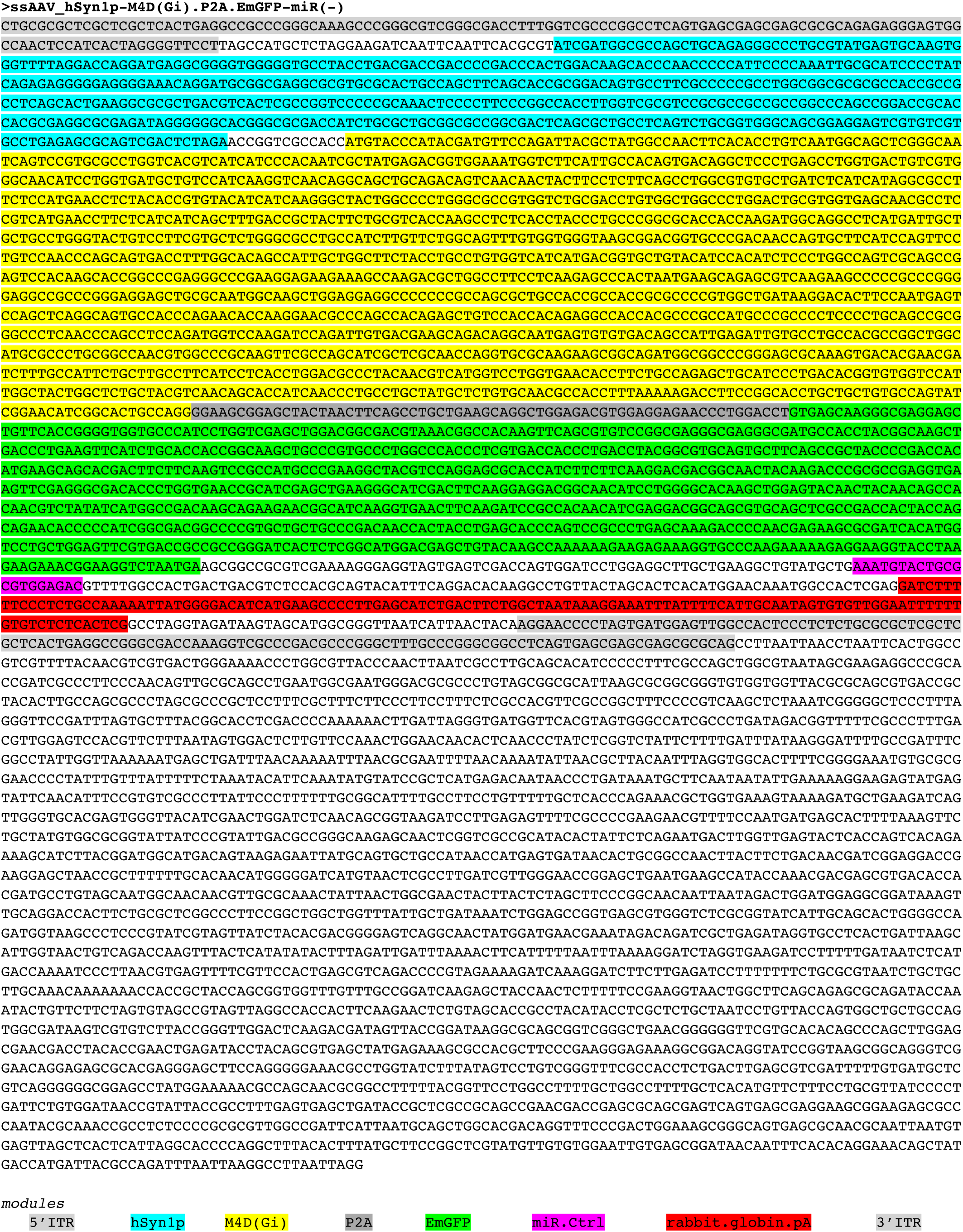
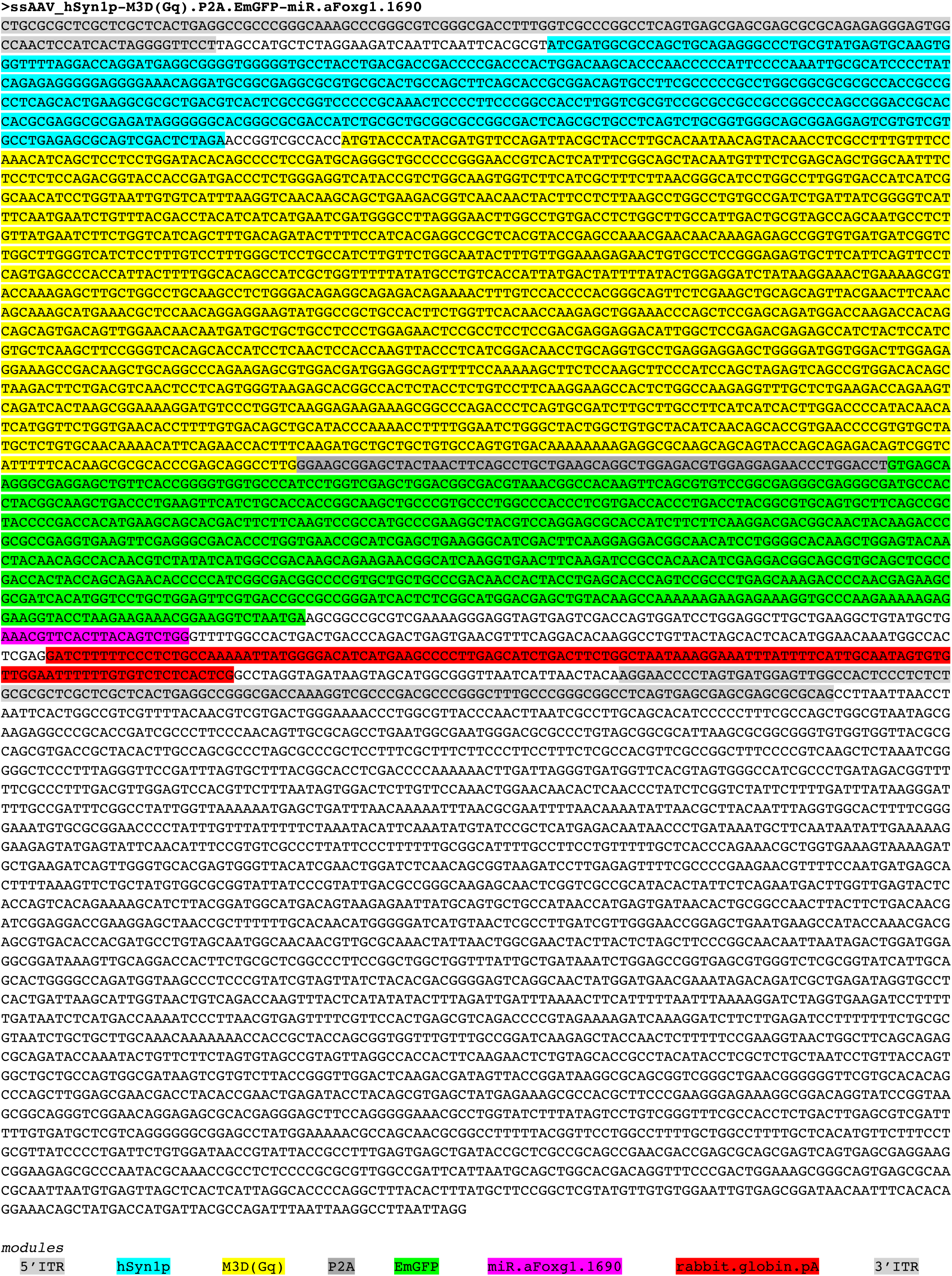
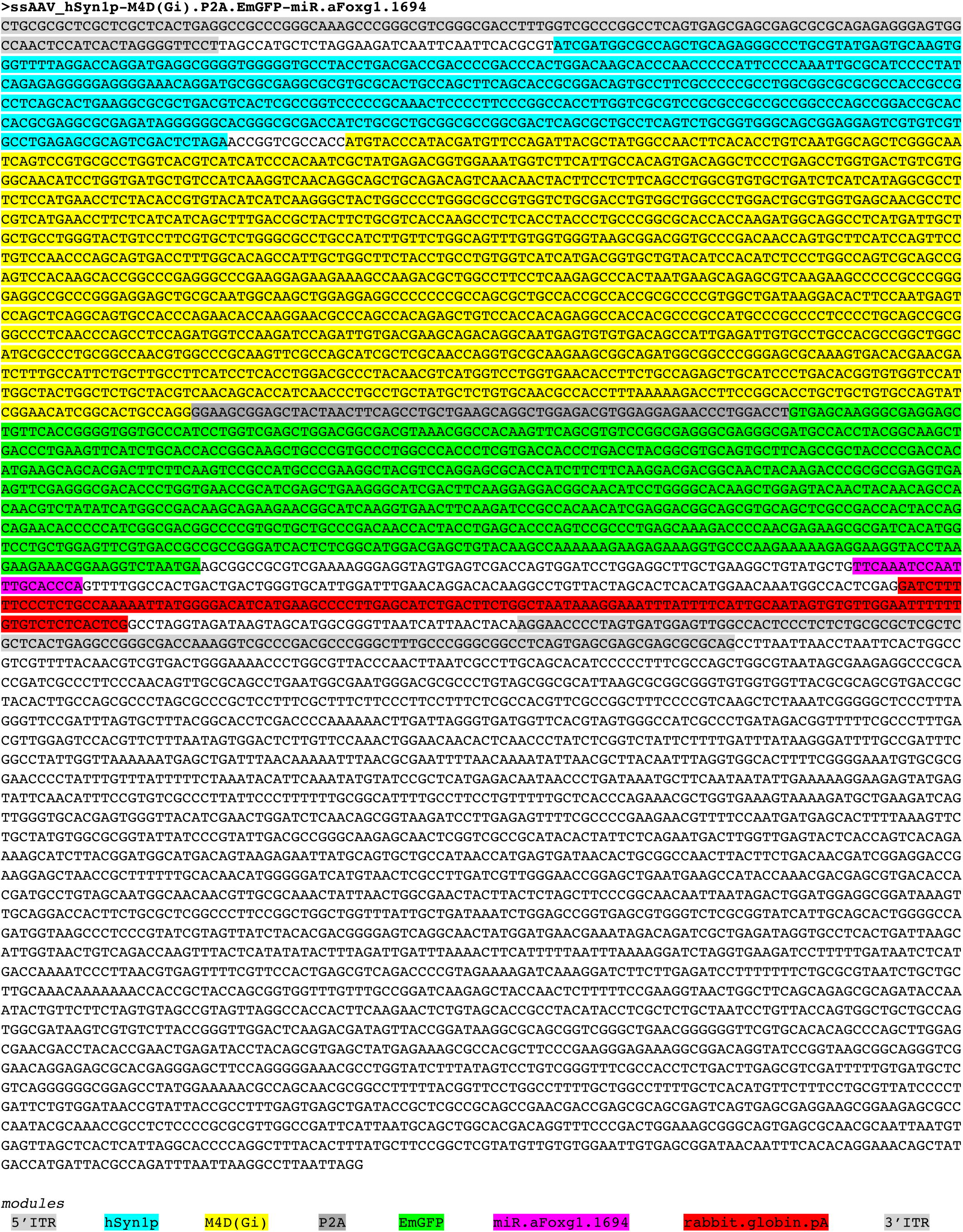
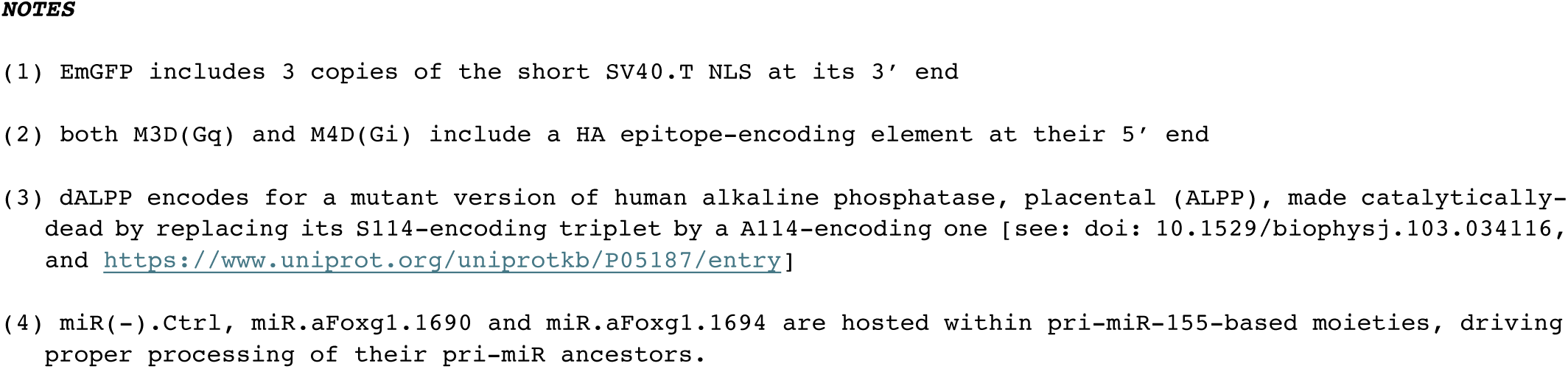
AAV-encoding cargo plasmids.

**Supplementary Table 2.**
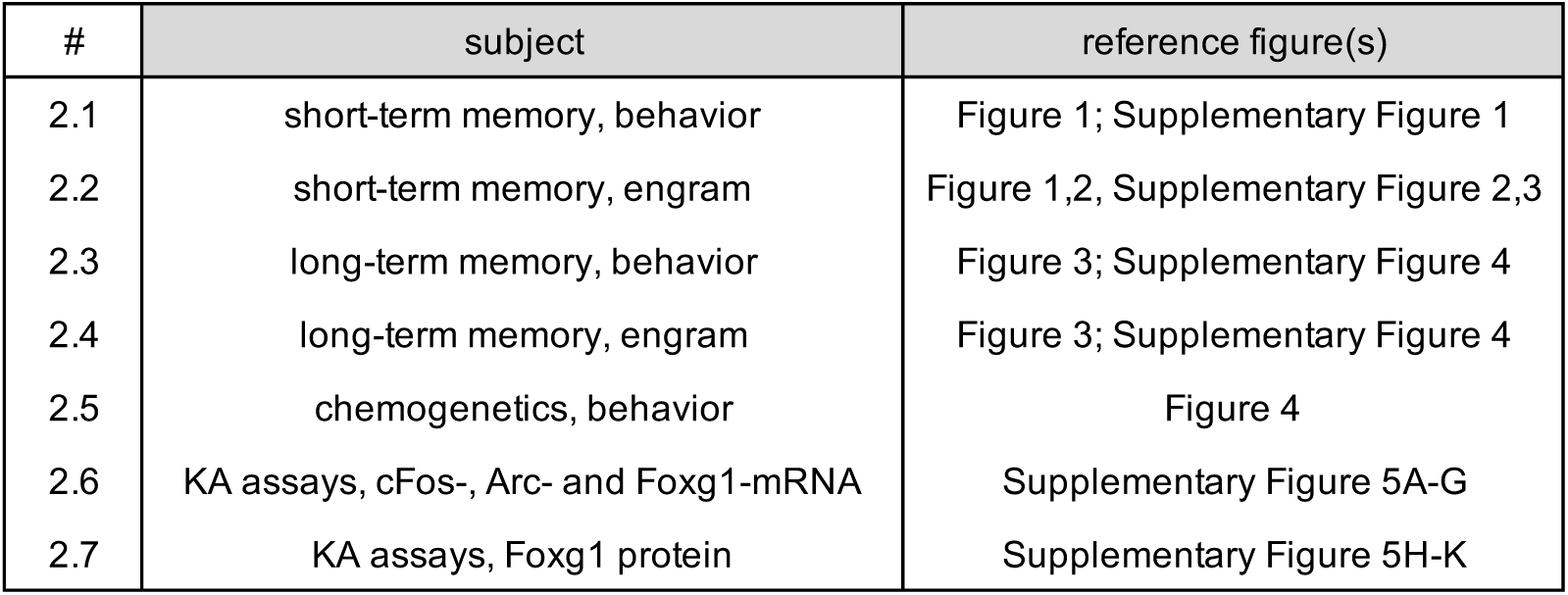
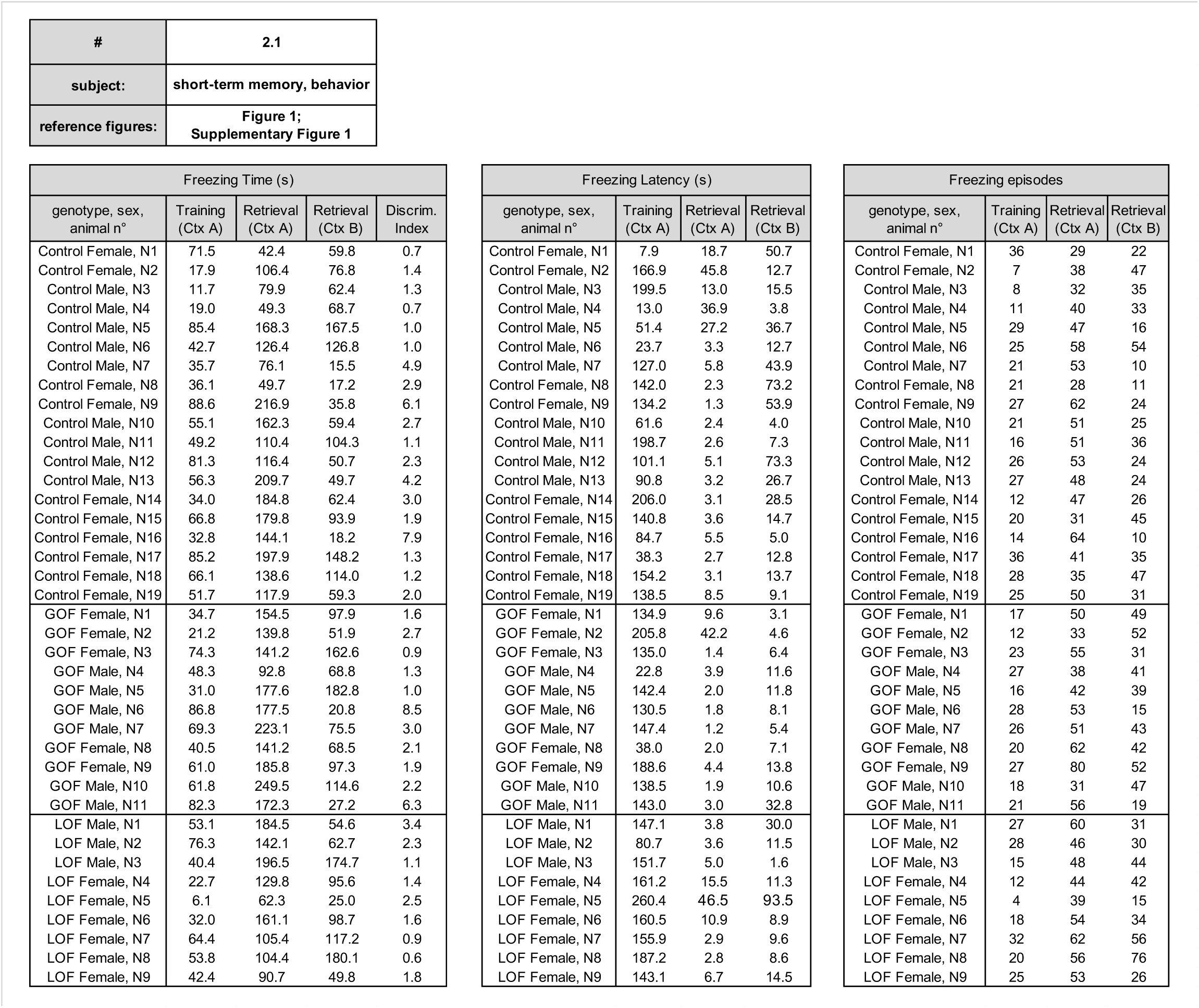
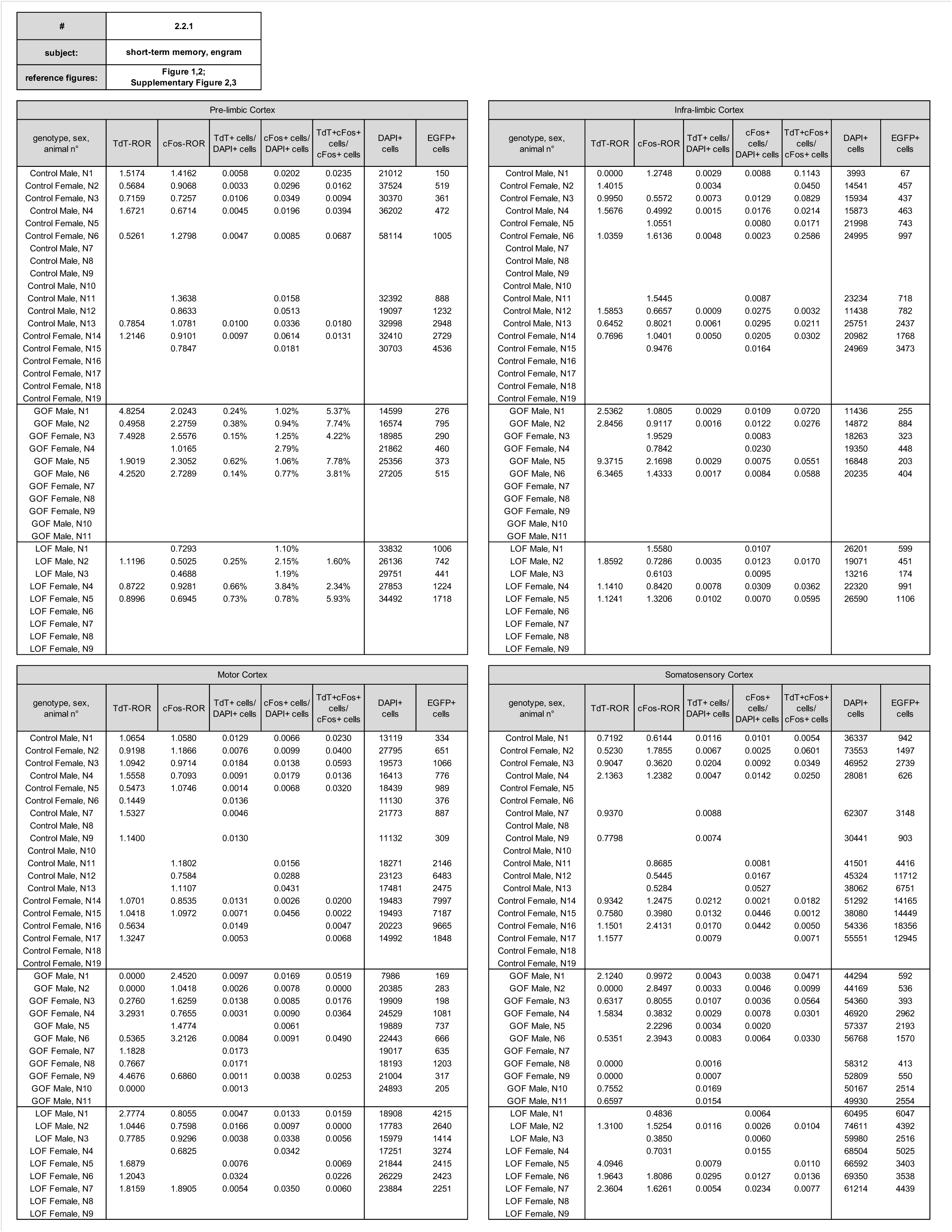
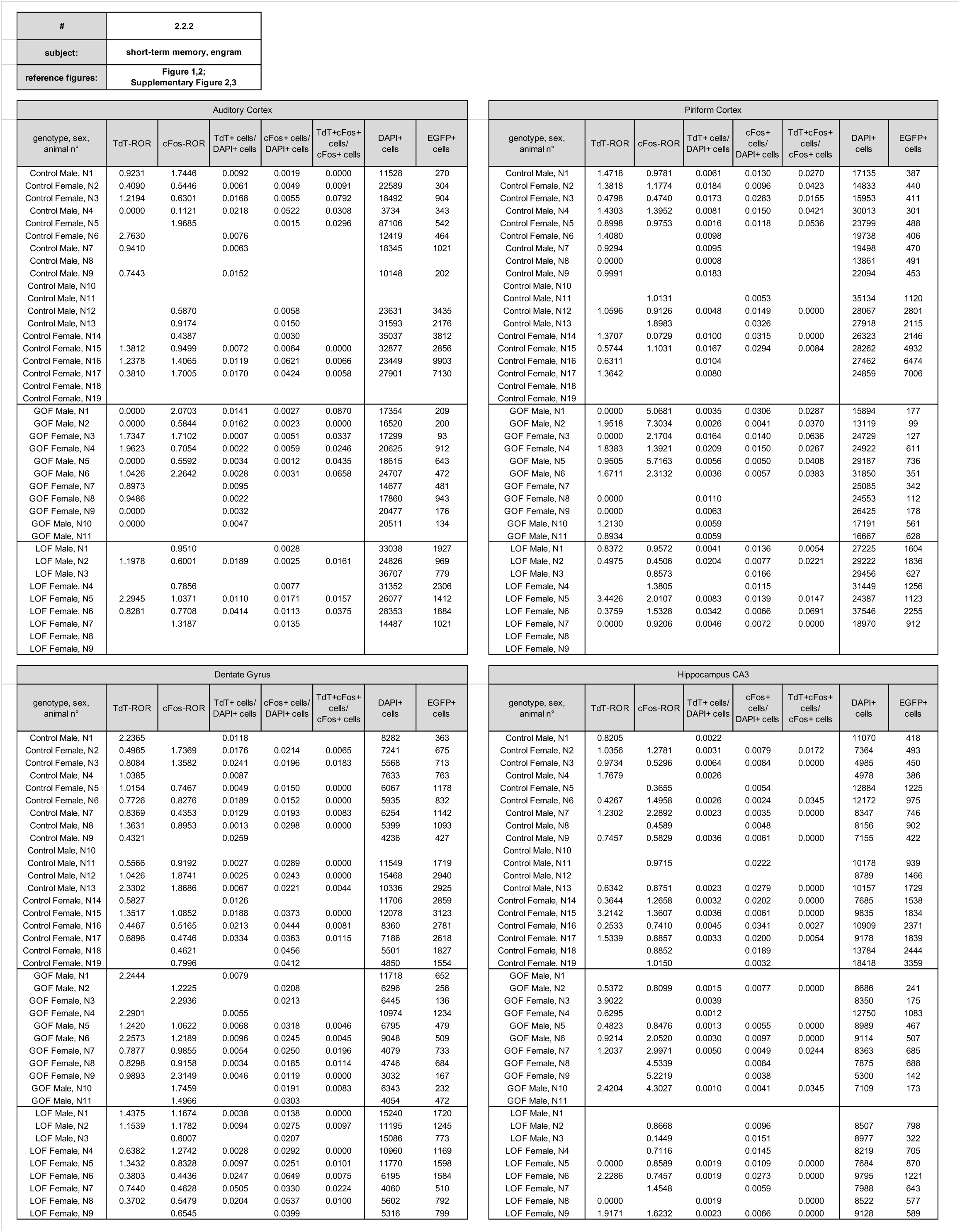
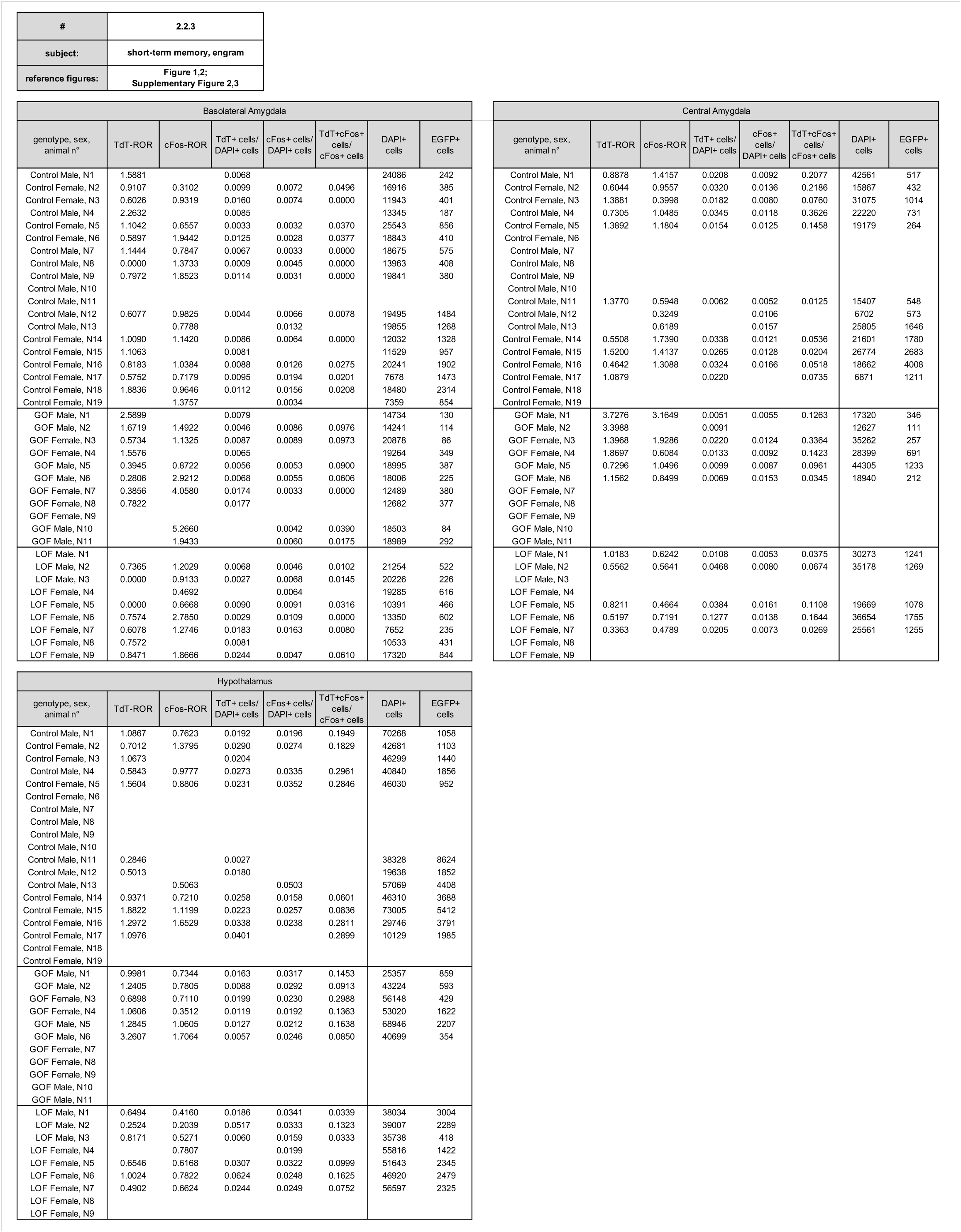
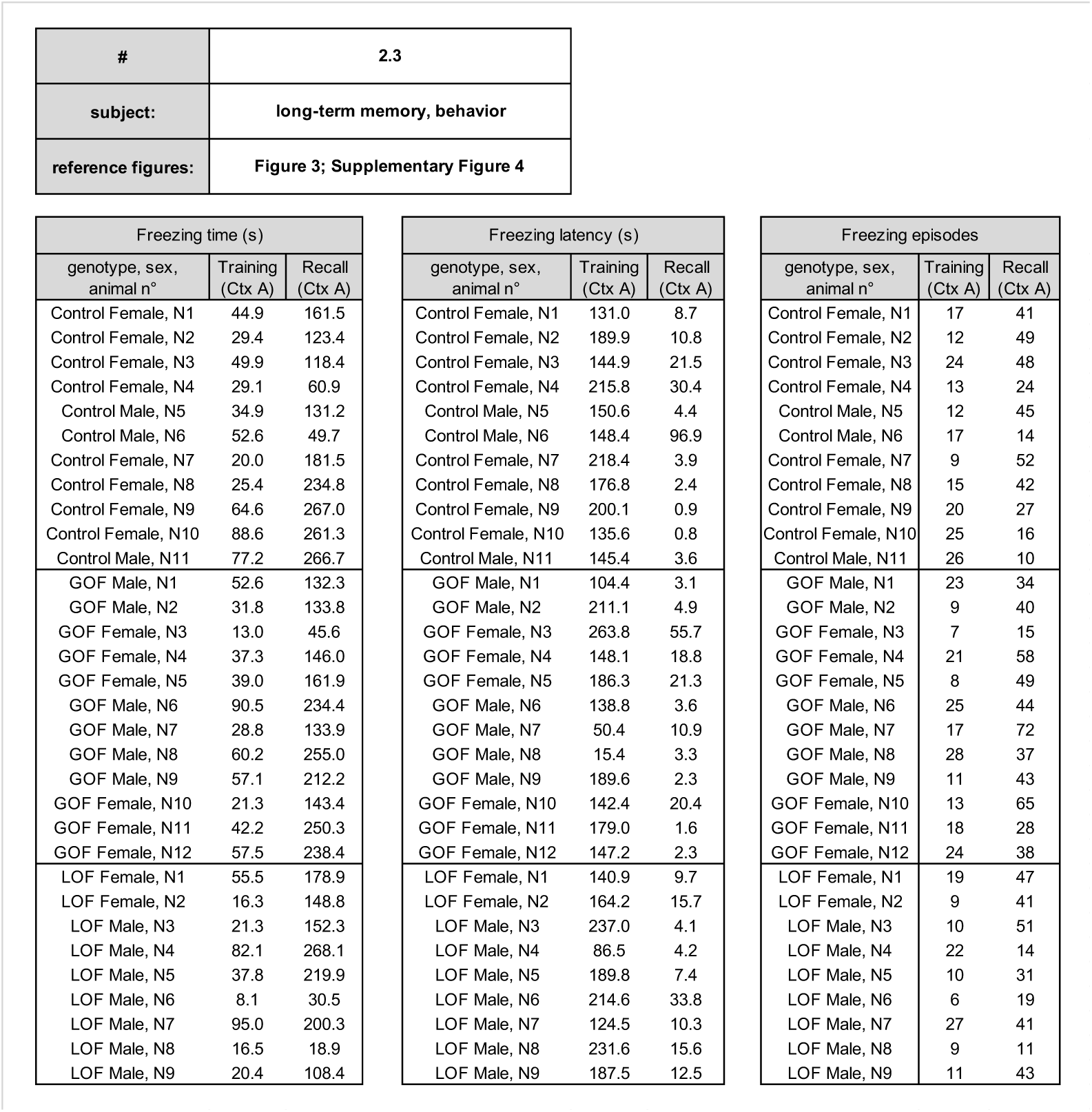
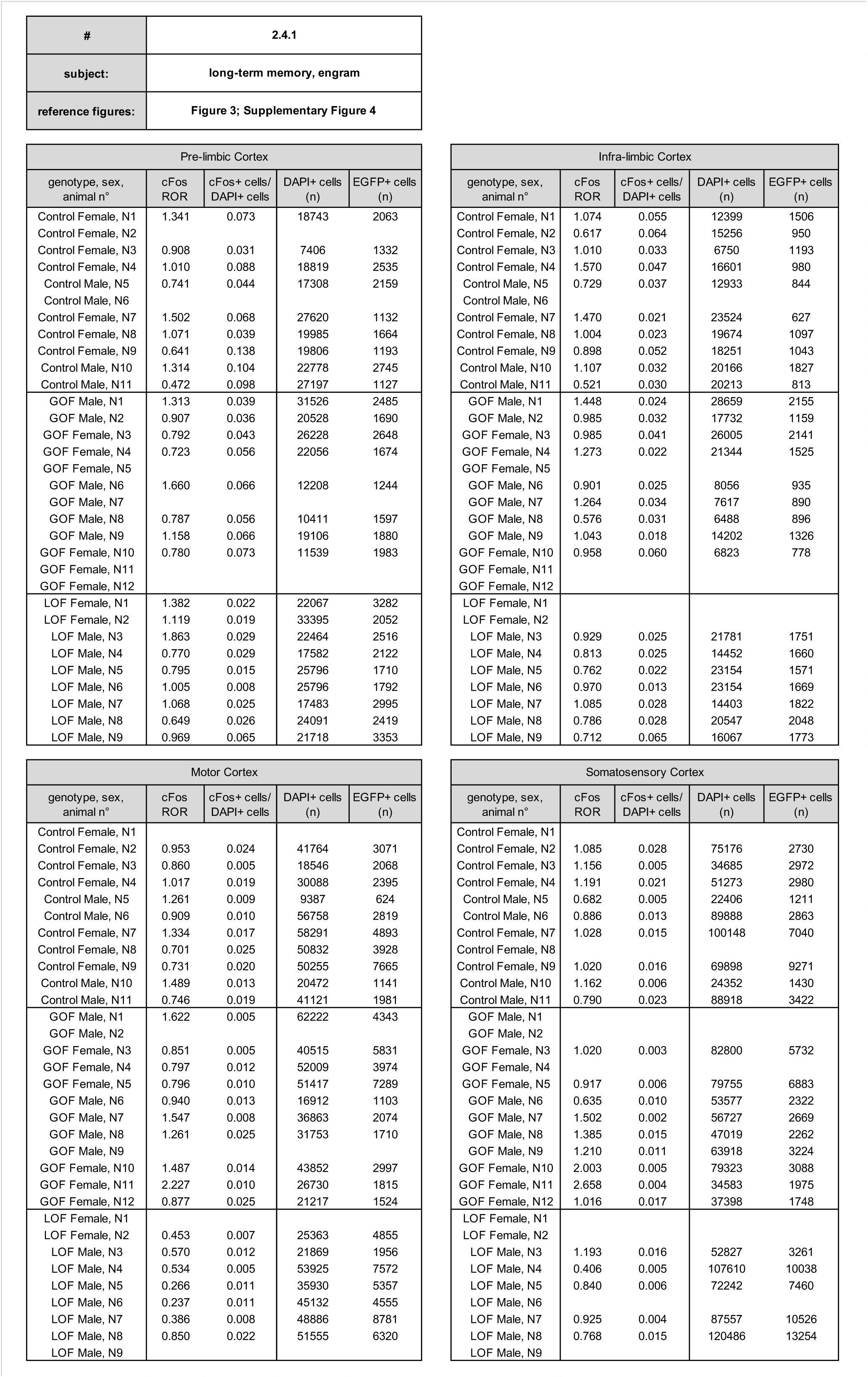
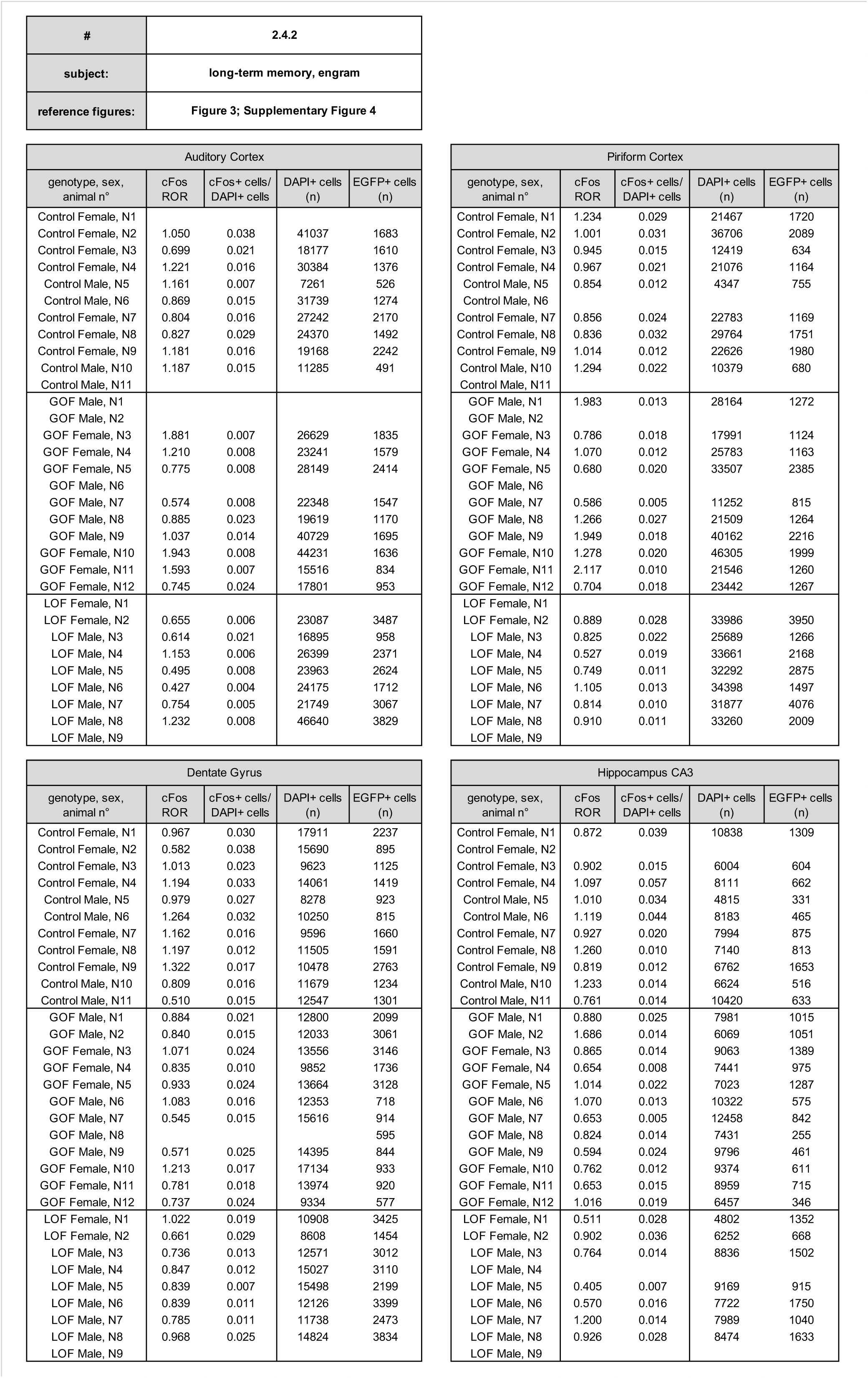
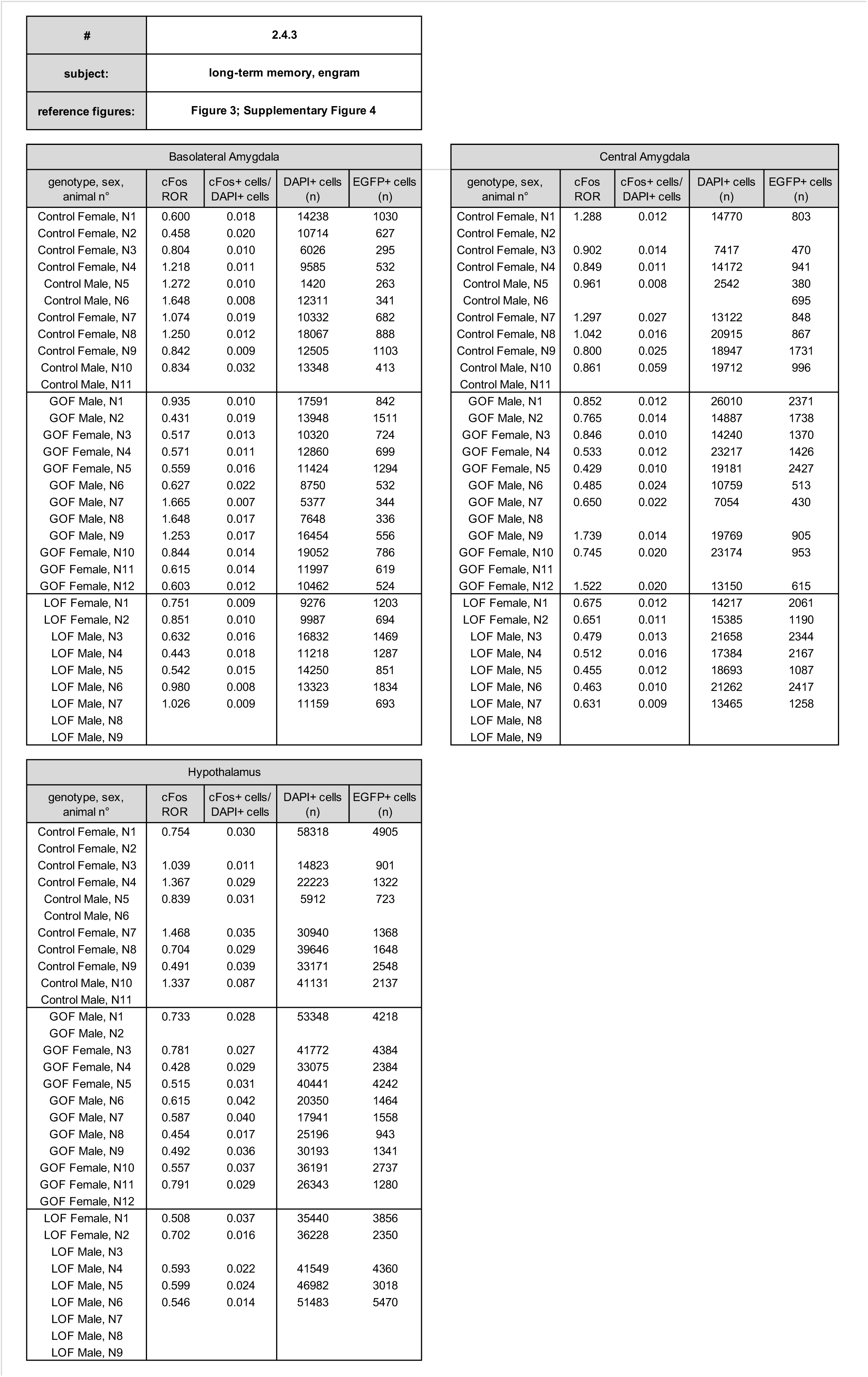
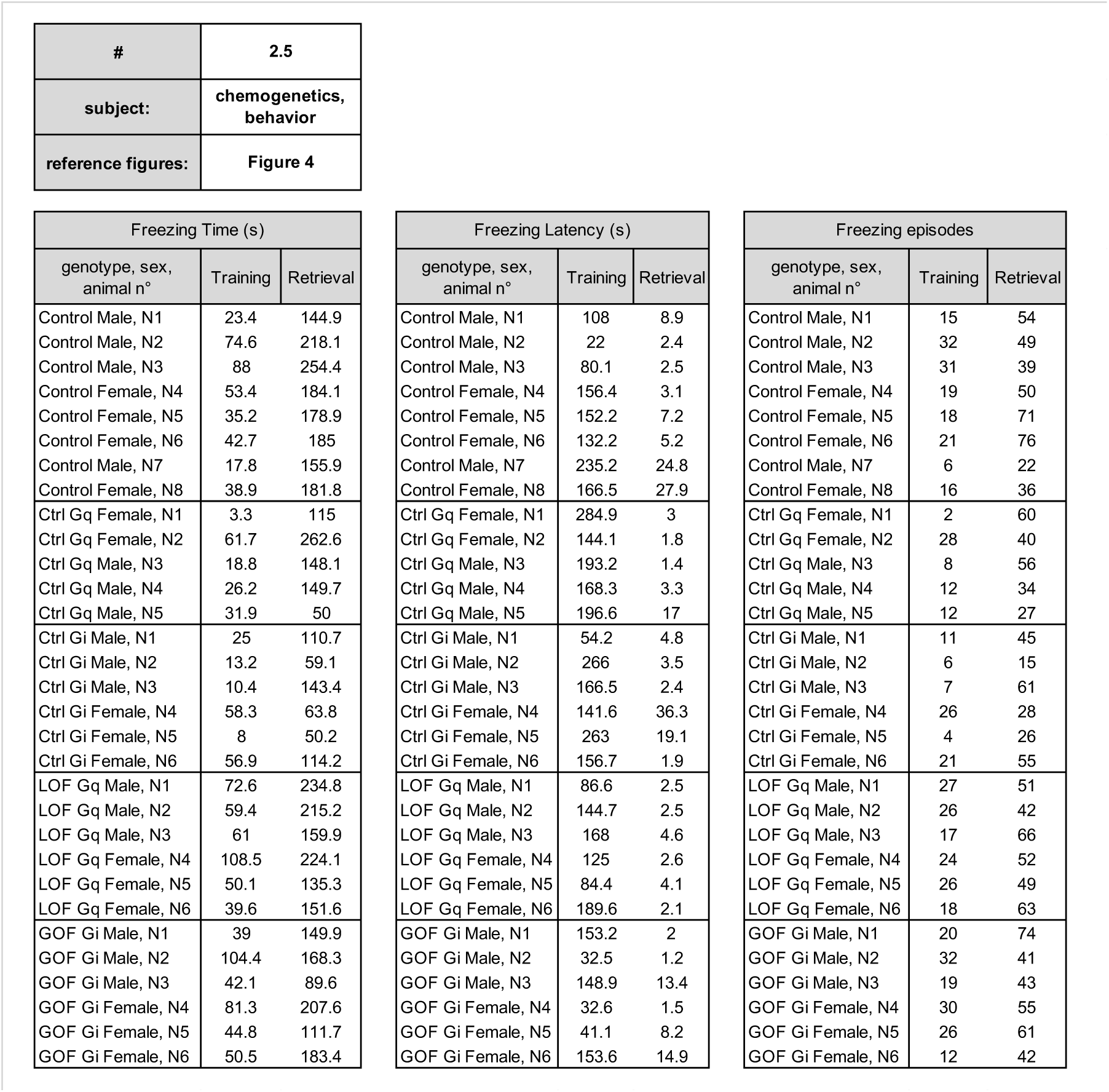
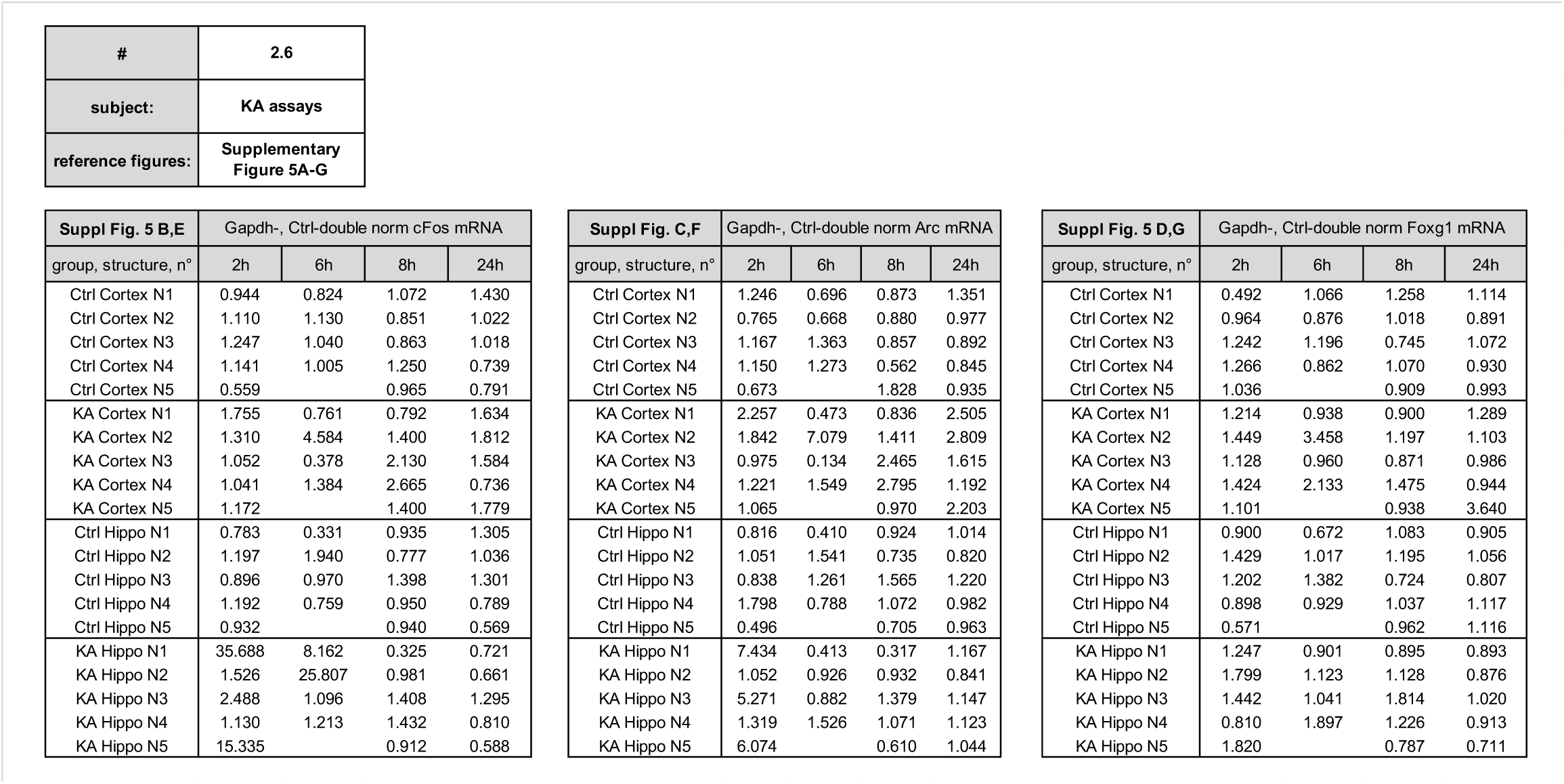
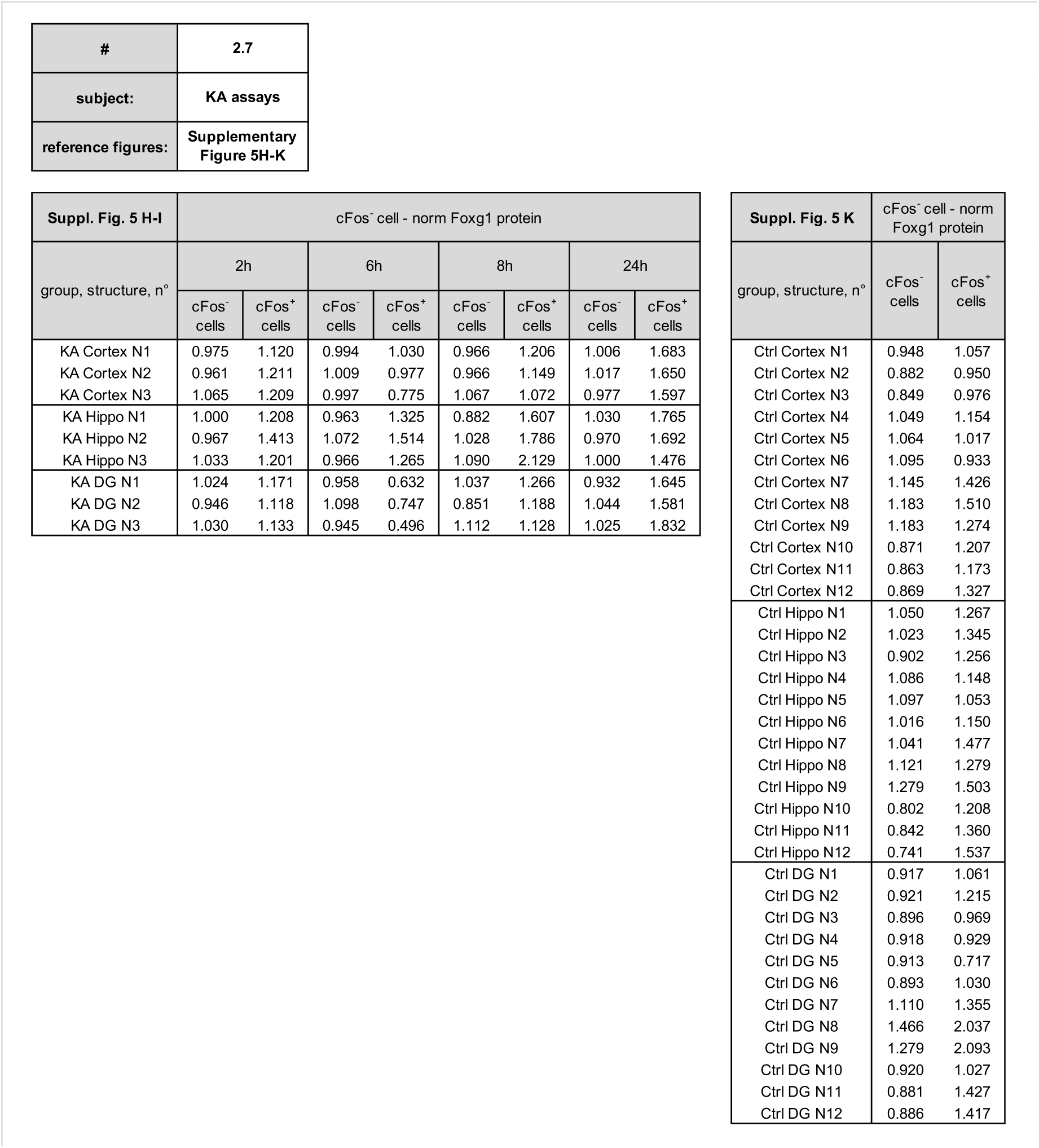
Primary data.

| # | subject | reference figure(s) |
| --- | --- | --- |
| 2.1 | short-term memory, behavior | Figure 1; Supplementary Figure 1 |
| 2.2 | short-term memory, engram | Figure 1,2, Supplementary Figure 2,3 |
| 2.3 | long-term memory, behavior | Figure 3; Supplementary Figure 4 |
| 2.4 | long-term memory, engram | Figure 3; Supplementary Figure 4 |
| 2.5 | chemogenetics, behavior | Figure 4 |
| 2.6 | KA assays, cFos-, Arc- and Foxg1-mRNA | Supplementary Figure 5A-G |
| 2.7 | KA assays, Foxg1 protein | Supplementary Figure 5H-K |

|  |  |
| --- | --- |
| # | 2.1 |
| subject: | short-term memory, behavior |
| reference figures: | Figure 1;<br>Supplementary Figure 1 |

| Freezing Time (s) |  |  |  |  |
| --- | --- | --- | --- | --- |
| genotype, sex, animal n° | Training (Ctx A) | Retrieval (Ctx A) | Retrieval (Ctx B) | Discrim. Index |
| Control Female, N1 | 71.5 | 42.4 | 59.8 | 0.7 |
| Control Female, N2 | 17.9 | 106.4 | 76.8 | 1.4 |
| Control Male, N3 | 11.7 | 79.9 | 62.4 | 1.3 |
| Control Male, N4 | 19.0 | 49.3 | 68.7 | 0.7 |
| Control Male, N5 | 85.4 | 168.3 | 167.5 | 1.0 |
| Control Male, N6 | 42.7 | 126.4 | 126.8 | 1.0 |
| Control Male, N7 | 35.7 | 76.1 | 15.5 | 4.9 |
| Control Female, N8 | 36.1 | 49.7 | 17.2 | 2.9 |
| Control Female, N9 | 88.6 | 216.9 | 35.8 | 6.1 |
| Control Male, N10 | 55.1 | 162.3 | 59.4 | 2.7 |
| Control Male, N11 | 49.2 | 110.4 | 104.3 | 1.1 |
| Control Male, N12 | 81.3 | 116.4 | 50.7 | 2.3 |
| Control Male, N13 | 56.3 | 209.7 | 49.7 | 4.2 |
| Control Female, N14 | 34.0 | 184.8 | 62.4 | 3.0 |
| Control Female, N15 | 66.8 | 179.8 | 93.9 | 1.9 |
| Control Female, N16 | 32.8 | 144.1 | 18.2 | 7.9 |
| Control Female, N17 | 85.2 | 197.9 | 148.2 | 1.3 |
| Control Female, N18 | 66.1 | 138.6 | 114.0 | 1.2 |
| Control Female, N19 | 51.7 | 117.9 | 59.3 | 2.0 |
| GOF Female, N1 | 34.7 | 154.5 | 97.9 | 1.6 |
| GOF Female, N2 | 21.2 | 139.8 | 51.9 | 2.7 |
| GOF Female, N3 | 74.3 | 141.2 | 162.6 | 0.9 |
| GOF Male, N4 | 48.3 | 92.8 | 68.8 | 1.3 |
| GOF Male, N5 | 31.0 | 177.6 | 182.8 | 1.0 |
| GOF Male, N6 | 86.8 | 177.5 | 20.8 | 8.5 |
| GOF Male, N7 | 69.3 | 223.1 | 75.5 | 3.0 |
| GOF Female, N8 | 40.5 | 141.2 | 68.5 | 2.1 |
| GOF Female, N9 | 61.0 | 185.8 | 97.3 | 1.9 |
| GOF Male, N10 | 61.8 | 249.5 | 114.6 | 2.2 |
| GOF Male, N11 | 82.3 | 172.3 | 27.2 | 6.3 |
| LOF Male, N1 | 53.1 | 184.5 | 54.6 | 3.4 |
| LOF Male, N2 | 76.3 | 142.1 | 62.7 | 2.3 |
| LOF Male, N3 | 40.4 | 196.5 | 174.7 | 1.1 |
| LOF Female, N4 | 22.7 | 129.8 | 95.6 | 1.4 |
| LOF Female, N5 | 6.1 | 62.3 | 25.0 | 2.5 |
| LOF Female, N6 | 32.0 | 161.1 | 98.7 | 1.6 |
| LOF Female, N7 | 64.4 | 105.4 | 117.2 | 0.9 |
| LOF Female, N8 | 53.8 | 104.4 | 180.1 | 0.6 |
| LOF Female, N9 | 42.4 | 90.7 | 49.8 | 1.8 |

| Freezing Latency (s) |  |  |  |
| --- | --- | --- | --- |
| genotype, sex, animal n° | Training (Ctx A) | Retrieval (Ctx A) | Retrieval (Ctx B) |
| Control Female, N1 | 7.9 | 18.7 | 50.7 |
| Control Female, N2 | 166.9 | 45.8 | 12.7 |
| Control Male, N3 | 199.5 | 13.0 | 15.5 |
| Control Male, N4 | 13.0 | 36.9 | 3.8 |
| Control Male, N5 | 51.4 | 27.2 | 36.7 |
| Control Male, N6 | 23.7 | 3.3 | 12.7 |
| Control Male, N7 | 127.0 | 5.8 | 43.9 |
| Control Female, N8 | 142.0 | 2.3 | 73.2 |
| Control Female, N9 | 134.2 | 1.3 | 53.9 |
| Control Male, N10 | 61.6 | 2.4 | 4.0 |
| Control Male, N11 | 198.7 | 2.6 | 7.3 |
| Control Male, N12 | 101.1 | 5.1 | 73.3 |
| Control Male, N13 | 90.8 | 3.2 | 26.7 |
| Control Female, N14 | 206.0 | 3.1 | 28.5 |
| Control Female, N15 | 140.8 | 3.6 | 14.7 |
| Control Female, N16 | 84.7 | 5.5 | 5.0 |
| Control Female, N17 | 38.3 | 2.7 | 12.8 |
| Control Female, N18 | 154.2 | 3.1 | 13.7 |
| Control Female, N19 | 138.5 | 8.5 | 9.1 |
| GOF Female, N1 | 134.9 | 9.6 | 3.1 |
| GOF Female, N2 | 205.8 | 42.2 | 4.6 |
| GOF Female, N3 | 135.0 | 1.4 | 6.4 |
| GOF Male, N4 | 22.8 | 3.9 | 11.6 |
| GOF Male, N5 | 142.4 | 2.0 | 11.8 |
| GOF Male, N6 | 130.5 | 1.8 | 8.1 |
| GOF Male, N7 | 147.4 | 1.2 | 5.4 |
| GOF Female, N8 | 38.0 | 2.0 | 7.1 |
| GOF Female, N9 | 188.6 | 4.4 | 13.8 |
| GOF Male, N10 | 138.5 | 1.9 | 10.6 |
| GOF Male, N11 | 143.0 | 3.0 | 32.8 |
| LOF Male, N1 | 147.1 | 3.8 | 30.0 |
| LOF Male, N2 | 80.7 | 3.6 | 11.5 |
| LOF Male, N3 | 151.7 | 5.0 | 1.6 |
| LOF Female, N4 | 161.2 | 15.5 | 11.3 |
| LOF Female, N5 | 260.4 | 46.5 | 93.5 |
| LOF Female, N6 | 160.5 | 10.9 | 8.9 |
| LOF Female, N7 | 155.9 | 2.9 | 9.6 |
| LOF Female, N8 | 187.2 | 2.8 | 8.6 |
| LOF Female, N9 | 143.1 | 6.7 | 14.5 |

| Freezing episodes |  |  |  |
| --- | --- | --- | --- |
| genotype, sex, animal n° | Training (Ctx A) | Retrieval (Ctx A) | Retrieval (Ctx B) |
| Control Female, N1 | 36 | 29 | 22 |
| Control Female, N2 | 7 | 38 | 47 |
| Control Male, N3 | 8 | 32 | 35 |
| Control Male, N4 | 11 | 40 | 33 |
| Control Male, N5 | 29 | 47 | 16 |
| Control Male, N6 | 25 | 58 | 54 |
| Control Male, N7 | 21 | 53 | 10 |
| Control Female, N8 | 21 | 28 | 11 |
| Control Female, N9 | 27 | 62 | 24 |
| Control Male, N10 | 21 | 51 | 25 |
| Control Male, N11 | 16 | 51 | 36 |
| Control Male, N12 | 26 | 53 | 24 |
| Control Male, N13 | 27 | 48 | 24 |
| Control Female, N14 | 12 | 47 | 26 |
| Control Female, N15 | 20 | 31 | 45 |
| Control Female, N16 | 14 | 64 | 10 |
| Control Female, N17 | 36 | 41 | 35 |
| Control Female, N18 | 28 | 35 | 47 |
| Control Female, N19 | 25 | 50 | 31 |
| GOF Female, N1 | 17 | 50 | 49 |
| GOF Female, N2 | 12 | 33 | 52 |
| GOF Female, N3 | 23 | 55 | 31 |
| GOF Male, N4 | 27 | 38 | 41 |
| GOF Male, N5 | 16 | 42 | 39 |
| GOF Male, N6 | 28 | 53 | 15 |
| GOF Male, N7 | 26 | 51 | 43 |
| GOF Female, N8 | 20 | 62 | 42 |
| GOF Female, N9 | 27 | 80 | 52 |
| GOF Male, N10 | 18 | 31 | 47 |
| GOF Male, N11 | 21 | 56 | 19 |
| LOF Male, N1 | 27 | 60 | 31 |
| LOF Male, N2 | 28 | 46 | 30 |
| LOF Male, N3 | 15 | 48 | 44 |
| LOF Female, N4 | 12 | 44 | 42 |
| LOF Female, N5 | 4 | 39 | 15 |
| LOF Female, N6 | 18 | 54 | 34 |
| LOF Female, N7 | 32 | 62 | 56 |
| LOF Female, N8 | 20 | 56 | 76 |
| LOF Female, N9 | 25 | 53 | 26 |

| # | 2.2.1 |
| --- | --- |
| subject: | short-term memory, engram |
| reference figures: | Figure 1,2;<br>Supplementary Figure 2,3 |

| Pre-limbic Cortex |  |  |  |  |  |  |  |
| --- | --- | --- | --- | --- | --- | --- | --- |
| genotype, sex,<br>animal n° | TdT-ROR | cFos-ROR | TdT+ cells/<br>DAPI+ cells | cFos+ cells/<br>DAPI+ cells | TdT+cFos+<br>cells/<br>cFos+ cells | DAPI+<br>cells | EGFP+<br>cells |
| Control Male, N1 | 1.5174 | 1.4162 | 0.0058 | 0.0202 | 0.0235 | 21012 | 150 |
| Control Female, N2 | 0.5684 | 0.9068 | 0.0033 | 0.0296 | 0.0162 | 37524 | 519 |
| Control Female, N3 | 0.7159 | 0.7257 | 0.0106 | 0.0349 | 0.0094 | 30370 | 361 |
| Control Male, N4 | 1.6721 | 0.6714 | 0.0045 | 0.0196 | 0.0394 | 36202 | 472 |
| Control Female, N5 |  |  |  |  |  |  |  |
| Control Female, N6 | 0.5261 | 1.2798 | 0.0047 | 0.0085 | 0.0687 | 58114 | 1005 |
| Control Male, N7 |  |  |  |  |  |  |  |
| Control Male, N8 |  |  |  |  |  |  |  |
| Control Male, N9 |  |  |  |  |  |  |  |
| Control Male, N10 |  |  |  |  |  |  |  |
| Control Male, N11 |  | 1.3638 |  | 0.0158 |  | 32392 | 888 |
| Control Male, N12 |  | 0.8633 |  | 0.0513 |  | 19097 | 1232 |
| Control Male, N13 | 0.7854 | 1.0781 | 0.0100 | 0.0336 | 0.0180 | 32998 | 2948 |
| Control Female, N14 | 1.2146 | 0.9101 | 0.0097 | 0.0614 | 0.0131 | 32410 | 2729 |
| Control Female, N15 |  | 0.7847 |  | 0.0181 |  | 30703 | 4536 |
| Control Female, N16 |  |  |  |  |  |  |  |
| Control Female, N17 |  |  |  |  |  |  |  |
| Control Female, N18 |  |  |  |  |  |  |  |
| Control Female, N19 |  |  |  |  |  |  |  |
| GOF Male, N1 | 4.8254 | 2.0243 | 0.24% | 1.02% | 5.37% | 14599 | 276 |
| GOF Male, N2 | 0.4958 | 2.2759 | 0.38% | 0.94% | 7.74% | 16574 | 795 |
| GOF Female, N3 | 7.4928 | 2.5576 | 0.15% | 1.25% | 4.22% | 18985 | 290 |
| GOF Female, N4 |  | 1.0165 |  | 2.79% |  | 21862 | 460 |
| GOF Male, N5 | 1.9019 | 2.3052 | 0.62% | 1.06% | 7.78% | 25356 | 373 |
| GOF Male, N6 | 4.2520 | 2.7289 | 0.14% | 0.77% | 3.81% | 27205 | 515 |
| GOF Female, N7 |  |  |  |  |  |  |  |
| GOF Female, N8 |  |  |  |  |  |  |  |
| GOF Female, N9 |  |  |  |  |  |  |  |
| GOF Male, N10 |  |  |  |  |  |  |  |
| GOF Male, N11 |  |  |  |  |  |  |  |
| LOF Male, N1 |  | 0.7293 |  | 1.10% |  | 33832 | 1006 |
| LOF Male, N2 | 1.1196 | 0.5025 | 0.25% | 2.15% | 1.60% | 26136 | 742 |
| LOF Male, N3 |  | 0.4688 |  | 1.19% |  | 29751 | 441 |
| LOF Female, N4 | 0.8722 | 0.9281 | 0.66% | 3.84% | 2.34% | 27853 | 1224 |
| LOF Female, N5 | 0.8996 | 0.6945 | 0.73% | 0.78% | 5.93% | 34492 | 1718 |
| LOF Female, N6 |  |  |  |  |  |  |  |
| LOF Female, N7 |  |  |  |  |  |  |  |
| LOF Female, N8 |  |  |  |  |  |  |  |
| LOF Female, N9 |  |  |  |  |  |  |  |

| Motor Cortex |  |  |  |  |  |  |  |
| --- | --- | --- | --- | --- | --- | --- | --- |
| genotype, sex,<br>animal n° | TdT-ROR | cFos-ROR | TdT+ cells/<br>DAPI+ cells | cFos+ cells/<br>DAPI+ cells | TdT+cFos+<br>cells/<br>cFos+ cells | DAPI+<br>cells | EGFP+<br>cells |
| Control Male, N1 | 1.0654 | 1.0580 | 0.0129 | 0.0066 | 0.0230 | 13119 | 334 |
| Control Female, N2 | 0.9198 | 1.1866 | 0.0076 | 0.0099 | 0.0400 | 27795 | 651 |
| Control Female, N3 | 1.0942 | 0.9714 | 0.0184 | 0.0138 | 0.0593 | 19573 | 1066 |
| Control Male, N4 | 1.5558 | 0.7093 | 0.0091 | 0.0179 | 0.0136 | 16413 | 776 |
| Control Female, N5 | 0.5473 | 1.0746 | 0.0014 | 0.0068 | 0.0320 | 18439 | 989 |
| Control Female, N6 | 0.1449 |  | 0.0136 |  |  | 11130 | 376 |
| Control Male, N7 | 1.5327 |  | 0.0046 |  |  | 21773 | 887 |
| Control Male, N8 |  |  |  |  |  |  |  |
| Control Male, N9 | 1.1400 |  | 0.0130 |  |  | 11132 | 309 |
| Control Male, N10 |  |  |  |  |  |  |  |
| Control Male, N11 |  | 1.1802 |  | 0.0156 |  | 18271 | 2146 |
| Control Male, N12 |  | 0.7584 |  | 0.0288 |  | 23123 | 6483 |
| Control Male, N13 |  | 1.1107 |  | 0.0431 |  | 17481 | 2475 |
| Control Female, N14 | 1.0701 | 0.8535 | 0.0131 | 0.0026 | 0.0200 | 19483 | 7997 |
| Control Female, N15 | 1.0418 | 1.0972 | 0.0071 | 0.0456 | 0.0022 | 19493 | 7187 |
| Control Female, N16 | 0.5634 |  | 0.0149 |  | 0.0047 | 20223 | 9665 |
| Control Female, N17 | 1.3247 |  | 0.0053 |  | 0.0068 | 14992 | 1848 |
| Control Female, N18 |  |  |  |  |  |  |  |
| Control Female, N19 |  |  |  |  |  |  |  |
| GOF Male, N1 | 0.0000 | 2.4520 | 0.0097 | 0.0169 | 0.0519 | 7986 | 169 |
| GOF Male, N2 | 0.0000 | 1.0418 | 0.0026 | 0.0078 | 0.0000 | 20385 | 283 |
| GOF Female, N3 | 0.2760 | 1.6259 | 0.0138 | 0.0085 | 0.0176 | 19909 | 198 |
| GOF Female, N4 | 3.2931 | 0.7655 | 0.0031 | 0.0090 | 0.0364 | 24529 | 1081 |
| GOF Male, N5 |  | 1.4774 |  | 0.0061 |  | 19889 | 737 |
| GOF Male, N6 | 0.5365 | 3.2126 | 0.0084 | 0.0091 | 0.0490 | 22443 | 666 |
| GOF Female, N7 | 1.1828 |  | 0.0173 |  |  | 19017 | 635 |
| GOF Female, N8 | 0.7667 |  | 0.0171 |  |  | 18193 | 1203 |
| GOF Female, N9 | 4.4676 | 0.6860 | 0.0011 | 0.0038 | 0.0253 | 21004 | 317 |
| GOF Male, N10 | 0.0000 |  | 0.0013 |  |  | 24893 | 205 |
| GOF Male, N11 |  |  |  |  |  |  |  |
| LOF Male, N1 | 2.7774 | 0.8055 | 0.0047 | 0.0133 | 0.0159 | 18908 | 4215 |
| LOF Male, N2 | 1.0446 | 0.7598 | 0.0166 | 0.0097 | 0.0000 | 17783 | 2640 |
| LOF Male, N3 | 0.7785 | 0.9296 | 0.0038 | 0.0338 | 0.0056 | 15979 | 1414 |
| LOF Female, N4 |  | 0.6825 |  | 0.0342 |  | 17251 | 3274 |
| LOF Female, N5 | 1.6879 |  | 0.0076 |  | 0.0069 | 21844 | 2415 |
| LOF Female, N6 | 1.2043 |  | 0.0324 |  | 0.0226 | 26229 | 2423 |
| LOF Female, N7 | 1.8159 | 1.8905 | 0.0054 | 0.0350 | 0.0060 | 23884 | 2251 |
| LOF Female, N8 |  |  |  |  |  |  |  |
| LOF Female, N9 |  |  |  |  |  |  |  |

| Infra-limbic Cortex |  |  |  |  |  |  |  |
| --- | --- | --- | --- | --- | --- | --- | --- |
| genotype, sex,<br>animal n° | TdT-ROR | cFos-ROR | TdT+ cells/<br>DAPI+ cells | cFos+ cells/<br>DAPI+ cells | TdT+cFos+<br>cells/<br>cFos+ cells | DAPI+<br>cells | EGFP+<br>cells |
| Control Male, N1 | 0.0000 | 1.2748 | 0.0029 | 0.0088 | 0.1143 | 3993 | 67 |
| Control Female, N2 | 1.4015 |  | 0.0034 |  | 0.0450 | 14541 | 457 |
| Control Female, N3 | 0.9950 | 0.5572 | 0.0073 | 0.0129 | 0.0829 | 15934 | 437 |
| Control Male, N4 | 1.5676 | 0.4992 | 0.0015 | 0.0176 | 0.0214 | 15873 | 463 |
| Control Female, N5 |  | 1.0551 |  | 0.0080 | 0.0171 | 21998 | 743 |
| Control Female, N6 | 1.0359 | 1.6136 | 0.0048 | 0.0023 | 0.2586 | 24995 | 997 |
| Control Male, N7 |  |  |  |  |  |  |  |
| Control Male, N8 |  |  |  |  |  |  |  |
| Control Male, N9 |  |  |  |  |  |  |  |
| Control Male, N10 |  |  |  |  |  |  |  |
| Control Male, N11 |  | 1.5445 |  | 0.0087 |  | 23234 | 718 |
| Control Male, N12 | 1.5853 | 0.6657 | 0.0009 | 0.0275 | 0.0032 | 11438 | 782 |
| Control Male, N13 | 0.6452 | 0.8021 | 0.0061 | 0.0295 | 0.0211 | 25751 | 2437 |
| Control Female, N14 | 0.7696 | 1.0401 | 0.0050 | 0.0205 | 0.0302 | 20982 | 1768 |
| Control Female, N15 |  | 0.9476 |  | 0.0164 |  | 24969 | 3473 |
| Control Female, N16 |  |  |  |  |  |  |  |
| Control Female, N17 |  |  |  |  |  |  |  |
| Control Female, N18 |  |  |  |  |  |  |  |
| Control Female, N19 |  |  |  |  |  |  |  |
| GOF Male, N1 | 2.5362 | 1.0805 | 0.0029 | 0.0109 | 0.0720 | 11436 | 255 |
| GOF Male, N2 | 2.8456 | 0.9117 | 0.0016 | 0.0122 | 0.0276 | 14872 | 884 |
| GOF Female, N3 |  | 1.9529 |  | 0.0083 |  | 18263 | 323 |
| GOF Female, N4 |  | 0.7842 |  | 0.0230 |  | 19350 | 448 |
| GOF Male, N5 | 9.3715 | 2.1698 | 0.0029 | 0.0075 | 0.0551 | 16848 | 203 |
| GOF Male, N6 | 6.3465 | 1.4333 | 0.0017 | 0.0084 | 0.0588 | 20235 | 404 |
| GOF Female, N7 |  |  |  |  |  |  |  |
| GOF Female, N8 |  |  |  |  |  |  |  |
| GOF Female, N9 |  |  |  |  |  |  |  |
| GOF Male, N10 |  |  |  |  |  |  |  |
| GOF Male, N11 |  |  |  |  |  |  |  |
| LOF Male, N1 |  | 1.5580 |  | 0.0107 |  | 26201 | 599 |
| LOF Male, N2 | 1.8592 | 0.7286 | 0.0035 | 0.0123 | 0.0170 | 19071 | 451 |
| LOF Male, N3 |  | 0.6103 |  | 0.0095 |  | 13216 | 174 |
| LOF Female, N4 | 1.1410 | 0.8420 | 0.0078 | 0.0309 | 0.0362 | 22320 | 991 |
| LOF Female, N5 | 1.1241 | 1.3206 | 0.0102 | 0.0070 | 0.0595 | 26590 | 1106 |
| LOF Female, N6 |  |  |  |  |  |  |  |
| LOF Female, N7 |  |  |  |  |  |  |  |
| LOF Female, N8 |  |  |  |  |  |  |  |
| LOF Female, N9 |  |  |  |  |  |  |  |

| Somatosensory Cortex |  |  |  |  |  |  |  |
| --- | --- | --- | --- | --- | --- | --- | --- |
| genotype, sex,<br>animal n° | TdT-ROR | cFos-ROR | TdT+ cells/<br>DAPI+ cells | cFos+ cells/<br>DAPI+ cells | TdT+cFos+<br>cells/<br>cFos+ cells | DAPI+<br>cells | EGFP+<br>cells |
| Control Male, N1 | 0.7192 | 0.6144 | 0.0116 | 0.0101 | 0.0054 | 36337 | 942 |
| Control Female, N2 | 0.5230 | 1.7855 | 0.0067 | 0.0025 | 0.0601 | 73553 | 1497 |
| Control Female, N3 | 0.9047 | 0.3620 | 0.0204 | 0.0092 | 0.0349 | 46952 | 2739 |
| Control Male, N4 | 2.1363 | 1.2382 | 0.0047 | 0.0142 | 0.0250 | 28081 | 626 |
| Control Female, N5 |  |  |  |  |  |  |  |
| Control Female, N6 |  |  |  |  |  |  |  |
| Control Male, N7 | 0.9370 |  | 0.0088 |  |  | 62307 | 3148 |
| Control Male, N8 |  |  |  |  |  |  |  |
| Control Male, N9 | 0.7798 |  | 0.0074 |  |  | 30441 | 903 |
| Control Male, N10 |  |  |  |  |  |  |  |
| Control Male, N11 |  | 0.8685 |  | 0.0081 |  | 41501 | 4416 |
| Control Male, N12 |  | 0.5445 |  | 0.0167 |  | 45324 | 11712 |
| Control Male, N13 |  | 0.5284 |  | 0.0527 |  | 38062 | 6751 |
| Control Female, N14 | 0.9342 | 1.2475 | 0.0212 | 0.0021 | 0.0182 | 51292 | 14165 |
| Control Female, N15 | 0.7580 | 0.3980 | 0.0132 | 0.0446 | 0.0012 | 38080 | 14449 |
| Control Female, N16 | 1.1501 | 2.4131 | 0.0170 | 0.0442 | 0.0050 | 54336 | 18356 |
| Control Female, N17 | 1.1577 |  | 0.0079 |  | 0.0071 | 55551 | 12945 |
| Control Female, N18 |  |  |  |  |  |  |  |
| Control Female, N19 |  |  |  |  |  |  |  |
| GOF Male, N1 | 2.1240 | 0.9972 | 0.0043 | 0.0038 | 0.0471 | 44294 | 592 |
| GOF Male, N2 | 0.0000 | 2.8497 | 0.0033 | 0.0046 | 0.0099 | 44169 | 536 |
| GOF Female, N3 | 0.6317 | 0.8055 | 0.0107 | 0.0036 | 0.0564 | 54360 | 393 |
| GOF Female, N4 | 1.5834 | 0.3832 | 0.0029 | 0.0078 | 0.0301 | 46920 | 2962 |
| GOF Male, N5 |  | 2.2296 | 0.0034 | 0.0020 |  | 57337 | 2193 |
| GOF Male, N6 | 0.5351 | 2.3943 | 0.0083 | 0.0064 | 0.0330 | 56768 | 1570 |
| GOF Female, N7 |  |  |  |  |  |  |  |
| GOF Female, N8 | 0.0000 |  | 0.0016 |  |  | 58312 | 413 |
| GOF Female, N9 | 0.0000 |  | 0.0007 |  |  | 52809 | 550 |
| GOF Male, N10 | 0.7552 |  | 0.0169 |  |  | 50167 | 2514 |
| GOF Male, N11 | 0.6597 |  | 0.0154 |  |  | 49930 | 2554 |
| LOF Male, N1 |  | 0.4836 |  | 0.0064 |  | 60495 | 6047 |
| LOF Male, N2 | 1.3100 | 1.5254 | 0.0116 | 0.0026 | 0.0104 | 74611 | 4392 |
| LOF Male, N3 |  | 0.3850 |  | 0.0060 |  | 59980 | 2516 |
| LOF Female, N4 |  | 0.7031 |  | 0.0155 |  | 68504 | 5025 |
| LOF Female, N5 | 4.0946 |  | 0.0079 |  | 0.0110 | 66592 | 3403 |
| LOF Female, N6 | 1.9643 | 1.8086 | 0.0295 | 0.0127 | 0.0136 | 69350 | 3538 |
| LOF Female, N7 | 2.3604 | 1.6261 | 0.0054 | 0.0234 | 0.0077 | 61214 | 4439 |
| LOF Female, N8 |  |  |  |  |  |  |  |
| LOF Female, N9 |  |  |  |  |  |  |  |

| # | 2.2.2 |
| --- | --- |
| subject: | short-term memory, engram |
| reference figures: | Figure 1,2;<br>Supplementary Figure 2,3 |

| Auditory Cortex |  |  |  |  |  |  |  |
| --- | --- | --- | --- | --- | --- | --- | --- |
| genotype, sex, animal n° | TdT-ROR | cFos-ROR | TdT+ cells/<br>DAPI+ cells | cFos+ cells/<br>DAPI+ cells | TdT+cFos+ cells/<br>cFos+ cells | DAPI+ cells | EGFP+ cells |
| Control Male, N1 | 0.9231 | 1.7446 | 0.0092 | 0.0019 | 0.0000 | 11528 | 270 |
| Control Female, N2 | 0.4090 | 0.5446 | 0.0061 | 0.0049 | 0.0091 | 22589 | 304 |
| Control Female, N3 | 1.2194 | 0.6301 | 0.0168 | 0.0055 | 0.0792 | 18492 | 904 |
| Control Male, N4 | 0.0000 | 0.1121 | 0.0218 | 0.0522 | 0.0308 | 3734 | 343 |
| Control Female, N5 |  | 1.9685 |  | 0.0015 | 0.0296 | 87106 | 542 |
| Control Female, N6 | 2.7630 |  | 0.0076 |  |  | 12419 | 464 |
| Control Male, N7 | 0.9410 |  | 0.0063 |  |  | 18345 | 1021 |
| Control Male, N8 |  |  |  |  |  |  |  |
| Control Male, N9 | 0.7443 |  | 0.0152 |  |  | 10148 | 202 |
| Control Male, N10 |  |  |  |  |  |  |  |
| Control Male, N11 |  |  |  |  |  |  |  |
| Control Male, N12 |  | 0.5870 |  | 0.0058 |  | 23631 | 3435 |
| Control Male, N13 |  | 0.9174 |  | 0.0150 |  | 31593 | 2176 |
| Control Female, N14 |  | 0.4387 |  | 0.0030 |  | 35037 | 3812 |
| Control Female, N15 | 1.3812 | 0.9499 | 0.0072 | 0.0064 | 0.0000 | 32877 | 2856 |
| Control Female, N16 | 1.2378 | 1.4065 | 0.0119 | 0.0621 | 0.0066 | 23449 | 9903 |
| Control Female, N17 | 0.3810 | 1.7005 | 0.0170 | 0.0424 | 0.0058 | 27901 | 7130 |
| Control Female, N18 |  |  |  |  |  |  |  |
| Control Female, N19 |  |  |  |  |  |  |  |
| GOF Male, N1 | 0.0000 | 2.0703 | 0.0141 | 0.0027 | 0.0870 | 17354 | 209 |
| GOF Male, N2 | 0.0000 | 0.5844 | 0.0162 | 0.0023 | 0.0000 | 16520 | 200 |
| GOF Female, N3 | 1.7347 | 1.7102 | 0.0007 | 0.0051 | 0.0337 | 17299 | 93 |
| GOF Female, N4 | 1.9623 | 0.7054 | 0.0022 | 0.0059 | 0.0246 | 20625 | 912 |
| GOF Male, N5 | 0.0000 | 0.5592 | 0.0034 | 0.0012 | 0.0435 | 18615 | 643 |
| GOF Male, N6 | 1.0426 | 2.2642 | 0.0028 | 0.0031 | 0.0658 | 24707 | 472 |
| GOF Female, N7 | 0.8973 |  | 0.0095 |  |  | 14677 | 481 |
| GOF Female, N8 | 0.9486 |  | 0.0022 |  |  | 17860 | 943 |
| GOF Female, N9 | 0.0000 |  | 0.0032 |  |  | 20477 | 176 |
| GOF Male, N10 | 0.0000 |  | 0.0047 |  |  | 20511 | 134 |
| GOF Male, N11 |  |  |  |  |  |  |  |
| LOF Male, N1 |  | 0.9510 |  | 0.0028 |  | 33038 | 1927 |
| LOF Male, N2 | 1.1978 | 0.6001 | 0.0189 | 0.0025 | 0.0161 | 24826 | 969 |
| LOF Male, N3 |  |  |  |  |  | 36707 | 779 |
| LOF Female, N4 |  | 0.7856 |  | 0.0077 |  | 31352 | 2306 |
| LOF Female, N5 | 2.2945 | 1.0371 | 0.0110 | 0.0171 | 0.0157 | 26077 | 1412 |
| LOF Female, N6 | 0.8281 | 0.7708 | 0.0414 | 0.0113 | 0.0375 | 28353 | 1884 |
| LOF Female, N7 |  | 1.3187 |  | 0.0135 |  | 14487 | 1021 |
| LOF Female, N8 |  |  |  |  |  |  |  |
| LOF Female, N9 |  |  |  |  |  |  |  |

| Dentate Gyrus |  |  |  |  |  |  |  |
| --- | --- | --- | --- | --- | --- | --- | --- |
| genotype, sex, animal n° | TdT-ROR | cFos-ROR | TdT+ cells/<br>DAPI+ cells | cFos+ cells/<br>DAPI+ cells | TdT+cFos+ cells/<br>cFos+ cells | DAPI+ cells | EGFP+ cells |
| Control Male, N1 | 2.2365 |  | 0.0118 |  |  | 8282 | 363 |
| Control Female, N2 | 0.4965 | 1.7369 | 0.0176 | 0.0214 | 0.0065 | 7241 | 675 |
| Control Female, N3 | 0.8084 | 1.3582 | 0.0241 | 0.0196 | 0.0183 | 5568 | 713 |
| Control Male, N4 | 1.0385 |  | 0.0087 |  |  | 7633 | 763 |
| Control Female, N5 | 1.0154 | 0.7467 | 0.0049 | 0.0150 | 0.0000 | 6067 | 1178 |
| Control Female, N6 | 0.7726 | 0.8276 | 0.0189 | 0.0152 | 0.0000 | 5935 | 832 |
| Control Male, N7 | 0.8369 | 0.4353 | 0.0129 | 0.0193 | 0.0083 | 6254 | 1142 |
| Control Male, N8 | 1.3631 | 0.8953 | 0.0013 | 0.0298 | 0.0000 | 5399 | 1093 |
| Control Male, N9 | 0.4321 |  | 0.0259 |  |  | 4236 | 427 |
| Control Male, N10 |  |  |  |  |  |  |  |
| Control Male, N11 | 0.5566 | 0.9192 | 0.0027 | 0.0289 | 0.0000 | 11549 | 1719 |
| Control Male, N12 | 1.0426 | 1.8741 | 0.0025 | 0.0243 | 0.0000 | 15468 | 2940 |
| Control Male, N13 | 2.3302 | 1.8686 | 0.0067 | 0.0221 | 0.0044 | 10336 | 2925 |
| Control Female, N14 | 0.5827 |  | 0.0126 |  |  | 11706 | 2859 |
| Control Female, N15 | 1.3517 | 1.0852 | 0.0188 | 0.0373 | 0.0000 | 12078 | 3123 |
| Control Female, N16 | 0.4467 | 0.5165 | 0.0213 | 0.0444 | 0.0081 | 8360 | 2781 |
| Control Female, N17 | 0.6896 | 0.4746 | 0.0334 | 0.0363 | 0.0115 | 7186 | 2618 |
| Control Female, N18 |  | 0.4621 |  | 0.0456 |  | 5501 | 1827 |
| Control Female, N19 |  | 0.7996 |  | 0.0412 |  | 4850 | 1554 |
| GOF Male, N1 | 2.2444 |  | 0.0079 |  |  | 11718 | 652 |
| GOF Male, N2 |  | 1.2225 |  | 0.0208 |  | 6296 | 256 |
| GOF Female, N3 |  | 2.2936 |  | 0.0213 |  | 6445 | 136 |
| GOF Female, N4 | 2.2901 |  | 0.0055 |  |  | 10974 | 1234 |
| GOF Male, N5 | 1.2420 | 1.0622 | 0.0068 | 0.0318 | 0.0046 | 6795 | 479 |
| GOF Male, N6 | 2.2573 | 1.2189 | 0.0096 | 0.0245 | 0.0045 | 9048 | 509 |
| GOF Female, N7 | 0.7877 | 0.9855 | 0.0054 | 0.0250 | 0.0196 | 4079 | 733 |
| GOF Female, N8 | 0.8298 | 0.9158 | 0.0034 | 0.0185 | 0.0114 | 4746 | 684 |
| GOF Female, N9 | 0.9893 | 2.3149 | 0.0046 | 0.0119 | 0.0000 | 3032 | 167 |
| GOF Male, N10 |  | 1.7459 |  | 0.0191 | 0.0083 | 6343 | 232 |
| GOF Male, N11 |  | 1.4966 |  | 0.0303 |  | 4054 | 472 |
| LOF Male, N1 | 1.4375 | 1.1674 | 0.0038 | 0.0138 | 0.0000 | 15240 | 1720 |
| LOF Male, N2 | 1.1539 | 1.1782 | 0.0094 | 0.0275 | 0.0097 | 11195 | 1245 |
| LOF Male, N3 |  | 0.6007 |  | 0.0207 |  | 15086 | 773 |
| LOF Female, N4 | 0.6382 | 1.2742 | 0.0028 | 0.0292 | 0.0000 | 10960 | 1169 |
| LOF Female, N5 | 1.3432 | 0.8328 | 0.0097 | 0.0251 | 0.0101 | 11770 | 1598 |
| LOF Female, N6 | 0.3803 | 0.4436 | 0.0247 | 0.0649 | 0.0075 | 6195 | 1584 |
| LOF Female, N7 | 0.7440 | 0.4628 | 0.0505 | 0.0330 | 0.0224 | 4060 | 510 |
| LOF Female, N8 | 0.3702 | 0.5479 | 0.0204 | 0.0537 | 0.0100 | 5602 | 792 |
| LOF Female, N9 |  | 0.6545 |  | 0.0399 |  | 5316 | 799 |

| Piriform Cortex |  |  |  |  |  |  |  |
| --- | --- | --- | --- | --- | --- | --- | --- |
| genotype, sex, animal n° | TdT-ROR | cFos-ROR | TdT+ cells/<br>DAPI+ cells | cFos+ cells/<br>DAPI+ cells | TdT+cFos+ cells/<br>cFos+ cells | DAPI+ cells | EGFP+ cells |
| Control Male, N1 | 1.4718 | 0.9781 | 0.0061 | 0.0130 | 0.0270 | 17135 | 387 |
| Control Female, N2 | 1.3818 | 1.1774 | 0.0184 | 0.0096 | 0.0423 | 14833 | 440 |
| Control Female, N3 | 0.4798 | 0.4740 | 0.0173 | 0.0283 | 0.0155 | 15953 | 411 |
| Control Male, N4 | 1.4303 | 1.3952 | 0.0081 | 0.0150 | 0.0421 | 30013 | 301 |
| Control Female, N5 | 0.8998 | 0.9753 | 0.0016 | 0.0118 | 0.0536 | 23799 | 488 |
| Control Female, N6 | 1.4080 |  | 0.0098 |  |  | 19738 | 406 |
| Control Male, N7 | 0.9294 |  | 0.0095 |  |  | 19498 | 470 |
| Control Male, N8 | 0.0000 |  | 0.0008 |  |  | 13861 | 491 |
| Control Male, N9 | 0.9991 |  | 0.0183 |  |  | 22094 | 453 |
| Control Male, N10 |  |  |  |  |  |  |  |
| Control Male, N11 |  | 1.0131 |  | 0.0053 |  | 35134 | 1120 |
| Control Male, N12 | 1.0596 | 0.9126 | 0.0048 | 0.0149 | 0.0000 | 28067 | 2801 |
| Control Male, N13 |  | 1.8983 |  | 0.0326 |  | 27918 | 2115 |
| Control Female, N14 | 1.3707 | 0.0729 | 0.0100 | 0.0315 | 0.0000 | 26323 | 2146 |
| Control Female, N15 | 0.5744 | 1.1031 | 0.0167 | 0.0294 | 0.0084 | 28262 | 4932 |
| Control Female, N16 | 0.6311 |  | 0.0104 |  |  | 27462 | 6474 |
| Control Female, N17 | 1.3642 |  | 0.0080 |  |  | 24859 | 7006 |
| Control Female, N18 |  |  |  |  |  |  |  |
| Control Female, N19 |  |  |  |  |  |  |  |
| GOF Male, N1 | 0.0000 | 5.0681 | 0.0035 | 0.0306 | 0.0287 | 15894 | 177 |
| GOF Male, N2 | 1.9518 | 7.3034 | 0.0026 | 0.0041 | 0.0370 | 13119 | 99 |
| GOF Female, N3 | 0.0000 | 2.1704 | 0.0164 | 0.0140 | 0.0636 | 24729 | 127 |
| GOF Female, N4 | 1.8383 | 1.3921 | 0.0209 | 0.0150 | 0.0267 | 24922 | 611 |
| GOF Male, N5 | 0.9505 | 5.7163 | 0.0056 | 0.0050 | 0.0408 | 29187 | 736 |
| GOF Male, N6 | 1.6711 | 2.3132 | 0.0036 | 0.0057 | 0.0383 | 31850 | 351 |
| GOF Female, N7 |  |  |  |  |  | 25085 | 342 |
| GOF Female, N8 | 0.0000 |  | 0.0110 |  |  | 24553 | 112 |
| GOF Female, N9 | 0.0000 |  | 0.0063 |  |  | 26425 | 178 |
| GOF Male, N10 | 1.2130 |  | 0.0059 |  |  | 17191 | 561 |
| GOF Male, N11 | 0.8934 |  | 0.0059 |  |  | 16667 | 628 |
| LOF Male, N1 | 0.8372 | 0.9572 | 0.0041 | 0.0136 | 0.0054 | 27225 | 1604 |
| LOF Male, N2 | 0.4975 | 0.4506 | 0.0204 | 0.0077 | 0.0221 | 29222 | 1836 |
| LOF Male, N3 |  | 0.8573 |  | 0.0166 |  | 29456 | 627 |
| LOF Female, N4 |  | 1.3805 |  | 0.0115 |  | 31449 | 1256 |
| LOF Female, N5 | 3.4426 | 2.0107 | 0.0083 | 0.0139 | 0.0147 | 24387 | 1123 |
| LOF Female, N6 | 0.3759 | 1.5328 | 0.0342 | 0.0066 | 0.0691 | 37546 | 2255 |
| LOF Female, N7 | 0.0000 | 0.9206 | 0.0046 | 0.0072 | 0.0000 | 18970 | 912 |
| LOF Female, N8 |  |  |  |  |  |  |  |
| LOF Female, N9 |  |  |  |  |  |  |  |

| Hippocampus CA3 |  |  |  |  |  |  |  |
| --- | --- | --- | --- | --- | --- | --- | --- |
| genotype, sex,<br>animal n° | TdT-ROR | cFos-ROR | TdT+ cells/<br>DAPI+ cells | cFos+<br>cells/<br>DAPI+ cells | TdT+cFos+<br>cells/<br>cFos+ cells | DAPI+<br>cells | EGFP+<br>cells |
| Control Male, N1 | 0.8205 |  | 0.0022 |  |  | 11070 | 418 |
| Control Female, N2 | 1.0356 | 1.2781 | 0.0031 | 0.0079 | 0.0172 | 7364 | 493 |
| Control Female, N3 | 0.9734 | 0.5296 | 0.0064 | 0.0084 | 0.0000 | 4985 | 450 |
| Control Male, N4 | 1.7679 |  | 0.0026 |  |  | 4978 | 386 |
| Control Female, N5 |  | 0.3655 |  | 0.0054 |  | 12884 | 1225 |
| Control Female, N6 | 0.4267 | 1.4958 | 0.0026 | 0.0024 | 0.0345 | 12172 | 975 |
| Control Male, N7 | 1.2302 | 2.2892 | 0.0023 | 0.0035 | 0.0000 | 8347 | 746 |
| Control Male, N8 |  | 0.4589 |  | 0.0048 |  | 8156 | 902 |
| Control Male, N9 | 0.7457 | 0.5829 | 0.0036 | 0.0061 | 0.0000 | 7155 | 422 |
| Control Male, N10 |  |  |  |  |  |  |  |
| Control Male, N11 |  | 0.9715 |  | 0.0222 |  | 10178 | 939 |
| Control Male, N12 |  |  |  |  |  | 8789 | 1466 |
| Control Male, N13 | 0.6342 | 0.8751 | 0.0023 | 0.0279 | 0.0000 | 10157 | 1729 |
| Control Female, N14 | 0.3644 | 1.2658 | 0.0032 | 0.0202 | 0.0000 | 7685 | 1538 |
| Control Female, N15 | 3.2142 | 1.3607 | 0.0036 | 0.0061 | 0.0000 | 9835 | 1834 |
| Control Female, N16 | 0.2533 | 0.7410 | 0.0045 | 0.0341 | 0.0027 | 10909 | 2371 |
| Control Female, N17 | 1.5339 | 0.8857 | 0.0033 | 0.0200 | 0.0054 | 9178 | 1839 |
| Control Female, N18 |  | 0.8852 |  | 0.0189 |  | 13784 | 2444 |
| Control Female, N19 |  | 1.0150 |  | 0.0032 |  | 18418 | 3359 |
| GOF Male, N1 |  |  |  |  |  |  |  |
| GOF Male, N2 | 0.5372 | 0.8099 | 0.0015 | 0.0077 | 0.0000 | 8686 | 241 |
| GOF Female, N3 | 3.9022 |  | 0.0039 |  |  | 8350 | 175 |
| GOF Female, N4 | 0.6295 |  | 0.0012 |  |  | 12750 | 1083 |
| GOF Male, N5 | 0.4823 | 0.8476 | 0.0013 | 0.0055 | 0.0000 | 8989 | 467 |
| GOF Male, N6 | 0.9214 | 2.0520 | 0.0030 | 0.0097 | 0.0000 | 9114 | 507 |
| GOF Female, N7 | 1.2037 | 2.9971 | 0.0050 | 0.0049 | 0.0244 | 8363 | 685 |
| GOF Female, N8 |  | 4.5339 |  | 0.0084 |  | 7875 | 688 |
| GOF Female, N9 |  | 5.2219 |  | 0.0038 |  | 5300 | 142 |
| GOF Male, N10 | 2.4204 | 4.3027 | 0.0010 | 0.0041 | 0.0345 | 7109 | 173 |
| GOF Male, N11 |  |  |  |  |  |  |  |
| LOF Male, N1 |  |  |  |  |  |  |  |
| LOF Male, N2 |  | 0.8668 |  | 0.0096 |  | 8507 | 798 |
| LOF Male, N3 |  | 0.1449 |  | 0.0151 |  | 8977 | 322 |
| LOF Female, N4 |  | 0.7116 |  | 0.0145 |  | 8219 | 705 |
| LOF Female, N5 | 0.0000 | 0.8589 | 0.0019 | 0.0109 | 0.0000 | 7684 | 870 |
| LOF Female, N6 | 2.2286 | 0.7457 | 0.0019 | 0.0273 | 0.0000 | 9795 | 1221 |
| LOF Female, N7 |  | 1.4548 |  | 0.0059 |  | 7988 | 643 |
| LOF Female, N8 | 0.0000 |  | 0.0019 |  | 0.0000 | 8522 | 577 |
| LOF Female, N9 | 1.9171 | 1.6232 | 0.0023 | 0.0066 | 0.0000 | 9128 | 589 |

| # | 2.2.3 |
| --- | --- |
| subject: | short-term memory, engram |
| reference figures: | Figure 1,2;<br>Supplementary Figure 2,3 |

| Basolateral Amygdala |  |  |  |  |  |  |  |
| --- | --- | --- | --- | --- | --- | --- | --- |
| genotype, sex, animal n° | TdT-ROR | cFos-ROR | TdT+ cells/<br>DAPI+ cells | cFos+ cells/<br>DAPI+ cells | TdT+cFos+<br>cells/<br>cFos+ cells | DAPI+<br>cells | EGFP+<br>cells |
| Control Male, N1 | 1.5881 |  | 0.0068 |  |  | 24086 | 242 |
| Control Female, N2 | 0.9107 | 0.3102 | 0.0099 | 0.0072 | 0.0496 | 16916 | 385 |
| Control Female, N3 | 0.6026 | 0.9319 | 0.0160 | 0.0074 | 0.0000 | 11943 | 401 |
| Control Male, N4 | 2.2632 |  | 0.0085 |  |  | 13345 | 187 |
| Control Female, N5 | 1.1042 | 0.6557 | 0.0033 | 0.0032 | 0.0370 | 25543 | 856 |
| Control Female, N6 | 0.5897 | 1.9442 | 0.0125 | 0.0028 | 0.0377 | 18843 | 410 |
| Control Male, N7 | 1.1444 | 0.7847 | 0.0067 | 0.0033 | 0.0000 | 18675 | 575 |
| Control Male, N8 | 0.0000 | 1.3733 | 0.0009 | 0.0045 | 0.0000 | 13963 | 408 |
| Control Male, N9 | 0.7972 | 1.8523 | 0.0114 | 0.0031 | 0.0000 | 19841 | 380 |
| Control Male, N10 |  |  |  |  |  |  |  |
| Control Male, N11 |  |  |  |  |  |  |  |
| Control Male, N12 | 0.6077 | 0.9825 | 0.0044 | 0.0066 | 0.0078 | 19495 | 1484 |
| Control Male, N13 |  | 0.7788 |  | 0.0132 |  | 19855 | 1268 |
| Control Female, N14 | 1.0090 | 1.1420 | 0.0086 | 0.0064 | 0.0000 | 12032 | 1328 |
| Control Female, N15 | 1.1063 |  | 0.0081 |  |  | 11529 | 957 |
| Control Female, N16 | 0.8183 | 1.0384 | 0.0088 | 0.0126 | 0.0275 | 20241 | 1902 |
| Control Female, N17 | 0.5752 | 0.7179 | 0.0095 | 0.0194 | 0.0201 | 7678 | 1473 |
| Control Female, N18 | 1.8836 | 0.9646 | 0.0112 | 0.0156 | 0.0208 | 18480 | 2314 |
| Control Female, N19 |  | 1.3757 |  | 0.0034 |  | 7359 | 854 |
| GOF Male, N1 | 2.5899 |  | 0.0079 |  |  | 14734 | 130 |
| GOF Male, N2 | 1.6719 | 1.4922 | 0.0046 | 0.0086 | 0.0976 | 14241 | 114 |
| GOF Female, N3 | 0.5734 | 1.1325 | 0.0087 | 0.0089 | 0.0973 | 20878 | 86 |
| GOF Female, N4 | 1.5576 |  | 0.0065 |  |  | 19264 | 349 |
| GOF Male, N5 | 0.3945 | 0.8722 | 0.0056 | 0.0053 | 0.0900 | 18995 | 387 |
| GOF Male, N6 | 0.2806 | 2.9212 | 0.0068 | 0.0055 | 0.0606 | 18006 | 225 |
| GOF Female, N7 | 0.3856 | 4.0580 | 0.0174 | 0.0033 | 0.0000 | 12489 | 380 |
| GOF Female, N8 | 0.7822 |  | 0.0177 |  |  | 12682 | 377 |
| GOF Female, N9 |  |  |  |  |  |  |  |
| GOF Male, N10 |  | 5.2660 |  | 0.0042 | 0.0390 | 18503 | 84 |
| GOF Male, N11 |  | 1.9433 |  | 0.0060 | 0.0175 | 18989 | 292 |
| LOF Male, N1 |  |  |  |  |  |  |  |
| LOF Male, N2 | 0.7365 | 1.2029 | 0.0068 | 0.0046 | 0.0102 | 21254 | 522 |
| LOF Male, N3 | 0.0000 | 0.9133 | 0.0027 | 0.0068 | 0.0145 | 20226 | 226 |
| LOF Female, N4 |  | 0.4692 |  | 0.0064 |  | 19285 | 616 |
| LOF Female, N5 | 0.0000 | 0.6668 | 0.0090 | 0.0091 | 0.0316 | 10391 | 466 |
| LOF Female, N6 | 0.7574 | 2.7850 | 0.0029 | 0.0109 | 0.0000 | 13350 | 602 |
| LOF Female, N7 | 0.6078 | 1.2746 | 0.0183 | 0.0163 | 0.0080 | 7652 | 235 |
| LOF Female, N8 | 0.7572 |  | 0.0081 |  |  | 10533 | 431 |
| LOF Female, N9 | 0.8471 | 1.8666 | 0.0244 | 0.0047 | 0.0610 | 17320 | 844 |

| Hypothalamus |  |  |  |  |  |  |  |
| --- | --- | --- | --- | --- | --- | --- | --- |
| genotype, sex, animal n° | TdT-ROR | cFos-ROR | TdT+ cells/<br>DAPI+ cells | cFos+ cells/<br>DAPI+ cells | TdT+cFos+<br>cells/<br>cFos+ cells | DAPI+<br>cells | EGFP+<br>cells |
| Control Male, N1 | 1.0867 | 0.7623 | 0.0192 | 0.0196 | 0.1949 | 70268 | 1058 |
| Control Female, N2 | 0.7012 | 1.3795 | 0.0290 | 0.0274 | 0.1829 | 42681 | 1103 |
| Control Female, N3 | 1.0673 |  | 0.0204 |  |  | 46299 | 1440 |
| Control Male, N4 | 0.5843 | 0.9777 | 0.0273 | 0.0335 | 0.2961 | 40840 | 1856 |
| Control Female, N5 | 1.5604 | 0.8806 | 0.0231 | 0.0352 | 0.2846 | 46030 | 952 |
| Control Female, N6 |  |  |  |  |  |  |  |
| Control Male, N7 |  |  |  |  |  |  |  |
| Control Male, N8 |  |  |  |  |  |  |  |
| Control Male, N9 |  |  |  |  |  |  |  |
| Control Male, N10 |  |  |  |  |  |  |  |
| Control Male, N11 | 0.2846 |  | 0.0027 |  |  | 38328 | 8624 |
| Control Male, N12 | 0.5013 |  | 0.0180 |  |  | 19638 | 1852 |
| Control Male, N13 |  | 0.5063 |  | 0.0503 |  | 57069 | 4408 |
| Control Female, N14 | 0.9371 | 0.7210 | 0.0258 | 0.0158 | 0.0601 | 46310 | 3688 |
| Control Female, N15 | 1.8822 | 1.1199 | 0.0223 | 0.0257 | 0.0836 | 73005 | 5412 |
| Control Female, N16 | 1.2972 | 1.6529 | 0.0338 | 0.0238 | 0.2811 | 29746 | 3791 |
| Control Female, N17 | 1.0976 |  | 0.0401 |  | 0.2899 | 10129 | 1985 |
| Control Female, N18 |  |  |  |  |  |  |  |
| Control Female, N19 |  |  |  |  |  |  |  |
| GOF Male, N1 | 0.9981 | 0.7344 | 0.0163 | 0.0317 | 0.1453 | 25357 | 859 |
| GOF Male, N2 | 1.2405 | 0.7805 | 0.0088 | 0.0292 | 0.0913 | 43224 | 593 |
| GOF Female, N3 | 0.6898 | 0.7110 | 0.0199 | 0.0230 | 0.2988 | 56148 | 429 |
| GOF Female, N4 | 1.0606 | 0.3512 | 0.0119 | 0.0192 | 0.1363 | 53020 | 1622 |
| GOF Male, N5 | 1.2845 | 1.0605 | 0.0127 | 0.0212 | 0.1638 | 68946 | 2207 |
| GOF Male, N6 | 3.2607 | 1.7064 | 0.0057 | 0.0246 | 0.0850 | 40699 | 354 |
| GOF Female, N7 |  |  |  |  |  |  |  |
| GOF Female, N8 |  |  |  |  |  |  |  |
| GOF Female, N9 |  |  |  |  |  |  |  |
| GOF Male, N10 |  |  |  |  |  |  |  |
| GOF Male, N11 |  |  |  |  |  |  |  |
| LOF Male, N1 | 0.6494 | 0.4160 | 0.0186 | 0.0341 | 0.0339 | 38034 | 3004 |
| LOF Male, N2 | 0.2524 | 0.2039 | 0.0517 | 0.0333 | 0.1323 | 39007 | 2289 |
| LOF Male, N3 | 0.8171 | 0.5271 | 0.0060 | 0.0159 | 0.0333 | 35738 | 418 |
| LOF Female, N4 |  | 0.7807 |  | 0.0199 |  | 55816 | 1422 |
| LOF Female, N5 | 0.6546 | 0.6168 | 0.0307 | 0.0322 | 0.0999 | 51643 | 2345 |
| LOF Female, N6 | 1.0024 | 0.7822 | 0.0624 | 0.0248 | 0.1625 | 46920 | 2479 |
| LOF Female, N7 | 0.4902 | 0.6624 | 0.0244 | 0.0249 | 0.0752 | 56597 | 2325 |
| LOF Female, N8 |  |  |  |  |  |  |  |
| LOF Female, N9 |  |  |  |  |  |  |  |

| Central Amygdala |  |  |  |  |  |  |  |
| --- | --- | --- | --- | --- | --- | --- | --- |
| genotype, sex, animal n° | TdT-ROR | cFos-ROR | TdT+ cells/<br>DAPI+ cells | cFos+ cells/<br>DAPI+ cells | TdT+cFos+<br>cells/<br>cFos+ cells | DAPI+<br>cells | EGFP+<br>cells |
| Control Male, N1 | 0.8878 | 1.4157 | 0.0208 | 0.0092 | 0.2077 | 42561 | 517 |
| Control Female, N2 | 0.6044 | 0.9557 | 0.0320 | 0.0136 | 0.2186 | 15867 | 432 |
| Control Female, N3 | 1.3881 | 0.3998 | 0.0182 | 0.0080 | 0.0760 | 31075 | 1014 |
| Control Male, N4 | 0.7305 | 1.0485 | 0.0345 | 0.0118 | 0.3626 | 22220 | 731 |
| Control Female, N5 | 1.3892 | 1.1804 | 0.0154 | 0.0125 | 0.1458 | 19179 | 264 |
| Control Female, N6 |  |  |  |  |  |  |  |
| Control Male, N7 |  |  |  |  |  |  |  |
| Control Male, N8 |  |  |  |  |  |  |  |
| Control Male, N9 |  |  |  |  |  |  |  |
| Control Male, N10 |  |  |  |  |  |  |  |
| Control Male, N11 | 1.3770 | 0.5948 | 0.0062 | 0.0052 | 0.0125 | 15407 | 548 |
| Control Male, N12 |  | 0.3249 |  | 0.0106 |  | 6702 | 573 |
| Control Male, N13 |  | 0.6189 |  | 0.0157 |  | 25805 | 1646 |
| Control Female, N14 | 0.5508 | 1.7390 | 0.0338 | 0.0121 | 0.0536 | 21601 | 1780 |
| Control Female, N15 | 1.5200 | 1.4137 | 0.0265 | 0.0128 | 0.0204 | 26774 | 2683 |
| Control Female, N16 | 0.4642 | 1.3088 | 0.0324 | 0.0166 | 0.0518 | 18662 | 4008 |
| Control Female, N17 | 1.0879 |  | 0.0220 |  | 0.0735 | 6871 | 1211 |
| Control Female, N18 |  |  |  |  |  |  |  |
| Control Female, N19 |  |  |  |  |  |  |  |
| GOF Male, N1 | 3.7276 | 3.1649 | 0.0051 | 0.0055 | 0.1263 | 17320 | 346 |
| GOF Male, N2 | 3.3988 |  | 0.0091 |  |  | 12627 | 111 |
| GOF Female, N3 | 1.3968 | 1.9286 | 0.0220 | 0.0124 | 0.3364 | 35262 | 257 |
| GOF Female, N4 | 1.8697 | 0.6084 | 0.0133 | 0.0092 | 0.1423 | 28399 | 691 |
| GOF Male, N5 | 0.7296 | 1.0496 | 0.0099 | 0.0087 | 0.0961 | 44305 | 1233 |
| GOF Male, N6 | 1.1562 | 0.8499 | 0.0069 | 0.0153 | 0.0345 | 18940 | 212 |
| GOF Female, N7 |  |  |  |  |  |  |  |
| GOF Female, N8 |  |  |  |  |  |  |  |
| GOF Female, N9 |  |  |  |  |  |  |  |
| GOF Male, N10 |  |  |  |  |  |  |  |
| GOF Male, N11 |  |  |  |  |  |  |  |
| LOF Male, N1 | 1.0183 | 0.6242 | 0.0108 | 0.0053 | 0.0375 | 30273 | 1241 |
| LOF Male, N2 | 0.5562 | 0.5641 | 0.0468 | 0.0080 | 0.0674 | 35178 | 1269 |
| LOF Male, N3 |  |  |  |  |  |  |  |
| LOF Female, N4 |  |  |  |  |  |  |  |
| LOF Female, N5 | 0.8211 | 0.4664 | 0.0384 | 0.0161 | 0.1108 | 19669 | 1078 |
| LOF Female, N6 | 0.5197 | 0.7191 | 0.1277 | 0.0138 | 0.1644 | 36654 | 1755 |
| LOF Female, N7 | 0.3363 | 0.4789 | 0.0205 | 0.0073 | 0.0269 | 25561 | 1255 |
| LOF Female, N8 |  |  |  |  |  |  |  |
| LOF Female, N9 |  |  |  |  |  |  |  |

|  |  |
| --- | --- |
| # | 2.3 |
| subject: | long-term memory, behavior |
| reference figures: | Figure 3; Supplementary Figure 4 |

| Freezing time (s) |  |  |
| --- | --- | --- |
| genotype, sex, animal n° | Training (Ctx A) | Recall (Ctx A) |
| Control Female, N1 | 44.9 | 161.5 |
| Control Female, N2 | 29.4 | 123.4 |
| Control Female, N3 | 49.9 | 118.4 |
| Control Female, N4 | 29.1 | 60.9 |
| Control Male, N5 | 34.9 | 131.2 |
| Control Male, N6 | 52.6 | 49.7 |
| Control Female, N7 | 20.0 | 181.5 |
| Control Female, N8 | 25.4 | 234.8 |
| Control Female, N9 | 64.6 | 267.0 |
| Control Female, N10 | 88.6 | 261.3 |
| Control Male, N11 | 77.2 | 266.7 |
| GOF Male, N1 | 52.6 | 132.3 |
| GOF Male, N2 | 31.8 | 133.8 |
| GOF Female, N3 | 13.0 | 45.6 |
| GOF Female, N4 | 37.3 | 146.0 |
| GOF Female, N5 | 39.0 | 161.9 |
| GOF Male, N6 | 90.5 | 234.4 |
| GOF Male, N7 | 28.8 | 133.9 |
| GOF Male, N8 | 60.2 | 255.0 |
| GOF Male, N9 | 57.1 | 212.2 |
| GOF Female, N10 | 21.3 | 143.4 |
| GOF Female, N11 | 42.2 | 250.3 |
| GOF Female, N12 | 57.5 | 238.4 |
| LOF Female, N1 | 55.5 | 178.9 |
| LOF Female, N2 | 16.3 | 148.8 |
| LOF Male, N3 | 21.3 | 152.3 |
| LOF Male, N4 | 82.1 | 268.1 |
| LOF Male, N5 | 37.8 | 219.9 |
| LOF Male, N6 | 8.1 | 30.5 |
| LOF Male, N7 | 95.0 | 200.3 |
| LOF Male, N8 | 16.5 | 18.9 |
| LOF Male, N9 | 20.4 | 108.4 |

| Freezing latency (s) |  |  |
| --- | --- | --- |
| genotype, sex, animal n° | Training (Ctx A) | Recall (Ctx A) |
| Control Female, N1 | 131.0 | 8.7 |
| Control Female, N2 | 189.9 | 10.8 |
| Control Female, N3 | 144.9 | 21.5 |
| Control Female, N4 | 215.8 | 30.4 |
| Control Male, N5 | 150.6 | 4.4 |
| Control Male, N6 | 148.4 | 96.9 |
| Control Female, N7 | 218.4 | 3.9 |
| Control Female, N8 | 176.8 | 2.4 |
| Control Female, N9 | 200.1 | 0.9 |
| Control Female, N10 | 135.6 | 0.8 |
| Control Male, N11 | 145.4 | 3.6 |
| GOF Male, N1 | 104.4 | 3.1 |
| GOF Male, N2 | 211.1 | 4.9 |
| GOF Female, N3 | 263.8 | 55.7 |
| GOF Female, N4 | 148.1 | 18.8 |
| GOF Female, N5 | 186.3 | 21.3 |
| GOF Male, N6 | 138.8 | 3.6 |
| GOF Male, N7 | 50.4 | 10.9 |
| GOF Male, N8 | 15.4 | 3.3 |
| GOF Male, N9 | 189.6 | 2.3 |
| GOF Female, N10 | 142.4 | 20.4 |
| GOF Female, N11 | 179.0 | 1.6 |
| GOF Female, N12 | 147.2 | 2.3 |
| LOF Female, N1 | 140.9 | 9.7 |
| LOF Female, N2 | 164.2 | 15.7 |
| LOF Male, N3 | 237.0 | 4.1 |
| LOF Male, N4 | 86.5 | 4.2 |
| LOF Male, N5 | 189.8 | 7.4 |
| LOF Male, N6 | 214.6 | 33.8 |
| LOF Male, N7 | 124.5 | 10.3 |
| LOF Male, N8 | 231.6 | 15.6 |
| LOF Male, N9 | 187.5 | 12.5 |

| Freezing episodes |  |  |
| --- | --- | --- |
| genotype, sex, animal n° | Training (Ctx A) | Recall (Ctx A) |
| Control Female, N1 | 17 | 41 |
| Control Female, N2 | 12 | 49 |
| Control Female, N3 | 24 | 48 |
| Control Female, N4 | 13 | 24 |
| Control Male, N5 | 12 | 45 |
| Control Male, N6 | 17 | 14 |
| Control Female, N7 | 9 | 52 |
| Control Female, N8 | 15 | 42 |
| Control Female, N9 | 20 | 27 |
| Control Female, N10 | 25 | 16 |
| Control Male, N11 | 26 | 10 |
| GOF Male, N1 | 23 | 34 |
| GOF Male, N2 | 9 | 40 |
| GOF Female, N3 | 7 | 15 |
| GOF Female, N4 | 21 | 58 |
| GOF Female, N5 | 8 | 49 |
| GOF Male, N6 | 25 | 44 |
| GOF Male, N7 | 17 | 72 |
| GOF Male, N8 | 28 | 37 |
| GOF Male, N9 | 11 | 43 |
| GOF Female, N10 | 13 | 65 |
| GOF Female, N11 | 18 | 28 |
| GOF Female, N12 | 24 | 38 |
| LOF Female, N1 | 19 | 47 |
| LOF Female, N2 | 9 | 41 |
| LOF Male, N3 | 10 | 51 |
| LOF Male, N4 | 22 | 14 |
| LOF Male, N5 | 10 | 31 |
| LOF Male, N6 | 6 | 19 |
| LOF Male, N7 | 27 | 41 |
| LOF Male, N8 | 9 | 11 |
| LOF Male, N9 | 11 | 43 |

| # | 2.4.1 |
| --- | --- |
| subject: | long-term memory, engram |
| reference figures: | Figure 3; Supplementary Figure 4 |

| Pre-limbic Cortex |  |  |  |  |
| --- | --- | --- | --- | --- |
| genotype, sex, animal n° | cFos ROR | cFos+ cells/ DAPI+ cells | DAPI+ cells (n) | EGFP+ cells (n) |
| Control Female, N1 | 1.341 | 0.073 | 18743 | 2063 |
| Control Female, N2 |  |  |  |  |
| Control Female, N3 | 0.908 | 0.031 | 7406 | 1332 |
| Control Female, N4 | 1.010 | 0.088 | 18819 | 2535 |
| Control Male, N5 | 0.741 | 0.044 | 17308 | 2159 |
| Control Male, N6 |  |  |  |  |
| Control Female, N7 | 1.502 | 0.068 | 27620 | 1132 |
| Control Female, N8 | 1.071 | 0.039 | 19985 | 1664 |
| Control Female, N9 | 0.641 | 0.138 | 19806 | 1193 |
| Control Male, N10 | 1.314 | 0.104 | 22778 | 2745 |
| Control Male, N11 | 0.472 | 0.098 | 27197 | 1127 |
| GOF Male, N1 | 1.313 | 0.039 | 31526 | 2485 |
| GOF Male, N2 | 0.907 | 0.036 | 20528 | 1690 |
| GOF Female, N3 | 0.792 | 0.043 | 26228 | 2648 |
| GOF Female, N4 | 0.723 | 0.056 | 22056 | 1674 |
| GOF Female, N5 |  |  |  |  |
| GOF Male, N6 | 1.660 | 0.066 | 12208 | 1244 |
| GOF Male, N7 |  |  |  |  |
| GOF Male, N8 | 0.787 | 0.056 | 10411 | 1597 |
| GOF Male, N9 | 1.158 | 0.066 | 19106 | 1880 |
| GOF Female, N10 | 0.780 | 0.073 | 11539 | 1983 |
| GOF Female, N11 |  |  |  |  |
| GOF Female, N12 |  |  |  |  |
| LOF Female, N1 | 1.382 | 0.022 | 22067 | 3282 |
| LOF Female, N2 | 1.119 | 0.019 | 33395 | 2052 |
| LOF Male, N3 | 1.863 | 0.029 | 22464 | 2516 |
| LOF Male, N4 | 0.770 | 0.029 | 17582 | 2122 |
| LOF Male, N5 | 0.795 | 0.015 | 25796 | 1710 |
| LOF Male, N6 | 1.005 | 0.008 | 25796 | 1792 |
| LOF Male, N7 | 1.068 | 0.025 | 17483 | 2995 |
| LOF Male, N8 | 0.649 | 0.026 | 24091 | 2419 |
| LOF Male, N9 | 0.969 | 0.065 | 21718 | 3353 |

| Motor Cortex |  |  |  |  |
| --- | --- | --- | --- | --- |
| genotype, sex, animal n° | cFos ROR | cFos+ cells/ DAPI+ cells | DAPI+ cells (n) | EGFP+ cells (n) |
| Control Female, N1 |  |  |  |  |
| Control Female, N2 | 0.953 | 0.024 | 41764 | 3071 |
| Control Female, N3 | 0.860 | 0.005 | 18546 | 2068 |
| Control Female, N4 | 1.017 | 0.019 | 30088 | 2395 |
| Control Male, N5 | 1.261 | 0.009 | 9387 | 624 |
| Control Male, N6 | 0.909 | 0.010 | 56758 | 2819 |
| Control Female, N7 | 1.334 | 0.017 | 58291 | 4893 |
| Control Female, N8 | 0.701 | 0.025 | 50832 | 3928 |
| Control Female, N9 | 0.731 | 0.020 | 50255 | 7665 |
| Control Male, N10 | 1.489 | 0.013 | 20472 | 1141 |
| Control Male, N11 | 0.746 | 0.019 | 41121 | 1981 |
| GOF Male, N1 | 1.622 | 0.005 | 62222 | 4343 |
| GOF Male, N2 |  |  |  |  |
| GOF Female, N3 | 0.851 | 0.005 | 40515 | 5831 |
| GOF Female, N4 | 0.797 | 0.012 | 52009 | 3974 |
| GOF Female, N5 | 0.796 | 0.010 | 51417 | 7289 |
| GOF Male, N6 | 0.940 | 0.013 | 16912 | 1103 |
| GOF Male, N7 | 1.547 | 0.008 | 36863 | 2074 |
| GOF Male, N8 | 1.261 | 0.025 | 31753 | 1710 |
| GOF Male, N9 |  |  |  |  |
| GOF Female, N10 | 1.487 | 0.014 | 43852 | 2997 |
| GOF Female, N11 | 2.227 | 0.010 | 26730 | 1815 |
| GOF Female, N12 | 0.877 | 0.025 | 21217 | 1524 |
| LOF Female, N1 |  |  |  |  |
| LOF Female, N2 | 0.453 | 0.007 | 25363 | 4855 |
| LOF Male, N3 | 0.570 | 0.012 | 21869 | 1956 |
| LOF Male, N4 | 0.534 | 0.005 | 53925 | 7572 |
| LOF Male, N5 | 0.266 | 0.011 | 35930 | 5357 |
| LOF Male, N6 | 0.237 | 0.011 | 45132 | 4555 |
| LOF Male, N7 | 0.386 | 0.008 | 48886 | 8781 |
| LOF Male, N8 | 0.850 | 0.022 | 51555 | 6320 |
| LOF Male, N9 |  |  |  |  |

| Infra-limbic Cortex |  |  |  |  |
| --- | --- | --- | --- | --- |
| genotype, sex, animal n° | cFos ROR | cFos+ cells/ DAPI+ cells | DAPI+ cells (n) | EGFP+ cells (n) |
| Control Female, N1 | 1.074 | 0.055 | 12399 | 1506 |
| Control Female, N2 | 0.617 | 0.064 | 15256 | 950 |
| Control Female, N3 | 1.010 | 0.033 | 6750 | 1193 |
| Control Female, N4 | 1.570 | 0.047 | 16601 | 980 |
| Control Male, N5 | 0.729 | 0.037 | 12933 | 844 |
| Control Male, N6 |  |  |  |  |
| Control Female, N7 | 1.470 | 0.021 | 23524 | 627 |
| Control Female, N8 | 1.004 | 0.023 | 19674 | 1097 |
| Control Female, N9 | 0.898 | 0.052 | 18251 | 1043 |
| Control Male, N10 | 1.107 | 0.032 | 20166 | 1827 |
| Control Male, N11 | 0.521 | 0.030 | 20213 | 813 |
| GOF Male, N1 | 1.448 | 0.024 | 28659 | 2155 |
| GOF Male, N2 | 0.985 | 0.032 | 17732 | 1159 |
| GOF Female, N3 | 0.985 | 0.041 | 26005 | 2141 |
| GOF Female, N4 | 1.273 | 0.022 | 21344 | 1525 |
| GOF Female, N5 |  |  |  |  |
| GOF Male, N6 | 0.901 | 0.025 | 8056 | 935 |
| GOF Male, N7 | 1.264 | 0.034 | 7617 | 890 |
| GOF Male, N8 | 0.576 | 0.031 | 6488 | 896 |
| GOF Male, N9 | 1.043 | 0.018 | 14202 | 1326 |
| GOF Female, N10 | 0.958 | 0.060 | 6823 | 778 |
| GOF Female, N11 |  |  |  |  |
| GOF Female, N12 |  |  |  |  |
| LOF Female, N1 |  |  |  |  |
| LOF Female, N2 |  |  |  |  |
| LOF Male, N3 | 0.929 | 0.025 | 21781 | 1751 |
| LOF Male, N4 | 0.813 | 0.025 | 14452 | 1660 |
| LOF Male, N5 | 0.762 | 0.022 | 23154 | 1571 |
| LOF Male, N6 | 0.970 | 0.013 | 23154 | 1669 |
| LOF Male, N7 | 1.085 | 0.028 | 14403 | 1822 |
| LOF Male, N8 | 0.786 | 0.028 | 20547 | 2048 |
| LOF Male, N9 | 0.712 | 0.065 | 16067 | 1773 |

| Somatosensory Cortex |  |  |  |  |
| --- | --- | --- | --- | --- |
| genotype, sex, animal n° | cFos ROR | cFos+ cells/ DAPI+ cells | DAPI+ cells (n) | EGFP+ cells (n) |
| Control Female, N1 |  |  |  |  |
| Control Female, N2 | 1.085 | 0.028 | 75176 | 2730 |
| Control Female, N3 | 1.156 | 0.005 | 34685 | 2972 |
| Control Female, N4 | 1.191 | 0.021 | 51273 | 2980 |
| Control Male, N5 | 0.682 | 0.005 | 22406 | 1211 |
| Control Male, N6 | 0.886 | 0.013 | 89888 | 2863 |
| Control Female, N7 | 1.028 | 0.015 | 100148 | 7040 |
| Control Female, N8 |  |  |  |  |
| Control Female, N9 | 1.020 | 0.016 | 69898 | 9271 |
| Control Male, N10 | 1.162 | 0.006 | 24352 | 1430 |
| Control Male, N11 | 0.790 | 0.023 | 88918 | 3422 |
| GOF Male, N1 |  |  |  |  |
| GOF Male, N2 |  |  |  |  |
| GOF Female, N3 | 1.020 | 0.003 | 82800 | 5732 |
| GOF Female, N4 |  |  |  |  |
| GOF Female, N5 | 0.917 | 0.006 | 79755 | 6883 |
| GOF Male, N6 | 0.635 | 0.010 | 53577 | 2322 |
| GOF Male, N7 | 1.502 | 0.002 | 56727 | 2669 |
| GOF Male, N8 | 1.385 | 0.015 | 47019 | 2262 |
| GOF Male, N9 | 1.210 | 0.011 | 63918 | 3224 |
| GOF Female, N10 | 2.003 | 0.005 | 79323 | 3088 |
| GOF Female, N11 | 2.658 | 0.004 | 34583 | 1975 |
| GOF Female, N12 | 1.016 | 0.017 | 37398 | 1748 |
| LOF Female, N1 |  |  |  |  |
| LOF Female, N2 |  |  |  |  |
| LOF Male, N3 | 1.193 | 0.016 | 52827 | 3261 |
| LOF Male, N4 | 0.406 | 0.005 | 107610 | 10038 |
| LOF Male, N5 | 0.840 | 0.006 | 72242 | 7460 |
| LOF Male, N6 |  |  |  |  |
| LOF Male, N7 | 0.925 | 0.004 | 87557 | 10526 |
| LOF Male, N8 | 0.768 | 0.015 | 120486 | 13254 |
| LOF Male, N9 |  |  |  |  |

| # | 2.4.2 |
| --- | --- |
| subject: | long-term memory, engram |
| reference figures: | Figure 3; Supplementary Figure 4 |

| Auditory Cortex |  |  |  |  |
| --- | --- | --- | --- | --- |
| genotype, sex, animal n° | cFos ROR | cFos+ cells/ DAPI+ cells | DAPI+ cells (n) | EGFP+ cells (n) |
| Control Female, N1 |  |  |  |  |
| Control Female, N2 | 1.050 | 0.038 | 41037 | 1683 |
| Control Female, N3 | 0.699 | 0.021 | 18177 | 1610 |
| Control Female, N4 | 1.221 | 0.016 | 30384 | 1376 |
| Control Male, N5 | 1.161 | 0.007 | 7261 | 526 |
| Control Male, N6 | 0.869 | 0.015 | 31739 | 1274 |
| Control Female, N7 | 0.804 | 0.016 | 27242 | 2170 |
| Control Female, N8 | 0.827 | 0.029 | 24370 | 1492 |
| Control Female, N9 | 1.181 | 0.016 | 19168 | 2242 |
| Control Male, N10 | 1.187 | 0.015 | 11285 | 491 |
| Control Male, N11 |  |  |  |  |
| GOF Male, N1 |  |  |  |  |
| GOF Male, N2 |  |  |  |  |
| GOF Female, N3 | 1.881 | 0.007 | 26629 | 1835 |
| GOF Female, N4 | 1.210 | 0.008 | 23241 | 1579 |
| GOF Female, N5 | 0.775 | 0.008 | 28149 | 2414 |
| GOF Male, N6 |  |  |  |  |
| GOF Male, N7 | 0.574 | 0.008 | 22348 | 1547 |
| GOF Male, N8 | 0.885 | 0.023 | 19619 | 1170 |
| GOF Male, N9 | 1.037 | 0.014 | 40729 | 1695 |
| GOF Female, N10 | 1.943 | 0.008 | 44231 | 1636 |
| GOF Female, N11 | 1.593 | 0.007 | 15516 | 834 |
| GOF Female, N12 | 0.745 | 0.024 | 17801 | 953 |
| LOF Female, N1 |  |  |  |  |
| LOF Female, N2 | 0.655 | 0.006 | 23087 | 3487 |
| LOF Male, N3 | 0.614 | 0.021 | 16895 | 958 |
| LOF Male, N4 | 1.153 | 0.006 | 26399 | 2371 |
| LOF Male, N5 | 0.495 | 0.008 | 23963 | 2624 |
| LOF Male, N6 | 0.427 | 0.004 | 24175 | 1712 |
| LOF Male, N7 | 0.754 | 0.005 | 21749 | 3067 |
| LOF Male, N8 | 1.232 | 0.008 | 46640 | 3829 |
| LOF Male, N9 |  |  |  |  |

| Dentate Gyrus |  |  |  |  |
| --- | --- | --- | --- | --- |
| genotype, sex, animal n° | cFos ROR | cFos+ cells/ DAPI+ cells | DAPI+ cells (n) | EGFP+ cells (n) |
| Control Female, N1 | 0.967 | 0.030 | 17911 | 2237 |
| Control Female, N2 | 0.582 | 0.038 | 15690 | 895 |
| Control Female, N3 | 1.013 | 0.023 | 9623 | 1125 |
| Control Female, N4 | 1.194 | 0.033 | 14061 | 1419 |
| Control Male, N5 | 0.979 | 0.027 | 8278 | 923 |
| Control Male, N6 | 1.264 | 0.032 | 10250 | 815 |
| Control Female, N7 | 1.162 | 0.016 | 9596 | 1660 |
| Control Female, N8 | 1.197 | 0.012 | 11505 | 1591 |
| Control Female, N9 | 1.322 | 0.017 | 10478 | 2763 |
| Control Male, N10 | 0.809 | 0.016 | 11679 | 1234 |
| Control Male, N11 | 0.510 | 0.015 | 12547 | 1301 |
| GOF Male, N1 | 0.884 | 0.021 | 12800 | 2099 |
| GOF Male, N2 | 0.840 | 0.015 | 12033 | 3061 |
| GOF Female, N3 | 1.071 | 0.024 | 13556 | 3146 |
| GOF Female, N4 | 0.835 | 0.010 | 9852 | 1736 |
| GOF Female, N5 | 0.933 | 0.024 | 13664 | 3128 |
| GOF Male, N6 | 1.083 | 0.016 | 12353 | 718 |
| GOF Male, N7 | 0.545 | 0.015 | 15616 | 914 |
| GOF Male, N8 |  |  |  | 595 |
| GOF Male, N9 | 0.571 | 0.025 | 14395 | 844 |
| GOF Female, N10 | 1.213 | 0.017 | 17134 | 933 |
| GOF Female, N11 | 0.781 | 0.018 | 13974 | 920 |
| GOF Female, N12 | 0.737 | 0.024 | 9334 | 577 |
| LOF Female, N1 | 1.022 | 0.019 | 10908 | 3425 |
| LOF Female, N2 | 0.661 | 0.029 | 8608 | 1454 |
| LOF Male, N3 | 0.736 | 0.013 | 12571 | 3012 |
| LOF Male, N4 | 0.847 | 0.012 | 15027 | 3110 |
| LOF Male, N5 | 0.839 | 0.007 | 15498 | 2199 |
| LOF Male, N6 | 0.839 | 0.011 | 12126 | 3399 |
| LOF Male, N7 | 0.785 | 0.011 | 11738 | 2473 |
| LOF Male, N8 | 0.968 | 0.025 | 14824 | 3834 |
| LOF Male, N9 |  |  |  |  |

| Piriform Cortex |  |  |  |  |
| --- | --- | --- | --- | --- |
| genotype, sex, animal n° | cFos ROR | cFos+ cells/ DAPI+ cells | DAPI+ cells (n) | EGFP+ cells (n) |
| Control Female, N1 | 1.234 | 0.029 | 21467 | 1720 |
| Control Female, N2 | 1.001 | 0.031 | 36706 | 2089 |
| Control Female, N3 | 0.945 | 0.015 | 12419 | 634 |
| Control Female, N4 | 0.967 | 0.021 | 21076 | 1164 |
| Control Male, N5 | 0.854 | 0.012 | 4347 | 755 |
| Control Male, N6 |  |  |  |  |
| Control Female, N7 | 0.856 | 0.024 | 22783 | 1169 |
| Control Female, N8 | 0.836 | 0.032 | 29764 | 1751 |
| Control Female, N9 | 1.014 | 0.012 | 22626 | 1980 |
| Control Male, N10 | 1.294 | 0.022 | 10379 | 680 |
| Control Male, N11 |  |  |  |  |
| GOF Male, N1 | 1.983 | 0.013 | 28164 | 1272 |
| GOF Male, N2 |  |  |  |  |
| GOF Female, N3 | 0.786 | 0.018 | 17991 | 1124 |
| GOF Female, N4 | 1.070 | 0.012 | 25783 | 1163 |
| GOF Female, N5 | 0.680 | 0.020 | 33507 | 2385 |
| GOF Male, N6 |  |  |  |  |
| GOF Male, N7 | 0.586 | 0.005 | 11252 | 815 |
| GOF Male, N8 | 1.266 | 0.027 | 21509 | 1264 |
| GOF Male, N9 | 1.949 | 0.018 | 40162 | 2216 |
| GOF Female, N10 | 1.278 | 0.020 | 46305 | 1999 |
| GOF Female, N11 | 2.117 | 0.010 | 21546 | 1260 |
| GOF Female, N12 | 0.704 | 0.018 | 23442 | 1267 |
| LOF Female, N1 |  |  |  |  |
| LOF Female, N2 | 0.889 | 0.028 | 33986 | 3950 |
| LOF Male, N3 | 0.825 | 0.022 | 25689 | 1266 |
| LOF Male, N4 | 0.527 | 0.019 | 33661 | 2168 |
| LOF Male, N5 | 0.749 | 0.011 | 32292 | 2875 |
| LOF Male, N6 | 1.105 | 0.013 | 34398 | 1497 |
| LOF Male, N7 | 0.814 | 0.010 | 31877 | 4076 |
| LOF Male, N8 | 0.910 | 0.011 | 33260 | 2009 |
| LOF Male, N9 |  |  |  |  |

| Hippocampus CA3 |  |  |  |  |
| --- | --- | --- | --- | --- |
| genotype, sex, animal n° | cFos ROR | cFos+ cells/ DAPI+ cells | DAPI+ cells (n) | EGFP+ cells (n) |
| Control Female, N1 | 0.872 | 0.039 | 10838 | 1309 |
| Control Female, N2 |  |  |  |  |
| Control Female, N3 | 0.902 | 0.015 | 6004 | 604 |
| Control Female, N4 | 1.097 | 0.057 | 8111 | 662 |
| Control Male, N5 | 1.010 | 0.034 | 4815 | 331 |
| Control Male, N6 | 1.119 | 0.044 | 8183 | 465 |
| Control Female, N7 | 0.927 | 0.020 | 7994 | 875 |
| Control Female, N8 | 1.260 | 0.010 | 7140 | 813 |
| Control Female, N9 | 0.819 | 0.012 | 6762 | 1653 |
| Control Male, N10 | 1.233 | 0.014 | 6624 | 516 |
| Control Male, N11 | 0.761 | 0.014 | 10420 | 633 |
| GOF Male, N1 | 0.880 | 0.025 | 7981 | 1015 |
| GOF Male, N2 | 1.686 | 0.014 | 6069 | 1051 |
| GOF Female, N3 | 0.865 | 0.014 | 9063 | 1389 |
| GOF Female, N4 | 0.654 | 0.008 | 7441 | 975 |
| GOF Female, N5 | 1.014 | 0.022 | 7023 | 1287 |
| GOF Male, N6 | 1.070 | 0.013 | 10322 | 575 |
| GOF Male, N7 | 0.653 | 0.005 | 12458 | 842 |
| GOF Male, N8 | 0.824 | 0.014 | 7431 | 255 |
| GOF Male, N9 | 0.594 | 0.024 | 9796 | 461 |
| GOF Female, N10 | 0.762 | 0.012 | 9374 | 611 |
| GOF Female, N11 | 0.653 | 0.015 | 8959 | 715 |
| GOF Female, N12 | 1.016 | 0.019 | 6457 | 346 |
| LOF Female, N1 | 0.511 | 0.028 | 4802 | 1352 |
| LOF Female, N2 | 0.902 | 0.036 | 6252 | 668 |
| LOF Male, N3 | 0.764 | 0.014 | 8836 | 1502 |
| LOF Male, N4 |  |  |  |  |
| LOF Male, N5 | 0.405 | 0.007 | 9169 | 915 |
| LOF Male, N6 | 0.570 | 0.016 | 7722 | 1750 |
| LOF Male, N7 | 1.200 | 0.014 | 7989 | 1040 |
| LOF Male, N8 | 0.926 | 0.028 | 8474 | 1633 |
| LOF Male, N9 |  |  |  |  |

| # | 2.4.3 |
| --- | --- |
| subject: | long-term memory, engram |
| reference figures: | Figure 3; Supplementary Figure 4 |

| Basolateral Amygdala |  |  |  |  |
| --- | --- | --- | --- | --- |
| genotype, sex, animal n° | cFos ROR | cFos+ cells/ DAPI+ cells | DAPI+ cells (n) | EGFP+ cells (n) |
| Control Female, N1 | 0.600 | 0.018 | 14238 | 1030 |
| Control Female, N2 | 0.458 | 0.020 | 10714 | 627 |
| Control Female, N3 | 0.804 | 0.010 | 6026 | 295 |
| Control Female, N4 | 1.218 | 0.011 | 9585 | 532 |
| Control Male, N5 | 1.272 | 0.010 | 1420 | 263 |
| Control Male, N6 | 1.648 | 0.008 | 12311 | 341 |
| Control Female, N7 | 1.074 | 0.019 | 10332 | 682 |
| Control Female, N8 | 1.250 | 0.012 | 18067 | 888 |
| Control Female, N9 | 0.842 | 0.009 | 12505 | 1103 |
| Control Male, N10 | 0.834 | 0.032 | 13348 | 413 |
| Control Male, N11 |  |  |  |  |
| GOF Male, N1 | 0.935 | 0.010 | 17591 | 842 |
| GOF Male, N2 | 0.431 | 0.019 | 13948 | 1511 |
| GOF Female, N3 | 0.517 | 0.013 | 10320 | 724 |
| GOF Female, N4 | 0.571 | 0.011 | 12860 | 699 |
| GOF Female, N5 | 0.559 | 0.016 | 11424 | 1294 |
| GOF Male, N6 | 0.627 | 0.022 | 8750 | 532 |
| GOF Male, N7 | 1.665 | 0.007 | 5377 | 344 |
| GOF Male, N8 | 1.648 | 0.017 | 7648 | 336 |
| GOF Male, N9 | 1.253 | 0.017 | 16454 | 556 |
| GOF Female, N10 | 0.844 | 0.014 | 19052 | 786 |
| GOF Female, N11 | 0.615 | 0.014 | 11997 | 619 |
| GOF Female, N12 | 0.603 | 0.012 | 10462 | 524 |
| LOF Female, N1 | 0.751 | 0.009 | 9276 | 1203 |
| LOF Female, N2 | 0.851 | 0.010 | 9987 | 694 |
| LOF Male, N3 | 0.632 | 0.016 | 16832 | 1469 |
| LOF Male, N4 | 0.443 | 0.018 | 11218 | 1287 |
| LOF Male, N5 | 0.542 | 0.015 | 14250 | 851 |
| LOF Male, N6 | 0.980 | 0.008 | 13323 | 1834 |
| LOF Male, N7 | 1.026 | 0.009 | 11159 | 693 |
| LOF Male, N8 |  |  |  |  |
| LOF Male, N9 |  |  |  |  |

| Hypothalamus |  |  |  |  |
| --- | --- | --- | --- | --- |
| genotype, sex, animal n° | cFos ROR | cFos+ cells/ DAPI+ cells | DAPI+ cells (n) | EGFP+ cells (n) |
| Control Female, N1 | 0.754 | 0.030 | 58318 | 4905 |
| Control Female, N2 |  |  |  |  |
| Control Female, N3 | 1.039 | 0.011 | 14823 | 901 |
| Control Female, N4 | 1.367 | 0.029 | 22223 | 1322 |
| Control Male, N5 | 0.839 | 0.031 | 5912 | 723 |
| Control Male, N6 |  |  |  |  |
| Control Female, N7 | 1.468 | 0.035 | 30940 | 1368 |
| Control Female, N8 | 0.704 | 0.029 | 39646 | 1648 |
| Control Female, N9 | 0.491 | 0.039 | 33171 | 2548 |
| Control Male, N10 | 1.337 | 0.087 | 41131 | 2137 |
| Control Male, N11 |  |  |  |  |
| GOF Male, N1 | 0.733 | 0.028 | 53348 | 4218 |
| GOF Male, N2 |  |  |  |  |
| GOF Female, N3 | 0.781 | 0.027 | 41772 | 4384 |
| GOF Female, N4 | 0.428 | 0.029 | 33075 | 2384 |
| GOF Female, N5 | 0.515 | 0.031 | 40441 | 4242 |
| GOF Male, N6 | 0.615 | 0.042 | 20350 | 1464 |
| GOF Male, N7 | 0.587 | 0.040 | 17941 | 1558 |
| GOF Male, N8 | 0.454 | 0.017 | 25196 | 943 |
| GOF Male, N9 | 0.492 | 0.036 | 30193 | 1341 |
| GOF Female, N10 | 0.557 | 0.037 | 36191 | 2737 |
| GOF Female, N11 | 0.791 | 0.029 | 26343 | 1280 |
| GOF Female, N12 |  |  |  |  |
| LOF Female, N1 | 0.508 | 0.037 | 35440 | 3856 |
| LOF Female, N2 | 0.702 | 0.016 | 36228 | 2350 |
| LOF Male, N3 |  |  |  |  |
| LOF Male, N4 | 0.593 | 0.022 | 41549 | 4360 |
| LOF Male, N5 | 0.599 | 0.024 | 46982 | 3018 |
| LOF Male, N6 | 0.546 | 0.014 | 51483 | 5470 |
| LOF Male, N7 |  |  |  |  |
| LOF Male, N8 |  |  |  |  |
| LOF Male, N9 |  |  |  |  |

| Central Amygdala |  |  |  |  |
| --- | --- | --- | --- | --- |
| genotype, sex, animal n° | cFos ROR | cFos+ cells/ DAPI+ cells | DAPI+ cells (n) | EGFP+ cells (n) |
| Control Female, N1 | 1.288 | 0.012 | 14770 | 803 |
| Control Female, N2 |  |  |  |  |
| Control Female, N3 | 0.902 | 0.014 | 7417 | 470 |
| Control Female, N4 | 0.849 | 0.011 | 14172 | 941 |
| Control Male, N5 | 0.961 | 0.008 | 2542 | 380 |
| Control Male, N6 |  |  |  | 695 |
| Control Female, N7 | 1.297 | 0.027 | 13122 | 848 |
| Control Female, N8 | 1.042 | 0.016 | 20915 | 867 |
| Control Female, N9 | 0.800 | 0.025 | 18947 | 1731 |
| Control Male, N10 | 0.861 | 0.059 | 19712 | 996 |
| Control Male, N11 |  |  |  |  |
| GOF Male, N1 | 0.852 | 0.012 | 26010 | 2371 |
| GOF Male, N2 | 0.765 | 0.014 | 14887 | 1738 |
| GOF Female, N3 | 0.846 | 0.010 | 14240 | 1370 |
| GOF Female, N4 | 0.533 | 0.012 | 23217 | 1426 |
| GOF Female, N5 | 0.429 | 0.010 | 19181 | 2427 |
| GOF Male, N6 | 0.485 | 0.024 | 10759 | 513 |
| GOF Male, N7 | 0.650 | 0.022 | 7054 | 430 |
| GOF Male, N8 |  |  |  |  |
| GOF Male, N9 | 1.739 | 0.014 | 19769 | 905 |
| GOF Female, N10 | 0.745 | 0.020 | 23174 | 953 |
| GOF Female, N11 |  |  |  |  |
| GOF Female, N12 | 1.522 | 0.020 | 13150 | 615 |
| LOF Female, N1 | 0.675 | 0.012 | 14217 | 2061 |
| LOF Female, N2 | 0.651 | 0.011 | 15385 | 1190 |
| LOF Male, N3 | 0.479 | 0.013 | 21658 | 2344 |
| LOF Male, N4 | 0.512 | 0.016 | 17384 | 2167 |
| LOF Male, N5 | 0.455 | 0.012 | 18693 | 1087 |
| LOF Male, N6 | 0.463 | 0.010 | 21262 | 2417 |
| LOF Male, N7 | 0.631 | 0.009 | 13465 | 1258 |
| LOF Male, N8 |  |  |  |  |
| LOF Male, N9 |  |  |  |  |

|  |  |
| --- | --- |
| # | 2.5 |
| subject: | chemogenetics,<br>behavior |
| reference figures: | Figure 4 |

| Freezing Time (s) |  |  |
| --- | --- | --- |
| genotype, sex,<br>animal n° | Training | Retrieval |
| Control Male, N1 | 23.4 | 144.9 |
| Control Male, N2 | 74.6 | 218.1 |
| Control Male, N3 | 88 | 254.4 |
| Control Female, N4 | 53.4 | 184.1 |
| Control Female, N5 | 35.2 | 178.9 |
| Control Female, N6 | 42.7 | 185 |
| Control Male, N7 | 17.8 | 155.9 |
| Control Female, N8 | 38.9 | 181.8 |
| Ctrl Gq Female, N1 | 3.3 | 115 |
| Ctrl Gq Female, N2 | 61.7 | 262.6 |
| Ctrl Gq Male, N3 | 18.8 | 148.1 |
| Ctrl Gq Male, N4 | 26.2 | 149.7 |
| Ctrl Gq Male, N5 | 31.9 | 50 |
| Ctrl Gi Male, N1 | 25 | 110.7 |
| Ctrl Gi Male, N2 | 13.2 | 59.1 |
| Ctrl Gi Male, N3 | 10.4 | 143.4 |
| Ctrl Gi Female, N4 | 58.3 | 63.8 |
| Ctrl Gi Female, N5 | 8 | 50.2 |
| Ctrl Gi Female, N6 | 56.9 | 114.2 |
| LOF Gq Male, N1 | 72.6 | 234.8 |
| LOF Gq Male, N2 | 59.4 | 215.2 |
| LOF Gq Male, N3 | 61 | 159.9 |
| LOF Gq Female, N4 | 108.5 | 224.1 |
| LOF Gq Female, N5 | 50.1 | 135.3 |
| LOF Gq Female, N6 | 39.6 | 151.6 |
| GOF Gi Male, N1 | 39 | 149.9 |
| GOF Gi Male, N2 | 104.4 | 168.3 |
| GOF Gi Male, N3 | 42.1 | 89.6 |
| GOF Gi Female, N4 | 81.3 | 207.6 |
| GOF Gi Female, N5 | 44.8 | 111.7 |
| GOF Gi Female, N6 | 50.5 | 183.4 |

| Freezing Latency (s) |  |  |
| --- | --- | --- |
| genotype, sex,<br>animal n° | Training | Retrieval |
| Control Male, N1 | 108 | 8.9 |
| Control Male, N2 | 22 | 2.4 |
| Control Male, N3 | 80.1 | 2.5 |
| Control Female, N4 | 156.4 | 3.1 |
| Control Female, N5 | 152.2 | 7.2 |
| Control Female, N6 | 132.2 | 5.2 |
| Control Male, N7 | 235.2 | 24.8 |
| Control Female, N8 | 166.5 | 27.9 |
| Ctrl Gq Female, N1 | 284.9 | 3 |
| Ctrl Gq Female, N2 | 144.1 | 1.8 |
| Ctrl Gq Male, N3 | 193.2 | 1.4 |
| Ctrl Gq Male, N4 | 168.3 | 3.3 |
| Ctrl Gq Male, N5 | 196.6 | 17 |
| Ctrl Gi Male, N1 | 54.2 | 4.8 |
| Ctrl Gi Male, N2 | 266 | 3.5 |
| Ctrl Gi Male, N3 | 166.5 | 2.4 |
| Ctrl Gi Female, N4 | 141.6 | 36.3 |
| Ctrl Gi Female, N5 | 263 | 19.1 |
| Ctrl Gi Female, N6 | 156.7 | 1.9 |
| LOF Gq Male, N1 | 86.6 | 2.5 |
| LOF Gq Male, N2 | 144.7 | 2.5 |
| LOF Gq Male, N3 | 168 | 4.6 |
| LOF Gq Female, N4 | 125 | 2.6 |
| LOF Gq Female, N5 | 84.4 | 4.1 |
| LOF Gq Female, N6 | 189.6 | 2.1 |
| GOF Gi Male, N1 | 153.2 | 2 |
| GOF Gi Male, N2 | 32.5 | 1.2 |
| GOF Gi Male, N3 | 148.9 | 13.4 |
| GOF Gi Female, N4 | 32.6 | 1.5 |
| GOF Gi Female, N5 | 41.1 | 8.2 |
| GOF Gi Female, N6 | 153.6 | 14.9 |

| Freezing episodes |  |  |
| --- | --- | --- |
| genotype, sex,<br>animal n° | Training | Retrieval |
| Control Male, N1 | 15 | 54 |
| Control Male, N2 | 32 | 49 |
| Control Male, N3 | 31 | 39 |
| Control Female, N4 | 19 | 50 |
| Control Female, N5 | 18 | 71 |
| Control Female, N6 | 21 | 76 |
| Control Male, N7 | 6 | 22 |
| Control Female, N8 | 16 | 36 |
| Ctrl Gq Female, N1 | 2 | 60 |
| Ctrl Gq Female, N2 | 28 | 40 |
| Ctrl Gq Male, N3 | 8 | 56 |
| Ctrl Gq Male, N4 | 12 | 34 |
| Ctrl Gq Male, N5 | 12 | 27 |
| Ctrl Gi Male, N1 | 11 | 45 |
| Ctrl Gi Male, N2 | 6 | 15 |
| Ctrl Gi Male, N3 | 7 | 61 |
| Ctrl Gi Female, N4 | 26 | 28 |
| Ctrl Gi Female, N5 | 4 | 26 |
| Ctrl Gi Female, N6 | 21 | 55 |
| LOF Gq Male, N1 | 27 | 51 |
| LOF Gq Male, N2 | 26 | 42 |
| LOF Gq Male, N3 | 17 | 66 |
| LOF Gq Female, N4 | 24 | 52 |
| LOF Gq Female, N5 | 26 | 49 |
| LOF Gq Female, N6 | 18 | 63 |
| GOF Gi Male, N1 | 20 | 74 |
| GOF Gi Male, N2 | 32 | 41 |
| GOF Gi Male, N3 | 19 | 43 |
| GOF Gi Female, N4 | 30 | 55 |
| GOF Gi Female, N5 | 26 | 61 |
| GOF Gi Female, N6 | 12 | 42 |

|  |  |
| --- | --- |
| # | 2.6 |
| subject: | KA assays |
| reference figures: | Supplementary Figure 5A-G |

| Suppl Fig. 5 B,E | Gapdh-, Ctrl-double norm cFos mRNA |  |  |  |
| --- | --- | --- | --- | --- |
| group, structure, n° | 2h | 6h | 8h | 24h |
| Ctrl Cortex N1 | 0.944 | 0.824 | 1.072 | 1.430 |
| Ctrl Cortex N2 | 1.110 | 1.130 | 0.851 | 1.022 |
| Ctrl Cortex N3 | 1.247 | 1.040 | 0.863 | 1.018 |
| Ctrl Cortex N4 | 1.141 | 1.005 | 1.250 | 0.739 |
| Ctrl Cortex N5 | 0.559 |  | 0.965 | 0.791 |
| KA Cortex N1 | 1.755 | 0.761 | 0.792 | 1.634 |
| KA Cortex N2 | 1.310 | 4.584 | 1.400 | 1.812 |
| KA Cortex N3 | 1.052 | 0.378 | 2.130 | 1.584 |
| KA Cortex N4 | 1.041 | 1.384 | 2.665 | 0.736 |
| KA Cortex N5 | 1.172 |  | 1.400 | 1.779 |
| Ctrl Hippo N1 | 0.783 | 0.331 | 0.935 | 1.305 |
| Ctrl Hippo N2 | 1.197 | 1.940 | 0.777 | 1.036 |
| Ctrl Hippo N3 | 0.896 | 0.970 | 1.398 | 1.301 |
| Ctrl Hippo N4 | 1.192 | 0.759 | 0.950 | 0.789 |
| Ctrl Hippo N5 | 0.932 |  | 0.940 | 0.569 |
| KA Hippo N1 | 35.688 | 8.162 | 0.325 | 0.721 |
| KA Hippo N2 | 1.526 | 25.807 | 0.981 | 0.661 |
| KA Hippo N3 | 2.488 | 1.096 | 1.408 | 1.295 |
| KA Hippo N4 | 1.130 | 1.213 | 1.432 | 0.810 |
| KA Hippo N5 | 15.335 |  | 0.912 | 0.588 |

| Suppl Fig. C,F | Gapdh-, Ctrl-double norm Arc mRNA |  |  |  |
| --- | --- | --- | --- | --- |
| group, structure, n° | 2h | 6h | 8h | 24h |
| Ctrl Cortex N1 | 1.246 | 0.696 | 0.873 | 1.351 |
| Ctrl Cortex N2 | 0.765 | 0.668 | 0.880 | 0.977 |
| Ctrl Cortex N3 | 1.167 | 1.363 | 0.857 | 0.892 |
| Ctrl Cortex N4 | 1.150 | 1.273 | 0.562 | 0.845 |
| Ctrl Cortex N5 | 0.673 |  | 1.828 | 0.935 |
| KA Cortex N1 | 2.257 | 0.473 | 0.836 | 2.505 |
| KA Cortex N2 | 1.842 | 7.079 | 1.411 | 2.809 |
| KA Cortex N3 | 0.975 | 0.134 | 2.465 | 1.615 |
| KA Cortex N4 | 1.221 | 1.549 | 2.795 | 1.192 |
| KA Cortex N5 | 1.065 |  | 0.970 | 2.203 |
| Ctrl Hippo N1 | 0.816 | 0.410 | 0.924 | 1.014 |
| Ctrl Hippo N2 | 1.051 | 1.541 | 0.735 | 0.820 |
| Ctrl Hippo N3 | 0.838 | 1.261 | 1.565 | 1.220 |
| Ctrl Hippo N4 | 1.798 | 0.788 | 1.072 | 0.982 |
| Ctrl Hippo N5 | 0.496 |  | 0.705 | 0.963 |
| KA Hippo N1 | 7.434 | 0.413 | 0.317 | 1.167 |
| KA Hippo N2 | 1.052 | 0.926 | 0.932 | 0.841 |
| KA Hippo N3 | 5.271 | 0.882 | 1.379 | 1.147 |
| KA Hippo N4 | 1.319 | 1.526 | 1.071 | 1.123 |
| KA Hippo N5 | 6.074 |  | 0.610 | 1.044 |

| Suppl Fig. 5 D,G | Gapdh-, Ctrl-double norm Foxg1 mRNA |  |  |  |
| --- | --- | --- | --- | --- |
| group, structure, n° | 2h | 6h | 8h | 24h |
| Ctrl Cortex N1 | 0.492 | 1.066 | 1.258 | 1.114 |
| Ctrl Cortex N2 | 0.964 | 0.876 | 1.018 | 0.891 |
| Ctrl Cortex N3 | 1.242 | 1.196 | 0.745 | 1.072 |
| Ctrl Cortex N4 | 1.266 | 0.862 | 1.070 | 0.930 |
| Ctrl Cortex N5 | 1.036 |  | 0.909 | 0.993 |
| KA Cortex N1 | 1.214 | 0.938 | 0.900 | 1.289 |
| KA Cortex N2 | 1.449 | 3.458 | 1.197 | 1.103 |
| KA Cortex N3 | 1.128 | 0.960 | 0.871 | 0.986 |
| KA Cortex N4 | 1.424 | 2.133 | 1.475 | 0.944 |
| KA Cortex N5 | 1.101 |  | 0.938 | 3.640 |
| Ctrl Hippo N1 | 0.900 | 0.672 | 1.083 | 0.905 |
| Ctrl Hippo N2 | 1.429 | 1.017 | 1.195 | 1.056 |
| Ctrl Hippo N3 | 1.202 | 1.382 | 0.724 | 0.807 |
| Ctrl Hippo N4 | 0.898 | 0.929 | 1.037 | 1.117 |
| Ctrl Hippo N5 | 0.571 |  | 0.962 | 1.116 |
| KA Hippo N1 | 1.247 | 0.901 | 0.895 | 0.893 |
| KA Hippo N2 | 1.799 | 1.123 | 1.128 | 0.876 |
| KA Hippo N3 | 1.442 | 1.041 | 1.814 | 1.020 |
| KA Hippo N4 | 0.810 | 1.897 | 1.226 | 0.913 |
| KA Hippo N5 | 1.820 |  | 0.787 | 0.711 |

|  |  |
| --- | --- |
| # | 2.7 |
| subject: | KA assays |
| reference figures: | Supplementary Figure 5H-K |

| Suppl. Fig. 5 H-I | cFos<sup>-</sup> cell - norm Foxg1 protein |  |  |  |  |  |  |  |
| --- | --- | --- | --- | --- | --- | --- | --- | --- |
| group, structure, n° | 2h |  | 6h |  | 8h |  | 24h |  |
|  | cFos<sup>-</sup> cells | cFos<sup>+</sup> cells | cFos<sup>-</sup> cells | cFos<sup>+</sup> cells | cFos<sup>-</sup> cells | cFos<sup>+</sup> cells | cFos<sup>-</sup> cells | cFos<sup>+</sup> cells |
| KA Cortex N1 | 0.975 | 1.120 | 0.994 | 1.030 | 0.966 | 1.206 | 1.006 | 1.683 |
| KA Cortex N2 | 0.961 | 1.211 | 1.009 | 0.977 | 0.966 | 1.149 | 1.017 | 1.650 |
| KA Cortex N3 | 1.065 | 1.209 | 0.997 | 0.775 | 1.067 | 1.072 | 0.977 | 1.597 |
| KA Hippo N1 | 1.000 | 1.208 | 0.963 | 1.325 | 0.882 | 1.607 | 1.030 | 1.765 |
| KA Hippo N2 | 0.967 | 1.413 | 1.072 | 1.514 | 1.028 | 1.786 | 0.970 | 1.692 |
| KA Hippo N3 | 1.033 | 1.201 | 0.966 | 1.265 | 1.090 | 2.129 | 1.000 | 1.476 |
| KA DG N1 | 1.024 | 1.171 | 0.958 | 0.632 | 1.037 | 1.266 | 0.932 | 1.645 |
| KA DG N2 | 0.946 | 1.118 | 1.098 | 0.747 | 0.851 | 1.188 | 1.044 | 1.581 |
| KA DG N3 | 1.030 | 1.133 | 0.945 | 0.496 | 1.112 | 1.128 | 1.025 | 1.832 |

| Suppl. Fig. 5 K | cFos<sup>-</sup> cell - norm Foxg1 protein |  |
| --- | --- | --- |
| group, structure, n° | cFos<sup>-</sup> cells | cFos<sup>+</sup> cells |
| Ctrl Cortex N1 | 0.948 | 1.057 |
| Ctrl Cortex N2 | 0.882 | 0.950 |
| Ctrl Cortex N3 | 0.849 | 0.976 |
| Ctrl Cortex N4 | 1.049 | 1.154 |
| Ctrl Cortex N5 | 1.064 | 1.017 |
| Ctrl Cortex N6 | 1.095 | 0.933 |
| Ctrl Cortex N7 | 1.145 | 1.426 |
| Ctrl Cortex N8 | 1.183 | 1.510 |
| Ctrl Cortex N9 | 1.183 | 1.274 |
| Ctrl Cortex N10 | 0.871 | 1.207 |
| Ctrl Cortex N11 | 0.863 | 1.173 |
| Ctrl Cortex N12 | 0.869 | 1.327 |
| Ctrl Hippo N1 | 1.050 | 1.267 |
| Ctrl Hippo N2 | 1.023 | 1.345 |
| Ctrl Hippo N3 | 0.902 | 1.256 |
| Ctrl Hippo N4 | 1.086 | 1.148 |
| Ctrl Hippo N5 | 1.097 | 1.053 |
| Ctrl Hippo N6 | 1.016 | 1.150 |
| Ctrl Hippo N7 | 1.041 | 1.477 |
| Ctrl Hippo N8 | 1.121 | 1.279 |
| Ctrl Hippo N9 | 1.279 | 1.503 |
| Ctrl Hippo N10 | 0.802 | 1.208 |
| Ctrl Hippo N11 | 0.842 | 1.360 |
| Ctrl Hippo N12 | 0.741 | 1.537 |
| Ctrl DG N1 | 0.917 | 1.061 |
| Ctrl DG N2 | 0.921 | 1.215 |
| Ctrl DG N3 | 0.896 | 0.969 |
| Ctrl DG N4 | 0.918 | 0.929 |
| Ctrl DG N5 | 0.913 | 0.717 |
| Ctrl DG N6 | 0.893 | 1.030 |
| Ctrl DG N7 | 1.110 | 1.355 |
| Ctrl DG N8 | 1.466 | 2.037 |
| Ctrl DG N9 | 1.279 | 2.093 |
| Ctrl DG N10 | 0.920 | 1.027 |
| Ctrl DG N11 | 0.881 | 1.427 |
| Ctrl DG N12 | 0.886 | 1.417 |

